# Uncovering novel roles of miR-122 in the pathophysiology of the liver: Potential interaction with NRF1 and E2F4 signaling

**DOI:** 10.1101/2023.06.30.547158

**Authors:** Martha Paluschinski, Jessica Schira-Heinen, Rossella Pellegrino, Lara R. Heij, Jan Bednarsch, Ulf P. Neumann, Thomas Longerich, Kai Stuehler, Tom Luedde, Mirco Castoldi

## Abstract

MicroRNA miR-122 plays a pivotal role in liver function. Despite numerous studies investigating this miRNA, the global network of genes regulated by miR-122 and its contribution to the underlying pathophysiological mechanisms remain largely unknown. To gain a deeper understanding of miR-122 activity, we employed two complementary approaches. Firstly, through transcriptome analysis of polyribosome-bound RNAs, we discovered that miR-122 exhibits potential antagonistic effects on specific transcription factors known to be dysregulated in liver disease, including nuclear respiratory factor-1 (NRF1) and the E2F Transcription Factor 4 (E2F4). Secondly, through proteome analysis of hepatoma cell transfected with either miR-122 mimic or antagomiR we discovered changes in several proteins associated with increased malignancy. Interestingly, many of these proteins were reported to be transcriptionally regulated by NRF1 and E2F4, six of which we validated as miR-122 targets. Among these, a negative correlation was observed between miR-122 and glucose-6-phosphate dehydrogenase levels in the livers of patients with hepatitis B virus-associated hepatocellular carcinoma. This study provides novel insights into potential alterations of molecular pathway occurring at the early stages of liver disease, driven by the dysregulation of miR-122 and its associated genes.

## Introduction

The liver plays a critical role in maintaining metabolic homeostasis by synthesizing, storing, and redistributing metabolites. It also acts as the body’s primary organ for detoxification, making it susceptible to toxicity. Acute or repeated insults to the liver are known risk factors for liver disease [1]. MicroRNAs (miRNAs) are small noncoding RNAs that regulate gene expression at the post-transcriptional level. Extensive evidence suggests that miRNAs have the ability to control networks comprising hundreds of genes, thus triggering pleiotropic effects in response to physiological and pathological changes in their expression. Among these miRNAs, miR-122 has been extensively studied and implicated in various aspects of hepatic metabolism, such as cholesterol metabolism [2] and iron homeostasis [3]. Additionally, miR-122 was shown to possess tumor suppressor and anti-inflammatory properties [4]. Notably, miR-122 was found significantly downregulated in the livers of approximately 70% of hepatocellular carcinoma (HCC) patients [5] and lower levels of miR-122 have been associated with shorter recurrence times in resected HCC patients [6]. Studies on miR-122 knockout (KO) animals have revealed that while these animals are viable, their livers exhibit infiltration by inflammatory cells, leading to the development of steatohepatitis, fibrosis, and HCC [4], while miR-122 inhibition in human liver organoid has been shown to lead to inflammation [7]. Collectively, these findings support the existence of a link between miR-122, inflammation, and liver cancer. Despite the extensive knowledge on miR-122, numerous questions remain unanswered. For instance, the mechanisms underlying miR-122 downregulation during the onset of liver diseases are still poorly understood. Is the downregulation of miR-122 merely a consequence of healthy parenchyma loss, or is it a physiological process with a specific regulatory function that has gone awry? Furthermore, what are the molecular networks that underlie its functional role? To further explore the impact of miR-122 on liver pathophysiology, a combination of transcriptomic and proteomic approaches was applied, providing evidence that alterations in miR-122 levels have the potential to modify the translational activity of thousands of genes. Our data suggest that miR-122 potentially modulate genes downstream of several transcription factors (TFs), including nuclear respiratory factor-1 (NRF1), transcription factor E2F4 (E2F4), and Yin-Yang 1 (YY1). Through our investigation, several novel miR-122 targets were identified and validated. Notably, elevated expression of these genes has been linked to an unfavorable clinical outcome in HCC patients, suggesting their potential prognostic value. Overall, this study sheds new light on the potential changes in signaling pathways that may occur during the onset of liver disease, which may be potentially linked to alterations in miR-122 and its associated mRNA targets.

## Results

### Preparation and isolation of polyribosomes from sucrose gradient

Despite the extensive literature on miRNA biology, the identification of miRNA targets remains a contentious issue. It was initially proposed that miRNAs inhibited protein synthesis via inducing degradation of their mRNA targets [8,9]. On the contrary, studies on isolated polyribosomes suggested that miRNAs primarily cause mRNA destabilization [10,11]. Consequently, analyzing polyribosomes could facilitate the identification of *bona fide* miRNA targets and their associated regulatory network (reviewed in; [12]). To evaluate the efficacy of polyribosome analysis in identifying genes responsive to miR-122, human hepatoma cell lines (Huh-7) were transfected with miR-122 mimics or inhibitors (antagomiRs). Polyribosomes were then isolated through fractionation on sucrose gradients. As a proof of principle, the relative distribution of genes of interest (GOI) on the polyribosomes was assessed using quantitative real-time PCR (Figure 1). We found that miR-122 overexpression led to a significant increase in the levels of miR-122 in the polyribosomal fractions (Figure 1B). In contrast, the expression and distribution of other miRNAs, here illustrated by miR-192, remained unaffected (Figure 1C). Consistently, the analysis of the best-known direct target of miR-122, SLC7A1 (high affinity cationic amino acid transporter 1; [13]), identified a significant increase in the abundance of SLC7A1 RNA in the polyribosomal fractions isolated from antagomiR transfected cells (Figure 1D). Notably also the distribution on polyribosimal fractions of Hemochromatosis (HFE), Hemojuvelin (HJV) and Hepcidin (HAMP) transcripts, all genes that were previously reported to respond to miR-122 [3], were found to be affected by the modulation of miR-122 levels (**Supplementary Figure S1**). Conversely, the distribution of genes that are not expected to respond to miR-122 including Transferrin-receptor 2 (TFR2) Bone morphogenetic protein 6 (BMP6), L-Ferritin and of the Growth differentiation factor 15 (GDF15; Figure 1E and **Supplementary Figure S1**), were not significantly affected. Thus, these results are supporting the conclusion that known miR-122 responsive genes can be identified by analyzing the distribution of genes of interest (GOIs) on polyribosomes.

**Figure 1:**
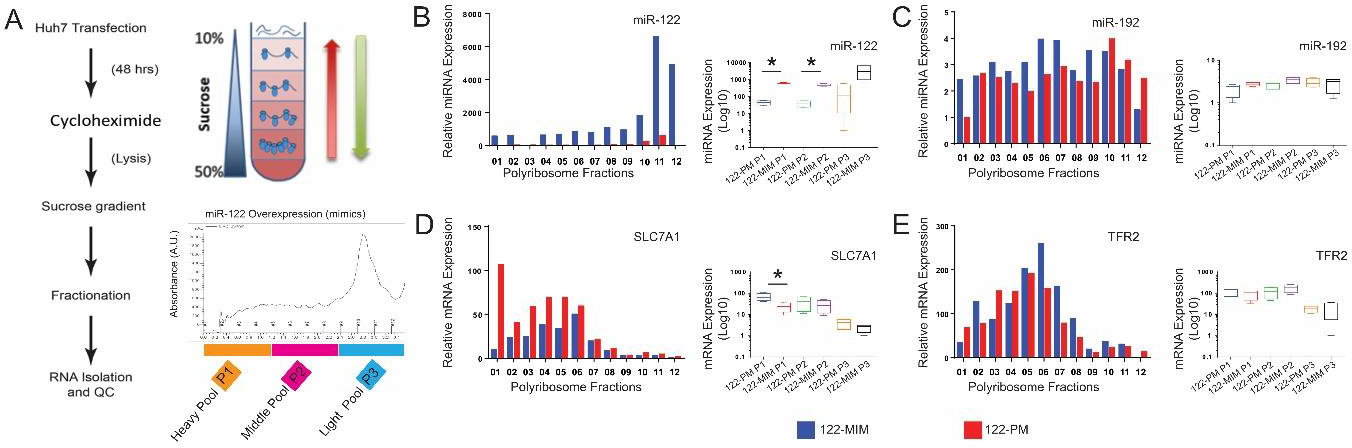
Direct identification of miR-122 targets by polyribosomal profiling on sucrose gradient. **(A)** Huh-7 cells were transfected with either miR-122 mimic (122-MIM) or miR-122 inhibitor (122-PM) and lysed 48 h after transfection. The cytosolic extracts were sedimented by ultracentrifugation on a 10-50% sucrose gradient. Upon fractionation, RNA isolation, quantification and quality control (QC) were performed for the corresponding polyribosomal fractions. UV absorbance profile of the nucleic acids associated to polyribosomes was measured at 254 nm from miR-122 overexpressing and miR-122 depleted cells. The collected fractions are depicted as (A1-A12). Analysis of miRNA and mRNA distribution in isolated polyribosomes of cells transfected with miR-122 mimics (blue) or cells treated with miR-122-antagomiR (red) by qPCR. **(B)** The analysis showed an increased level of miR-122 associated to polyribosomes in cells treated with miR-122 mimics **(C)**, while distribution of unrelated miRNAs as shown for miR-194 was unaffected by the transfection. **(D)** In polyribosomes isolated from antagomiRs treated cells, a significant enrichment towards the heavy transcribed fraction in the distribution of SLC7A1 mRNA, the best characterized target for miR-122 was observed. **(E)** In contrast, the transferrin receptor 2 (TFR2) mRNA, which is not a miR-122 target, did not show any significant changes in its polyribosomal distribution. Data are shown either as miRNA/mRNA expressions in individual fractions or as box plot using mean ± SD for the pooled fractions (n = 4). Statistical analysis was performed with two-tailed T-test, p ≤ 0.05 was considered significant. *, p ≤ 0.05.

### Modulation of miR-122 expression results in changes in polyribosome occupancy for thousands of transcripts

To better comprehend the impact of miR-122 on the cellular transcriptome and its molecular networks, we took a comprehensive approach. RNAs were isolated from the individual polyribosomal fractions of cells transfected with either miR-122 mimics (122-MIM) or antagomiRs (122-PM). These RNAs were then combined to create polyribosomal pools of decreasing densities [i.e., P1 = Heavy (fractions A1-A4), P2 = Middle (fractions A5-A8), and P3 = Light (fractions A9-A12); visualized in Figure 2A]. To conduct an in-depth analysis, these six different

**Figure 2:**
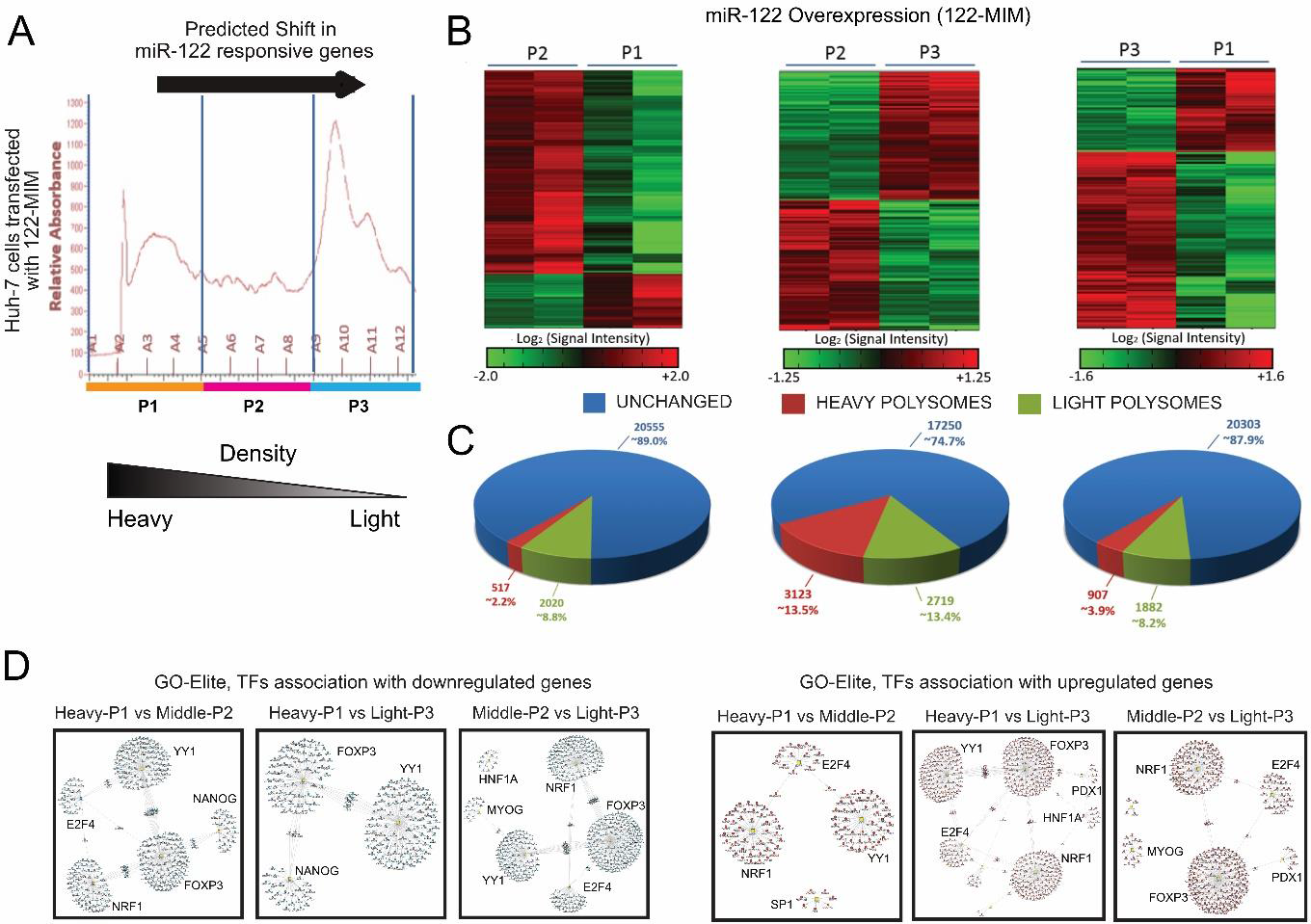
Changes in mRNA distribution on polyribosomes in response to miR-122 overexpression (122-MIM) Huh-7 cells were treated with miR-122 mimic and lysed 48 h later. Following sedimentation on 10-50% sucrose gradient, lysates were fractionated and fractions were pooled as indicated **(A)** to create a heavy pool (P1), a middle pool (P2), and a light pool (P3). RNA isolated from each individual pool were hybridized to Affymetrix arrays and data were analyzed by AltAnalyze [14] using a cutoff of 1.5 fold changes (significance level p ≤ 0.05, one-way ANOVA) The arrow indicates the expected target gene shift across polyribosomes in response to miR-122 overexpression. Microarray data are available via the GEO repository under the accession number **GSE234690**. **(B)** Hierarchical clustering representing the differential distribution of mRNAs associated to the polyribosomal pools (P2 *vs.* P1, P2 *vs.* P3, and P3 *vs.* P1). Cousine Matrix was applied to generate hierarchical tree of gene clusters (n = 2). **(C)** Cake chart illustrating the number of regulated transcripts between the different pools P2 *vs.* P1 (left), P2 *vs.* P3 (middle), and P3 *vs.* P1 (right). **(D)** Evaluation of microarray data by GO-Elite algorithm identified a link between miR-122 responsive genes and the nuclear respiratory factor 1 (NRF1), and E2F transcription factor 4 (E2F4), transcription Forkhead box protein P3 (FOXP3), Yin Yang 1 (YY1).

RNA pools (i.e., three densities for two conditions for two different experiments) were hybridized to microarrays, and bioinformatics analysis was carried out to identify miR-122 responsive genes. Specifically, AltAnalyze [14] was used to compare the changes in the relative amount of genes across the three different pools (i.e., P1 vs P2, P2 vs P3, and P1 vs P3) within each condition (i.e., 122-MIM or 122-PM). Moreover, comparisons were carried out across the conditions (i.e., 122-MIM vs 122-PM) within each pool (i.e., P1, P2 or P3) for a total of nine comparisons. In detail, we measured: i) the significant shift of genes from high-density to low-density fractions in cells transfected with the miR-122 mimic (122-MIM; Figure 2B), which led to the shift of 6621 transcripts towards lighter density polyribosomes indicating a reduced translational turnover of the associated genes in response to miR-122 upregulation. ii) The significant shift of genes from low-density to high-density fractions in cells transfected with the antagomiR (122-PM; **Supplementary Figure S2**), which identified the shifting of 6163 transcripts towards polyribosomes with heavier density indicating an increase in translational turnover of these genes in response to miR-122 inhibition. iii) The significant change in the expression of genes across the same fractions in 122-MIM versus 122-PM (**Supplementary Figure S3**), which identified the significant enrichment of 1028 transcripts in polyribosomes isolated from 122-PM transfected cells.

### Potential regulatory role of miR-122 on NRF1, E2F4 and YY1 mediated gene regulation

To explore the functional significance associated with the shift in mRNA occupancy in response to changes in miR-122, the GO-Elite algorithm [15] was used to analyze the differentially regulated genes (DEGs) from transcriptomic data. GO-Elite allows the identification of common biological ontology pathways that describe a specific set of genes. Specifically, GO-Elite identified a significant enrichment for genes associated to the transcription factors (TFs) YY1, E2F4, NRF1 and Forkhead box P3 (FOXP3; Figure 2D and **Supplementary Table S1**). Markedly, these transcription factors have been found to be dysregulated in patients with liver diseases including steatosis, nonalcoholic steatosis and HCC [16–19]. Interestingly, the mining of sequencing data of cohort of patients with liver hepatocellular carcinoma (LIHC) included in The Cancer Genome Atlas (TCGA) shows that YY1, E2F4 and NRF1 expressions were significantly upregulated in the liver of HCC patients (**Supplementary Figure S4**). Moreover, the Kaplan-Meier curves from these data shows that LIHC patients with lower YY1, E2F4 and NRF1 expression have a significantly higher probability of survival (**Supplementary Figure S4**). Contrariwise, FOXP3 (Figure 2D), which is mainly expressed in Regulatory T-cells (Treg; [17]), was found to be significantly downregulated in TCGA LIHC-cohort and that LIHC patients with higher FOXP3 expression had a significantly higher probability of survival (**Supplementary Figure S5**). The upregulation of YY1, E2F4 and NRF1 in the liver of HCC patients was also confirmed through the analysis of cohorts of patients downloaded from GEO (i.e., GSE62232 and GSE6764; **Supplementary Figure S6**). Significantly, the application of GO-Elite to the analysis of patients’ cohorts with grade III HCC (accession number GSE45050; [20]) identified a significant enrichment for genes associated to YY1, NRF1 and FOXP3 transcription factors (**Supplementary Figure S7A-S7B**), indicating that modulations of similar changes might be extrapolated through the analysis of the whole liver. Remarkably, a similar enrichment for YY1, NRF1, E2F4 and FOXP3 TFs was also identified by GO-Elite in both the livers of miR-122 KO animals (GSE31453) and in mouse model for liver cancer (GSE27713; **Supplementary Figure S7C)**.

### miR 122-responsive genes are under NRF1, E2F4 and YY1 transcriptional control

To validate the microarray and GO-Elite data, we followed a rigorous selection process resulting in the selection of 43 genes based on the following criteria. Firstly, a sub-list of genes was generated by selecting TFs-associated genes [**Supplementary Data S2,** list for individual TFs are found in **Supplementary Tables S2 (E2F4), S3 (FOXP3), S4 (NRF1) and S5 (YY1)]** responsive to miR-122 and present in at least two polyribosomal pool comparisons. Secondly, to identify potential miR-122 targets, this sub-list was overlaid with a list of miR-122 predicted targets generated by *miRWalk* [21]. Subsequently, genes that exhibited less than a 1.5-fold differential expression were filtered out. Finally, the remaining genes were thoroughly reviewed in the literature, specifically focusing on their links to "liver disease," "infection," "inflammation," or "cancer" generating a final list of 45 genes (**Supplementary Table S6**). Next, qPCR was used to assess the expression of the target genes in Huh-7 cells following transfection with 122-MIM (Figure 3), resulting in the significant reduction in the expression of 24 genes (i.e., ∼56% of the selected genes). Specifically, the genes affected after 24 hours were BAG1, BAX, CCDC43, CD47, DIXDC1, DSG2, F2RL2, G3BP2, KPNA6, KPNB1, NUP210, P4HA1, PCDC4, PDCD2, and TBC1D22B. After 48 hours, the genes affected were CMTM7, HCFC1, KIF1B, KIF3A, MINK1, PDCD2, RNF26, SPRED2, and TNPO1 (Figure 3B and 3C). It is important to note that the identification of ’unresponsive’ genes does not exclude them from potentially being targeted by miR-122, as their regulation could occur at the translational level. Furthermore, we made an intriguing observation that the overexpression of miR-122 led to a significant reduction in E2F4 and NRF1 levels, while the expression of YY1 remained unaffected (Figure 3B). Notably, GO-Elite enrichment analysis also identified a significant enrichment of genes associated with the FOXP3 transcription factor (Figure 2D, **Supplementary Tables S1**). Although FOXP3 mRNA was not detected in Huh-7 cells our analysis suggested that Huh-7 cells expressed a subset of FOXP3-responsive genes that also respondeds to changes in miR-122 levels (**Supplementary Table S6**). Overall, these findings highlight that alterations in miR-122 levels can influence the ribosomal occupancy of numerous transcripts, which have been linked to chronic liver diseases and liver cancer. Consequently, we propose that miR-122 may directly or indirectly contribute to modifying the translational turnover of gene subsets linked to the transcription factors E2F4, NRF1, and YY1, as well as their associated molecular networks.

**Figure 3:**
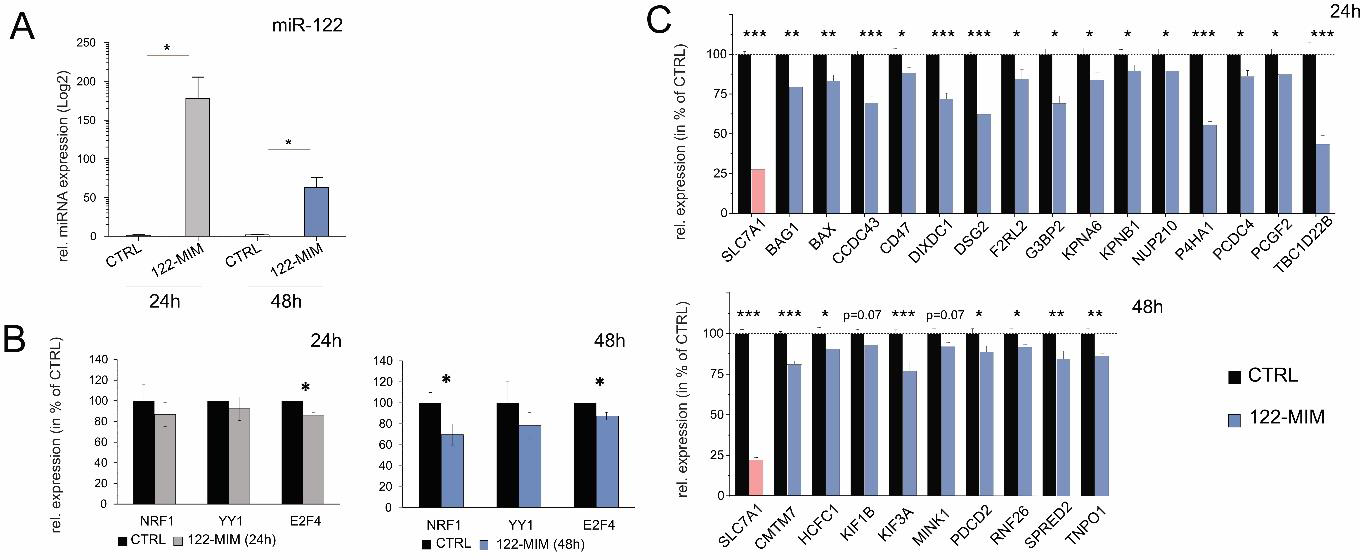
qPCR validation of potential miR-122 target genes in miR-122 overexpressing Huh-7 cells. Human hepatoma cells Huh-7 were transfected with miR-122 mimic and RNA isolated 24 h and 48 h after transfection. **(A)** The overexpression of cellular miR-122 was measured 24 h and 48 h after transfection by miQPCR [46]. Relative expression changes for selected target gene candidates are depicted for 24 h **(B)** and 48 h **(C)** post transfection. Death effector domain-containing protein (DEDD) mRNA was identified as the most stable transcript in the data set using *GeNorm* algorithm for reference gene quality. While the validated miR-122 target SLC7A1 mRNA served as positive control. The values of control cells were set to 100%. Data are shown as mean ± SD (n = 5). Statistical analyses were performed with two-tailed T-test, p ≤ 0.05 was considered significant. ns = Not significant; * p ≤ 0.05; ** p ≤ 0.01; *** p ≤ 0.001. Full gene names are listed in the **Supplementary Data S2.**

### Proteomic analysis uncovers proteins linked to liver diseases and reveals miR-122 potential role in regulating energy metabolism and EVs secretion

Further investigations were carried out to evaluate the impact of changes in miR-122 expression on protein levels. Huh-7 cells were transfected with either miR-122 mimics (122-MIM) or antagomiRs (122-PM). The proteins extracted from these cells were then analyzed by mass spectrometry (Figure 4). Among the 1940 identified proteins, 504 proteins exhibited significant regulation (p ≤ 0.05; Figure 4A, 4B and **Supplementary Table S7**). To gain insight into the functional implications of these changes, ShinyGO [22] was used to perform enrichment analysis. The results, shown in Figure 4C, shed some light into the potential effects of miR-122 alterations on the cellular proteome. Notably, an examination of the GO terms related to "Biological Processes" suggest that changes in miR-122 expression might affect energy metabolism (Figure 4C, **top panel**). Additionally, an analysis of the GO terms related to "Cellular Components" identified a significant reduction in terms associated with “extracellular vesicles”, while a significant enrichment of terms associated with mitochondria and organelle was detected (Figure 4C, **lower panel**). These findings support existing evidence suggesting the involvement of miR-122 in regulating both mitochondrial [23] and energy metabolism [24].

**Figure 4:**
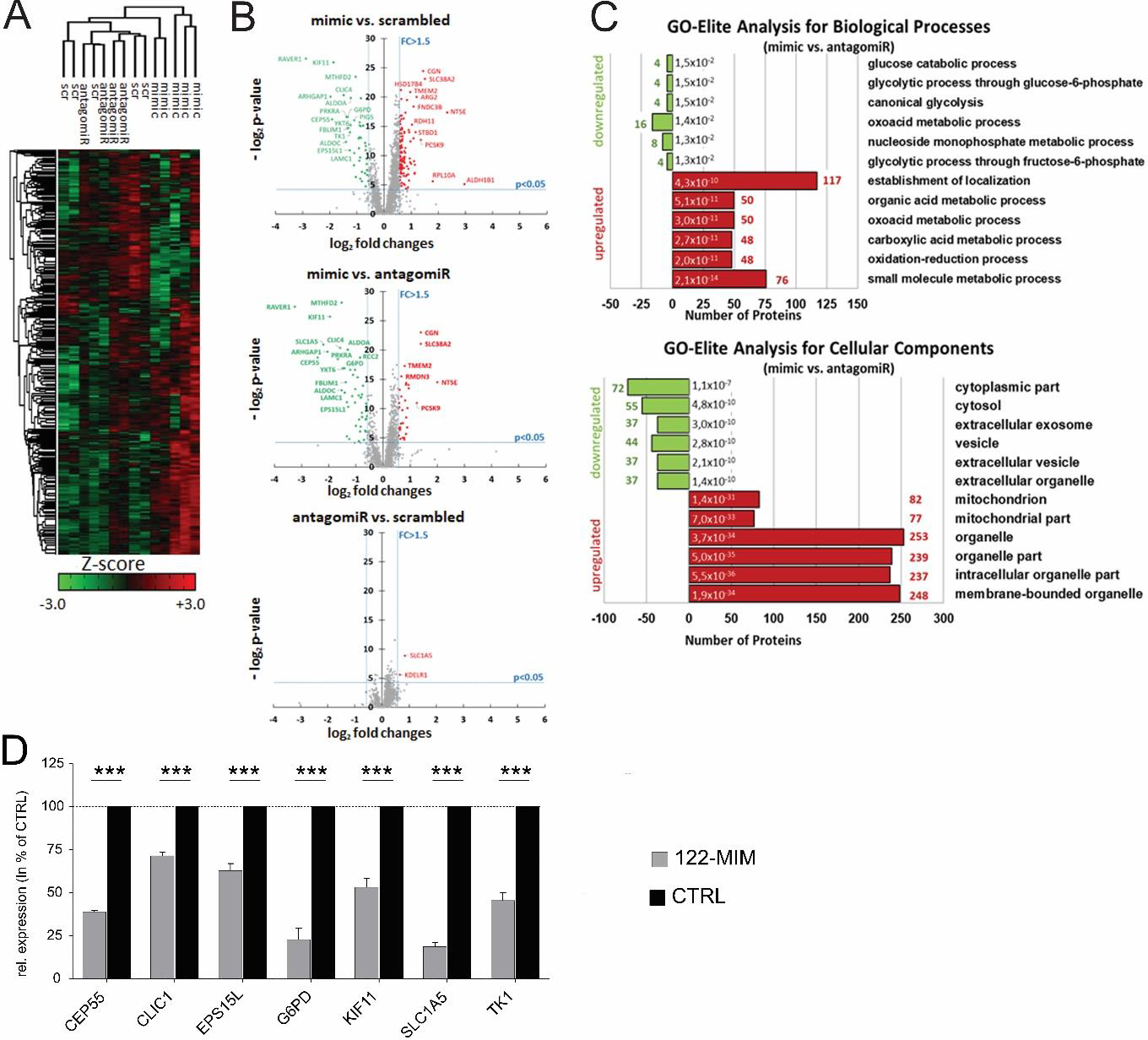
Proteome analysis of miR-122 enriched and depleted Huh-7 cells. **(A)** Quantitative proteome analysis of Huh-7 cell lines transfected with miR-122 mimics, antagomiRs or scrambled oligo control (scr). Unsupervised hierarchical clustering illustrating the relative abundance (Z-Score) of all quantified proteins in Huh-7 upon modulation of cellular miR-122 levels (n = 5). **(B)** Volcano blot illustrating the pairwise differences in protein abundances in Huh-7 treated with miR-122 mimic, antagomiR and scrambled oligo transfected cells. Expression changes between groups are plotted in log_2_ scale on the abscissa (x-axis), while p-values measured by one-way ANOVA were plotted in – log_2_ scale on the ordinate (y-axis). Thresholds for fold changes (FC ≥ 1.5 or FC ≤ −1.5) are shown as perpendicular dashed lines, while a threshold for statistical relevance (p ≤ 0.05) is shown as horizontal dashed line. Proteins highlighted in green were found in significant lower abundance, while significantly more abundant proteins are highlighted in red. **(C)** Functional analysis of regulated proteins was conducted using Gene Ontology enrichment tool GOrilla [52]. The schematic representation illustrates the number of proteins associated with the 6 most significantly (i.e. with lowest p-value) depleted (green) and enriched (red) GO terms regarding biological process (upper) and cellular components (lower). Number of proteins related to the GO terms is given in red (enriched) or green (depleted) next to the bars, while p-value for each GO term is written next to the y-axis. **(D)** Relative expression changes for selected target proteins at mRNA levels in response to miR-122 overexpression (122-MIM). The values of scrambled control (CTRL) transfected cells were set to 100%. Data are shown as mean ± SD (n = 5). Statistical analyses were performed with two-tailed T-test, p ≤ 0.05 was considered significant. *** p ≤ 0.001.

### miR-122 regulates the expression of proteins potentially contributing to liver cancer development

To validate the proteomic data, a rigorous selection process was followed. First, a list of proteins that showed significant responsiveness to changes in miR-122 levels was compiled. From this list, we selected 43 proteins that exhibited more than 1.5-fold change and were predicted as potential miR-122 targets by the *miRWalk* target prediction database. To examine the connections between transcriptomic and proteomic data, transcription factors enrichment analysis (TFEA) was performed. For this purpose, ShinyGO was used to cross-references the list of 43 miR-122 responsive proteins with the ENCODE-motifs database [25]. The TFEA results revealed that approximately 90% (40 out of 43) of the genes contained binding motifs for E2F4, and around 86% (37 out of 43) contained binding motifs for NRF1 (**Supplementary Table S8**). Next, a literature search was conducted to focus on proteins specifically associated with liver disease in general, which selected 20 proteins (**Supplementary Table S9** and **Supplementary Data S2A**), seven of which (i.e., CEP55, CLIC1, EPS15L1, G6PD, KIF11, SLC1A5, and TK1) were found to be specifically associated with hepatocellular carcinoma. This conclusion was supported not only by literature mining but also through the analysis of publicly available databases, including GEO (i.e., accession numbers GSE62232 and GSE6764, **Supplementary Figure S8**) and TCGA (**Supplementary Figure S9**). To this end, detailed analysis of the sequencing data from the TCGA LIHC cohort showed that CEP55, CLIC1, EPS15L1, G6PD, KIF11, SLC1A5, and TK1 were significantly upregulated in the livers of HCC patients. Notably, the Kaplan-Meier curve indicated that patients with lower expression of these genes had a significantly higher probability of survival. Next, the regulatory effect of miR-122 on the mRNAs encoding these proteins was evaluated. Here we show that a significant decrease in the mRNA levels of these genes was measured in 122-MIM transfected cells (Figure 4D), supporting the conclusion that miR-122 affects the stability of these transcripts, either directly or indirectly.

### CEP55, CLIC1, G6PD, KIF11, SLC1A5 and TK1 but not EPS15L1 are direct target of miR-122

To investigate the influence of miR-122 on the stability of specific transcripts, the full length 3’UTRs of CEP55, CLIC1, EPS15L1, G6PD, KIF11, SLC1A5, and TK1 were cloned in the pMIR(+) and pMIR(-) Luciferase reporter vectors, and co-transfected with miR-122 mimics into HEK293 cells (Figure 5). Here we show that in cells transfected with miR-122 mimics, the Luciferase activity of the pMIR(+) vectors (i.e., carrying the 3’UTRs in their native orientation) was significantly reduced. This effect was observed for all 3’UTRs, except for EPS15L1. Conversely, miR-122 overexpression had no impact on the Luciferase activity of the pMIR(-) control vectors, which contained the 3’UTRs cloned in the reverse complement orientation. Consequently, it can be inferred that miR-122 directly regulates the expression of CEP55, CLIC1, G6PD, KIF11, SLC1A5, and TK1. However, in the case of EPS15L1, the regulation by miR-122 is likely indirect, as demonstrated by a noticeable decrease in EPS15L1 protein levels observed in Western blot analysis (Figure 5B) upon miR-122 upregulation.

**Figure 5:**
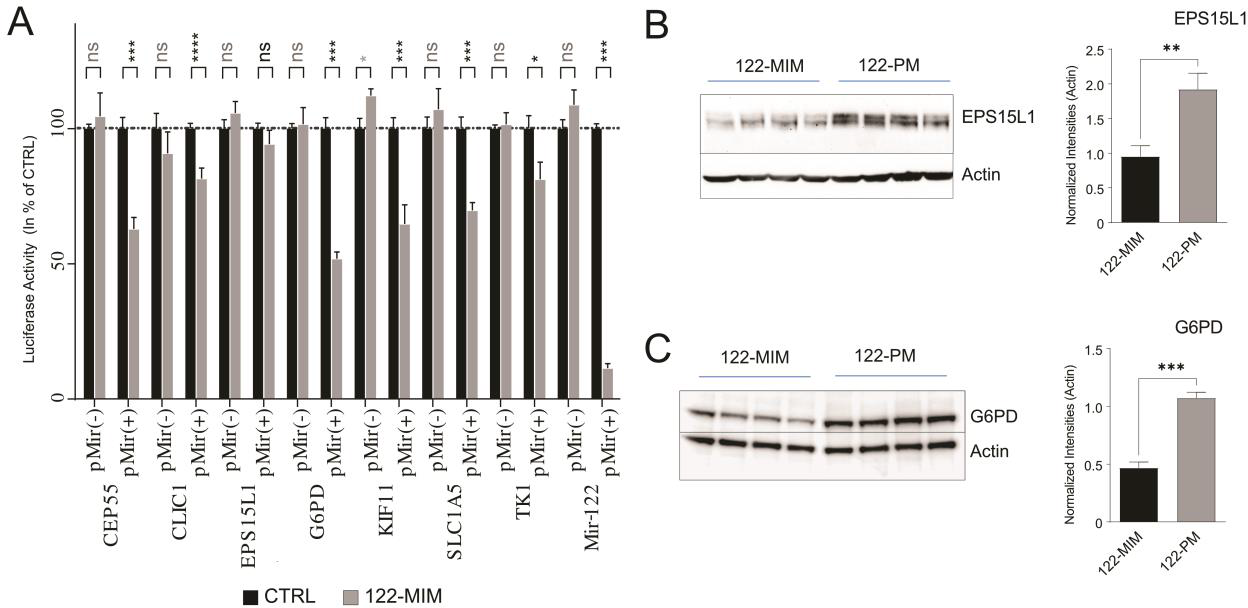
Correlation between miR-122 and G6PD expression in tumor tissue of HCC patients with or without HBV infection. **(A)** The full length of 3’UTR for the 7 GOIS were cloned into luciferase promoter vector in the sense orientation [pMir(+)] or in the antisense orientation [pMir(-)] as negative control. The plasmids pMir(+)_122 containing the perfect miR-122 binding site in sense orientation, and pMir(-)_122 (miR-122 binding site in the antisense orientation) served as positive and negative control, respectively. Plasmids were transfected in presence (grey) or absence (black) of miR-122 mimic into HEK293 cells and luciferase activity was measured 24 h post transfection. Relative luciferase activities are shown as percentage of plasmid transfection control (n = 4). Western Blot analysis demonstrated differential **(B)** EPS15L1 and **(C)** G6PD expression in Huh-7 treated with miR-122 mimic (122-MIM) or miR-122 inhibitor (122-PM) for 48 h (n = 4). Statistical analyses were performed with two-tailed T-test, p ≤ 0.05 was considered significant. ns = Not significant; * p ≤ 0.05; ** p ≤ 0.01; *** p ≤ 0.001.

### The expression of G6PD and miR-122 is inversely correlated in the liver of HBV patients with HCC

Our data indicates that miR-122 directly controls the translation of genes that may contribute to HCC development in human (Figure 5A). In order to delve deeper into these findings and establish a connection to human hepatocarcinogenesis, the focus was placed on G6PD. Consistent with previous research [26], our study confirms that miR-122 interacts with G6PD 3’UTR to regulates its expression both at the mRNA (Figure 4D) and at the protein level (Figure 5C). Analysis of G6PD expression in the TCGA cohorts of LIHC indicates that G6PD serves as a prognostic marker for HCC, with higher expression associated with a worse prognosis (Figure 7A). Notably, analysis of miR-122 expression in TCGA LIHC cohorts indicated the inverse correlation between G6PD and miR-122 in liver cancer patients (**Supplementary Figure S9**). Multiple studies have reported an elevation in G6PD levels in various types of human cancer, including HCC [27–29]. To this end, G6PD protein levels were measured by Western blot in the liver tissue of HCC patients and found to be significantly upregulated (Figure 6, patient description can be found in the **Supplementary Data S2B**).

**Figure 6:**
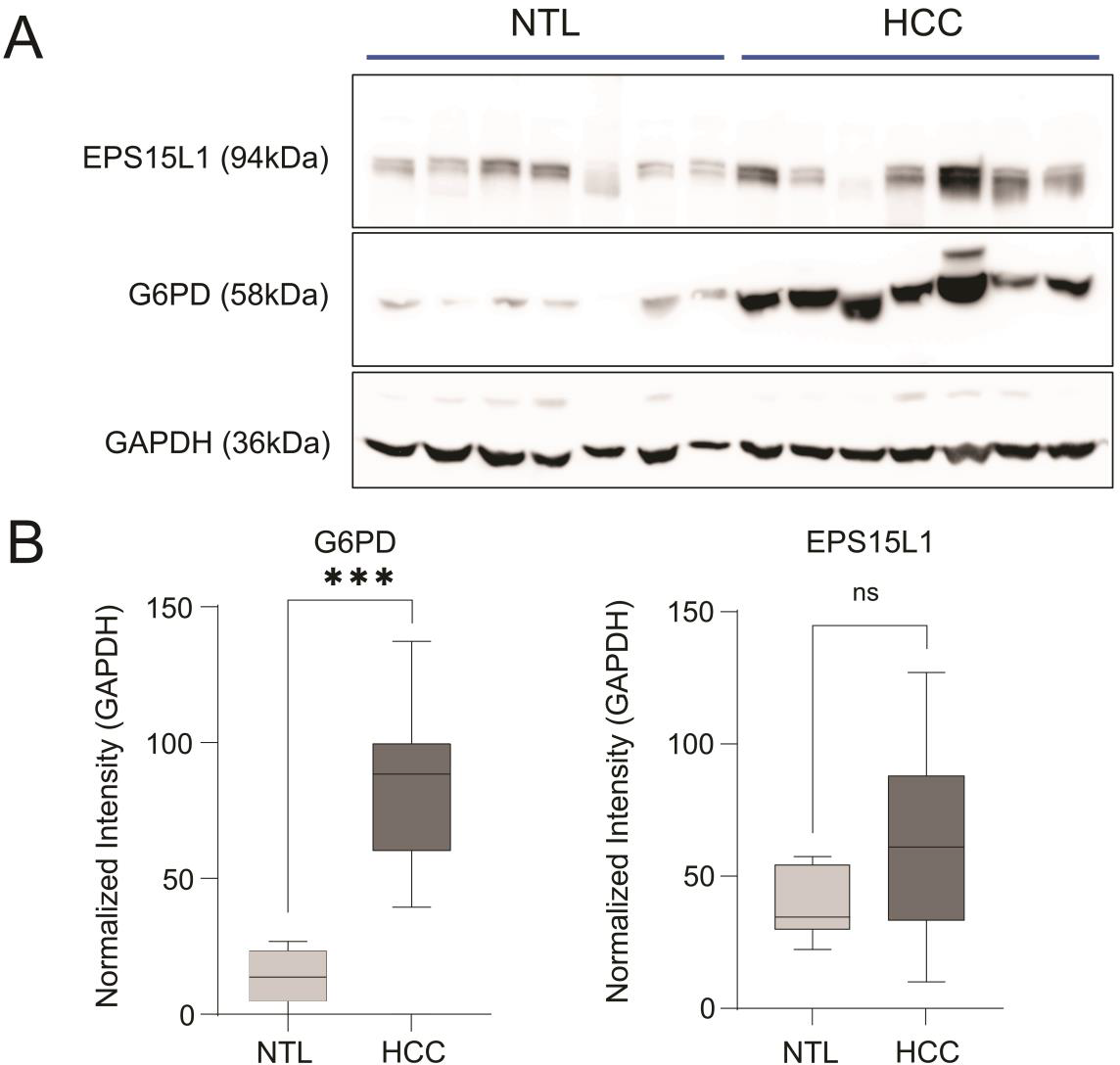
G6PD protein levels are significantly increased in the tumor tissue isolated from the livers of HCC patients. **(A)** Total proteins were isolated from the livers tissue of HCC patients, and the Western blot analysis for EPSL15L1 and G6PD was carried out in the Tumor (HCC, n = 7) and in the non-tumor (NTL, n = 7). **(B)** Quantification of protein expression normalized to GAPDH. Statistical analyses were performed with two-tailed T-test, p ≤ 0.05 was considered significant. ns = Not significant; *** p ≤ 0.001.

**Figure 7:**
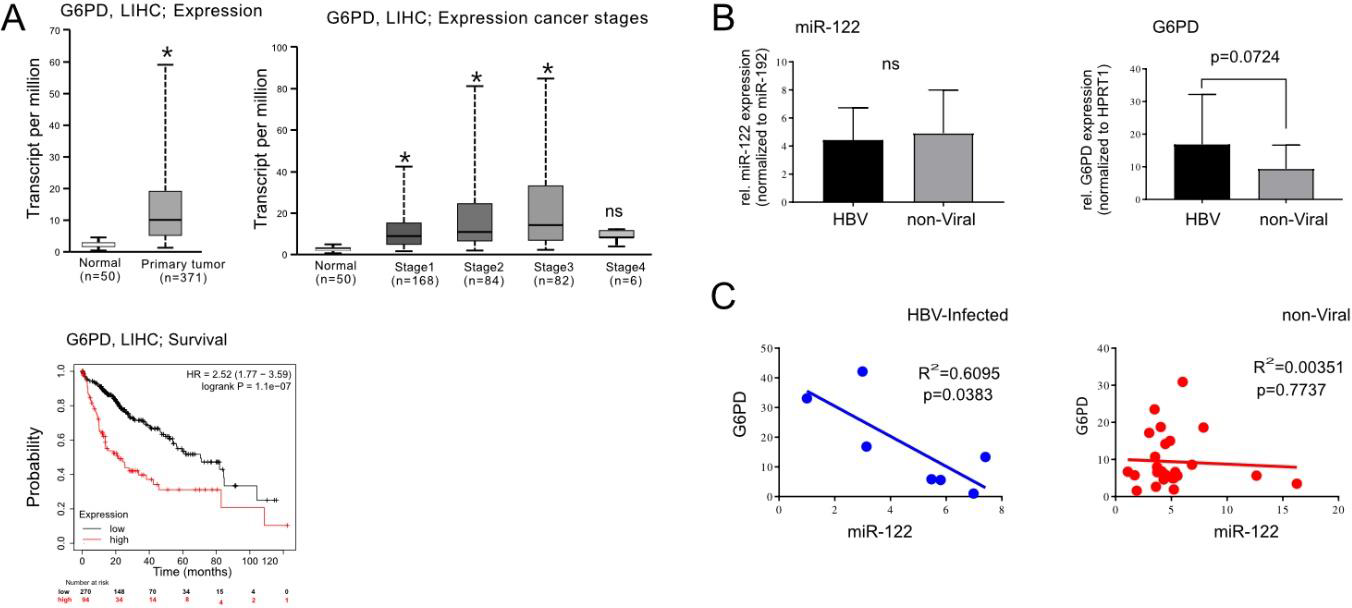
Correlation between miR-122 and G6PD expression in tumor tissue of HCC patients with or without HBV infection. **(A)** Analysis of LIHC cohorts from TCGA. Expression of G6PD was significantly increased in normal vs primary tumor tissue (Left panel) as well as in the livers of patients with cancer Stage 1 to 3 (Right panel). **(B)** Total RNAs were isolated from FFPE liver tissue of HCC patients with (HBV-Infected; n = 7) or without (non-viral: n = 26) HBV infection. Expression profiling of miR-122 (Left panel) and G6PD (Right panel) were measured by using qPCR. **(C)** Linear regression analysis showed a significant (p = 0.0383) inverse correlation between miR-122 and G6PD in the livers of HBV-associated HCC (HBV-HCC) patients, whereas no correlation was found in non-viral HCC patients. Two-tailed T-test was used to compare two groups, One-Way ANOVA was used to compare three or more groups, whereas liner regression (R^2^) was used to assess correlation between samples. p ≤ 0.05 was considered significant. ns = Not significant; * p ≤ 0.05.

In order to investigate further the link between miR-122, G6PD in liver cancer development, we specifically focused on HCC patients with or without hepatitis B virus (HBV)-infection. Independent studies have shown that while HBV infection enhances G6PD expression through hepatitis B viral protein X (HBx)-mediated Nrf2 activation [30], miR-122 was downregulated in the livers of HBV-infected patients [31]. To assess the relationship between miR-122 and G6PD in the livers of HBV-infected patients, RNA samples were extracted from FFPE tissue slices of patients with HCC with HBV (n = 7) or without HBV infection (n = 21; patient description can be found in the **Supplementary Data S2B**) and analyzed by qPCR. Although no differences were observed in the overall expression of miR-122 between the two groups (Figure 7B), a trend toward increased G6PD expression (p = 0.0724) was observed in the tumor tissue of HBV-infected patients. Importantly, when linear regression analysis was applied to the datasets, an inverse correlation between miR-122 and G6PD was observed in the HBV-infected HCC group (R2 = 0.6095, p = 0.0383; Figure 7C), but not in the non-viral HCC group. Based on these findings we propose that the HBV-mediated suppression of miR-122 may directly contribute to the upregulation of G6PD observed in the livers of patients with chronic HBV infection.

## Discussion

The dysregulation of miR-122 has been implicated in the development and progression of liver diseases, although the exact link between aberrant miR-122 expression and liver disease pathogenesis remains unclear [32,33]. In our study, we aimed to investigate the functional role of miR-122 by examining changes in polyribosome-associated mRNA upon miR-122 overexpression and inhibition. Our results revealed that changes in miR-122 expression significantly affected the occupancy of multiple transcripts on polyribosomes, confirming the broad regulatory influence of miR-122. Interestingly, bioinformatics analysis identified the potential interplay between E2F4, NRF1, YY1, and FOXP3 transcription factors and a large number of miR-122-responsive genes. Notably, previous studies have linked NRF1 inactivation to the development of nonalcoholic steatohepatitis [30], while YY1 upregulation has been associated with fatty liver diseases [16], whereas E2F4 was found to promote HCC proliferation [34,35]. Furthermore, the interplay between NRF1 and E2F4 transcription factors was previously shown to contribute to cancer development and progression [36]. Based on these findings, we propose that miR-122 may “function” to “counteract” or “fine-tune” the activity of E2F4, NRF1 and YY1 thus contributing to the maintenance of liver homeostasis as proposed in Figure 8A. Conversely, the downregulation of miR-122 observed in patients with liver diseases, along with the consequent dysregulation of these transcription factors and their associate networks, may contribute to the “chronicization” process associated with liver diseases in these patients as proposed in Figure 8B. Interestingly, although FOXP3 was not detectable in Huh-7 cells, our data suggest that a subset of FOXP3-associated genes is expressed in these cells and responds to miR-122. FOXP3 is primarily expressed in Treg cells [17], where it plays a crucial role in immune tolerance by limiting immune activation of the microenvironment. Analysis of TCGA data further supports the prognostic value of FOXP3 in liver hepatocellular carcinoma (LIHC), with lower expression being unfavorable (**Supplementary Figure S5**). We propose that the increased secretion of miR-122 observed in patients with liver diseases [37] may lead to the downregulation of FOXP3-associated targets in Treg, hence contributing to the establishment of a pro-inflammatory milieu in the liver, as reported for miR-122 KO mice [4] and human liver organoid [7]. It is noteworthy that although there are a growing number of studies describing the role played by circulating miR-122 in the progression of human diseases, including cancer [38,39], to the best of our knowledge, our work is the first to suggest that cellular miR-122 may itself have a significant impact on the cellular secretome (see Figure 4). Altogether, the proposed activities of miR-122 on cellular secretome and on FOXP3 will be evaluate in planned follow-up studies.

**Figure 8:**
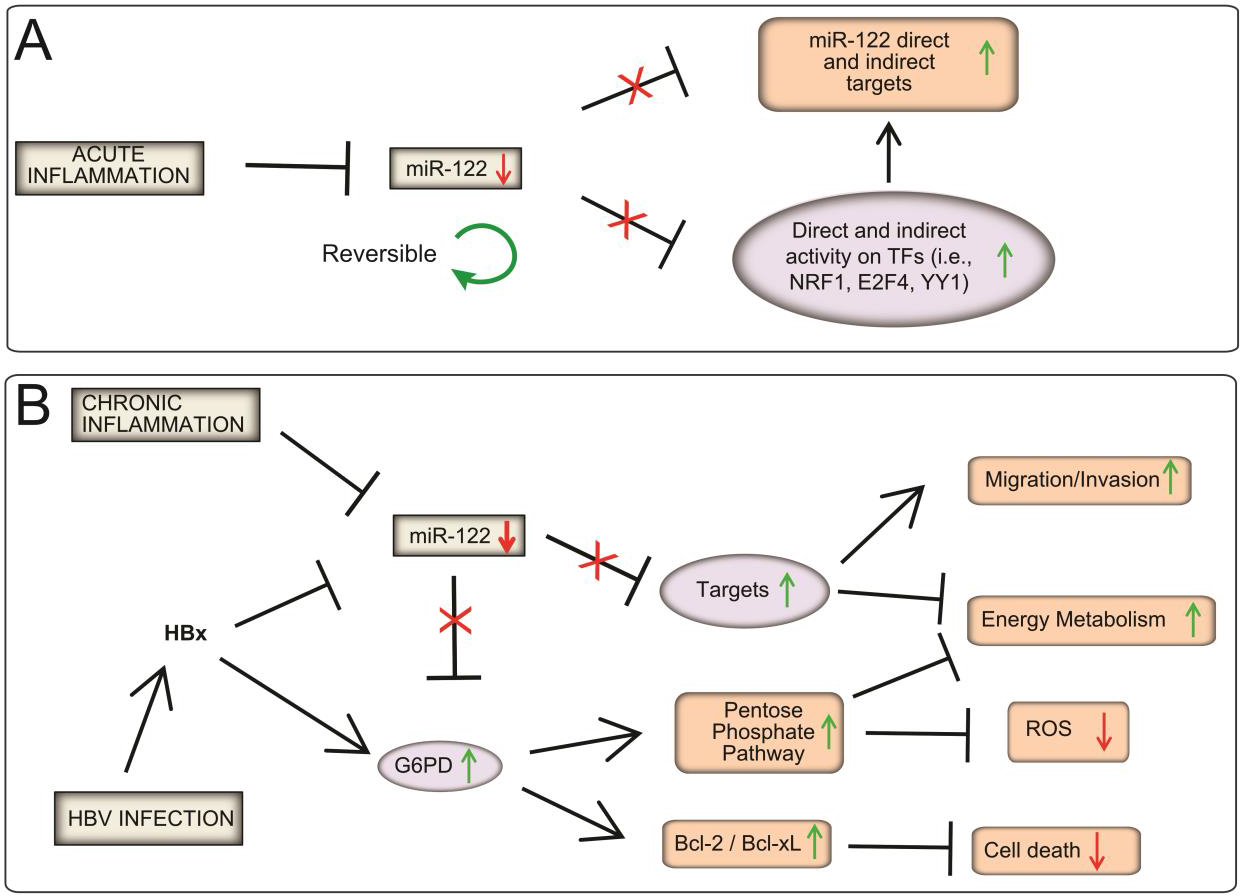
Proposed model for miR-122 in health and HBV-associated HCC. **(A) Physiological regulation of miR-122:** Physiological regulation of miR-122: miR-122 is transiently downregulated in response to (yet) unknown factors during inflammation, leading to upregulation of its targets and fine-tuning of gene expression downstream of NRF1, E2F4, and YY1 TFs. This model suggests the transient downregulation of miR-122 lead to the fine-tuning of immune response and regeneration-associated networks. **(B) Contribution of miR-122 in HBV-associated HCC:** We propose that the dysregulation of miR-122 expression is a central factor in HCC development among HBV-infected patients. We propose that miR-122 transcription is inhibited by inflammation (**model A**). In addition, in HBV-infected patients, the viral synthesized **HBx** has a dual role: i) inhibiting miR-122 [56] and ii) activating G6PD expression [30]. The combined downregulation of miR-122 and the activation of HBx lead to a "super activation" of G6PD, promoting cell survival (i.e., Bcl2/Bcl-xL activation; [44]), energy metabolism [30], and ROS detoxification [57]. We propose that these changes might promote the transformation of hepatocytes to a malignant phenotype in HBV-infected patients.

Although transcriptomic data provides valuable information about the cellular state, it is crucial to acknowledge that proteins are the primary agents responsible for cellular functions. Currently, the correlation between transcriptomic and proteomic data is limited [40–42]. The fact that changes in gene expression do not always lead to a parallel change in protein levels can be attributed both to technical constraints related to mass spectrometry, such as the inability to detect less abundant proteins [43], and to the effect of the post-transcriptional machinery on mRNA translation and protein synthesis. Therefore, transcriptomic data were combined with mass spectrometry analysis, confirming that a number of miR-122 responsive genes enriched on polyribosomes were also influenced at the protein level by changes in miR-122. Literature mining in combination with GEO/TCGA database analysis showed that a subset of miR-122-responsive proteins, including G6PD, CEP55, KIF11, SLC1A5, EPS15L1, TK1, and CLIC1, were upregulated in liver tumor tissues of HCC patients (**Supplementary Figure S8**), with their increased expression being unfavorable in liver cancer (**Supplementary Figure S9**). Overall, we strongly believe that assessing the impact of these genes on liver cancer’s development and progression is crucial next step, making further studies warranted.

Given the global distribution of hepatitis B, its association with poor treatment response, and the increased risk of HCC development, we specifically investigated the functional role of miR-122 in regulating G6PD in chronic hepatitis B (CHB) patients. Enhanced G6PD activity has been linked to the upregulation of anti-apoptotic factors such as Bcl-2 and Bcl-xL [30], which contribute to promoting cell growth and tumor development [44]. Additionally, increased G6PD activity may contribute to the accelerated turnover of the pentose phosphate pathway [45]. This pathway is crucial for cancer cell growth as it provides nucleic acids and NADPH for biosynthesis, as well as glutathione for combating oxidative stress. G6PD has been suggested as a potential therapeutic target in HCC, as inhibiting its activity could help prevent tumor progression [28]. In our study, we confirmed that G6PD was a direct target of miR-122, results that are consistent with other published research [26]. These findings support the hypothesis that HBV suppression of miR-122 may contribute directly to increased G6PD levels in the livers CHB patients. We propose that this may be an additional "risk factor" contributing to the increased risk of developing hepatocellular carcinoma in individuals with CHB (Figure 8B).

Overall, our research confirmed the central role of miR-122 in the regulation and fine-tuning of hepatic homeostasis. In this study, we discovered a new, hitherto uncharacterized molecular network downstream of miR-122, highlighting the possible interaction between miR-122 and the transcription factors NRF1, E2F4 and YY1. We propose that a combination of physiological and pathological signals, yet to be established, drives the downregulation of miR-122 in parenchymal cells and, possibly, its upregulation in Treg cells, thus contributing to the establishment of a pro-inflammatory and pro-tumorigenic environment in the liver.

## Materials and Methods

### Isolation and fractionation of polyribosomes

Polyribosomes were isolated from Huh-7 cells transfected with miR-122 mimic or miR-122 inhibitor for 48 h. Cells were treated with 200 µg/mL cycloheximide (Sigma-Aldrich, Taufkirchen, Germany, C4859-1ML) for 10 min, washed with PBS/100 µg/mL cycloheximide, and centrifuged (1000 rpm/4°C/5 min). The pelleted cells were lysed in 750 µL lysis buffer (5 mM Tris HCl pH 7.5, 1.5 mM KCl, 2.5 mM MgCl_2_, 0.2 mM cycloheximide) supplemented with 120 U/mL RNase Inhibitor (RiboLock, Thermo Fisher Scientific, Meerbusch, Germany, EO0381), 120 U/µL DNase I (New England BioLabs, Frankfurt a.M., Germany, M0303S), 0.5% sodium deoxycholate (Sigma-Aldrich, Taufkirchen, Germany, 30970-25G) and 0.5% Triton X-100. Cell nuclei were removed by centrifugation, the supernatant was layered on top of a linear sucrose gradient (10–50% sucrose) loaded in a SW40 Ti swinging-bucket rotor (Beckman Coulter, Krefeld, Germany) and centrifuged (33.500 rpm/4°C/3h, brake off) in a Optima XPN-80 ultracentrifuge (Beckman Coulter, Krefeld, Germany). Sucrose gradients were eluted into a BR-188 density gradient fractionation system (Brandel Inc, Gaithersburg, MD, USA), and nucleic acids passing through a UV recorder were detected at a wavelength of λ = 254 nm. Fractions of approximately 800 µL were collected and RNAs were recovered from each fraction by using phenol-chloroform-isoamyl alcohol (25:24:1, (v/v/v); Sigma-Aldrich, Taufkirchen, Germany, P2609) followed by ethanol precipitation.

### miR-122 overexpression and inhibition in Huh-7 cells

Huh-7 cells were transiently transfected with Lipofectamine RNAiMAX (Thermo Fisher Scientific, Meerbusch, Germany, 13778030) according to the manufactureŕs instructions with either 10 µM pre-miR-122 miRNA Precursor (mirVana miRNA mimic, Thermo Fisher Scientific, Meerbusch, Germany, 4464066), or 20 µM miR-122 antagomiR (Miravirsen, SPC-3649, Roche, Mannheim, Germany, formely Santaris Pharma) or scrambled oligo control (SPC-3744, Roche, Mannheim, Germany, formely Santaris Pharma). The sequence and efficacy of antagomiR (122-PM/SPC-3649) and scrambled oligo (SCR/ SPC-3744) have been published by the authors in a previous work [3].

### RNA isolation, QC and qPCR analysis

RNA isolation for qPCR analysis was performed with QIAzol Lysis Reagent (Qiagen, Hilden, Germany, 79306) followed by purification with miRNeasy Mini Kit (Qiagen, Hilden, Germany, 217004) according to the manufactureŕs instructions. RNA was quantified using Qubit Fluorometer 2.0 (Thermo Fisher Scientific, Meerbusch, Germany). RNA isolation of PFA liver sections was performed as described in the **Supplementary Data S1**. cDNA synthesis for mRNA quantification was carried out as described in [3]. cDNA synthesis for miRNAs detection by qPCR was performed with the miQPCR method [46]. Notably, this method enables the universal reverse transcription of all the miRNAs contained in the reverse transcribed RNAs. qPCR reactions were carried out on a StepOne Plus cycler (Thermo Fisher Scientific, Meerbusch, Germany) and amplicons were detected by using SYBR Green I (GO-Taq PCR Master mix, Promega, Walldorf, Germany; A6002). Suitable reference genes were identified by using geNorm [47], and relative quantities were calculated with ΔΔCt [48]. qPCR data were analyzed by qBase v.1.3.5 [49]. Primers used for miRNA and mRNA quantification are listed in **Supplementary Data S2**.

### Cloning and target validation by Dual-Luciferase reporter assay

*RNA22* [50] was used to predicts potential miR-122 binding sites in the 3’UTRs of CEP55, CLIC1, EPS15L1, G6PD, KIF11, SLC1A5, and TK1 (**Supplementary Figure S11**). Full lengths 3’UTR of human CEP55, CLIC1, G6PD, EPS15L1, KIF11, SLC1A5, and TK1 were amplified by using Expand High Fidelity PCR System (Roche, Mannheim, Germany, 11732641001) as indicated by the vendor. Primers used for 3’UTR amplification are listed in **Supplementary Data S2**. The resulting 3’UTRs were cloned into pMIR(+) and pMIR(-) reporter vectors as previously described [3]. Constructs carrying the 3’UTRs were transiently co-transfected with vectors carrying Renilla-Luciferase and either miR-122 mimic (122-MIM) or scramble oligos (SCR) into HEK293 cells with Lipofectamine RNAiMAX (Thermo Fisher Scientific, Meerbusch, Germany, 13778030) according to the manufactureŕs instructions. Dual-Luciferase Reporter Assay (Promega, Walldorf, Germany, E1910) was conducted in accordance with the manufactureŕs instructions. The chemiluminescence assay was performed in white opaque 96-well plates with 50 µL cell lysate and 50 µL of both, LARII reagent (Firefly luciferase substrate) and Stop and Glo reagent (Renilla luciferase substrate), in a GloMax Multi Plus Multiplate Reader (Promega, Walldorf, Germany) according to the preset protocol with an integrity time of 10 seconds for each read. Data were normalized by calculating ratios of Firefly/Renilla activities to correct for possible variations in transfection efficiencies.

### Affymetrix microarray and Gene Ontology analyses

RNA was isolated from polyribosomal fractions as described above. In order to assess RNA integrity, 1 µL of individual polyribosomal RNAs were loaded on a PicoChip and run on a 2100 Bioanalyzer (Agilent Technologies, Santa Clara, CA, USA), according to the supplied protocol. Following QC and pooling, pooled RNA were hybridized to Affymetrix Gene Chip Human Gene 1.0 ST arrays (Thermo Fisher Scientific, Meerbusch, Germany, 901086). cDNA syntheses, labeling and hybridizytions were performed by the genomics core facility of the European Molecular Biology Laboratory (EMBL, Heidelberg, Germany). Normalization and data analysis was carried out with AltAnalyze [14]. Microarray data are available via the Gene Expression Omnibus (GEO; [51]) repository (accession number **GSE234690**). Gene ontology (GO) enrichment analysis was performed by using ShinyGo [22] and GOrilla [52].

### Protein isolation, Western blot analysis and antibodies

Protein were isolated by using RIPA buffer with protease inhibitor cocktail (cOmplete Protease Inhibitor Cocktail, Roche, Mannheim, Germany, 04693116001), and concentrations were measured using the Qubit Protein Assay Kit (Thermo Fisher Scientific, Meerbusch, Germany, Q33211) according to the manufacturer’s instructions. Western blot analyses were carried out using 20 µg of total proteins. Protein lysates were loaded together with 10 µL of Pro-tein Ladder (BioRad, Dusseldorf, Germany, 161-0373) on 10% or 12% SDS polyacrylamide gels and transferred to nitrocellulose membranes using semidry blotting systems according to standard protocols. Membranes were blocked with 5% milk powder (Carl Roth, Karlsruhe, Germany, T145.3) in Tris-buffered saline with Tween20 (TBST) and incubated with incubation primary antibody for 1h at RT or overnight at 4 °C, Following, the appropriate horseradish peroxidase (HRP)-coupled secondary antibody was incubated for 2h at room temperature. Chemiluminescence was detected with ECL Western blotting Substrate (Promega, Walldorf, Germany, W1001) using the ChemiDoc MP Imaging System (BioRad, Dusseldorf, Germany). Signal intensities of Western blot protein bands were analyzed using ImageLab (BioRad, Dusseldorf, Germany, version 6.0.1). Antibodies used in protein analysis by Western blot: anti-EPS15L1 antibody (Abcam, Cambridge, UK, ab53006), anti-G6PD antibody (Sigma-Aldrich Chemie GmbH, Taufkirchen, Germany, HPA000834), anti-β Actin antibody (Abcam, Cambridge, UK, ab8226), anti-GAPDH antibody (AbD Serotec, Puchheim, Germany, MCA4739).

### Sample preparation and proteome analysis by liquid chromatography-tandem mass spectrometry (LC-MS/MS)

Proteins were isolated from transfected cells or liver tissue as described above. For proteomics, LC-MS/MS was performed essentially as described by Schira *et al.* [53,54]. Detailed procedures and technical information is given in the **Supplementary Data S1**.

### Statistical analysis and imaging software

Statistical analyses were carried out using GraphPad Prism (version 9.4.1). Data are expressed as mean ± SD. Results compared between groups were analyzed by 2-tailed Student’s t test when two samples were considered or by 1-way ANOVA for three or more samples. When groups were compared using 1-way ANOVA, we assumed that data were normally distributed. The data were considered significant at a p value ≤ 0.05. Images were prepared using Affinity Designer (version 1.10.4.1198).

### Data mining from the Gene Expression Omnibus (GEO) repository, and data analysis

The miRNA and mRNA profiling data from animal models and liver cancer patients were retrieved from the GEO repository [51]. Enrichment analysis was performed with ShinyGO [22] by uploading list of significantly upregulated genes, which were predicted to be miR-122 targets by *miRWalk* [21] target prediction database.

### The Cancer Genome Atlas (TCGA) data and Kaplan-Meier survival curves

This study includes data from the TCGA Research Network (https://www.cancer.gov/tcga), and uses the KM Plotter [55] to generate Kaplan-Meier survival curves for hepatocellular carcinoma patients.

## Author Contributions

M.P., designed research, performed experiments and wrote the manuscript. J.S. and K.S., carried out protein extraction, proteome analysis including statistical analysis. R.P. and T.L. (Longerich), provided PFA fixed specimens from HCC patients. L.R.H., J.B. and U.P.N., curated and provided frozen specimens from HCC patients. T.L. (Luedde), contributed to research. M.C., conceived, designed and performed experiments, and wrote the manuscript.

All authors read and approved the manuscript.

## Institutional Review Board Statement

The ethic committee of the Medical Faculty of RWTH Aachen gave the approval (ethical vote EK122/16) for using human HCC tissue samples, in conformity with the Declaration of Helsinki ethical guidelines.

## Informed Consent Statement

Informed consent was obtained from all subjects involved in the study.

## Data Availability Statement

All study data are provided in the manuscript.

## Acknowledgements

The authors are grateful to Claudia Rupprecht (University of Düsseldorf, Düsseldorf, Germany) and Janina Thies (University of Düsseldorf, Düsseldorf, Germany) for the technical assistance. We would also like to thank the EMBL Genomics Core facility (EMBL, Heidelberg, Germany) for hybridizing the polyribosomal RNAs on Affymetrix arrays.

## List of Abbreviations

miRNA: microRNA
HCC: hepatocellular carcinoma
GOI: Gene of interest
GEO: Gene Expression Omnibus
TCGA: The Cancer Genome Atlas
GO: Gene ontology
122-MIM: miR-122 mimic
122-PM: miR-122 inhibitor
G6PD: glucose-6-phosphate dehydrogenase
HBV: hepatitis B virus
CHB: chronic hepatitis B
HBx: hepatitis B viral protein X
qPCR: quantitative *real-time* PCR
SLC7A1: Solute Carrier Family 7 Member 1
YY1: Yin Yang 1
FOXP3: Forkhead box P3
E2F4: E2F transcription factor 4
NRF1: nuclear respiratory factor 1
TFs: transcription factors
CEP55: centrosomal protein 55
KIF11: kinesin family member 11
SLC1A5: solute carrier family 1 member 5
EPS15L: epidermal growth factor receptor pathway substrate 15 Like 1
TK1: thymidine kinase 1
CLIC1: chloride intracellular channel 1

Abbreviations for additional gene names are given in the **Supplementary Data S2**.

## References

1. Navarro, V.J.; Senior, J.R. Drug-related hepatotoxicity. N Engl J Med 2006, 354, 731–739, doi:10.1056/NEJMra052270.

2. Elmen, J.; Lindow, M.; Silahtaroglu, A.; Bak, M.; Christensen, M.; Lind-Thomsen, A.; Hedtjarn, M.; Hansen, J.B.; Hansen, H.F.; Straarup, E.M.;, et al. Antagonism of microRNA-122 in mice by systemically administered LNA-antimiR leads to up-regulation of a large set of predicted target mRNAs in the liver. Nucleic Acids Res 2008, 36, 1153–1162, doi:10.1093/nar/gkm1113.

3. Castoldi, M.; Vujic Spasic, M.; Altamura, S.; Elmen, J.; Lindow, M.; Kiss, J.; Stolte, J.; Sparla, R.; D’Alessandro, L.A.; Klingmuller, U.;, et al. The liver-specific microRNA miR-122 controls systemic iron homeostasis in mice. J Clin Invest 2011, 121, 1386–1396, doi:10.1172/JCI44883.

4. Hsu, S.H.; Wang, B.; Kota, J.; Yu, J.; Costinean, S.; Kutay, H.; Yu, L.; Bai, S.; La Perle, K.; Chivukula, R.R.;, et al. Essential metabolic, anti-inflammatory, and anti-tumorigenic functions of miR-122 in liver. J Clin Invest 2012, 122, 2871–2883, doi:10.1172/JCI63539.

5. Callegari, E.; Elamin, B.K.; Sabbioni, S.; Gramantieri, L.; Negrini, M. Role of microRNAs in hepatocellular carcinoma: a clinical perspective. Onco Targets Ther 2013, 6, 1167–1178, doi:10.2147/OTT.S36161.

6. Ha, S.Y.; Yu, J.I.; Choi, C.; Kang, S.Y.; Joh, J.W.; Paik, S.W.; Kim, S.; Kim, M.; Park, H.C.; Park, C.K. Prognostic significance of miR-122 expression after curative resection in patients with hepatocellular carcinoma. Sci Rep 2019, 9, 14738, doi:10.1038/s41598-019-50594-2.

7. Sendi, H.; Mead, I.; Wan, M.; Mehrab-Mohseni, M.; Koch, K.; Atala, A.; Bonkovsky, H.L.; Bishop, C.E. miR-122 inhibition in a human liver organoid model leads to liver inflammation, necrosis, steatofibrosis and dysregulated insulin signaling. PLoS One 2018, 13, e0200847, doi:10.1371/journal.pone.0200847.

8. Beilharz, T.H.; Humphreys, D.T.; Clancy, J.L.; Thermann, R.; Martin, D.I.; Hentze, M.W.; Preiss, T. microRNA-mediated messenger RNA deadenylation contributes to translational repression in mammalian cells. PLoS One 2009, 4, e6783, doi:10.1371/journal.pone.0006783.

9. Macfarlane, L.A.; Murphy, P.R. MicroRNA: Biogenesis, Function and Role in Cancer. Curr Genomics 2010, 11, 537–561, doi:10.2174/138920210793175895.

10. Hendrickson, D.G.; Hogan, D.J.; McCullough, H.L.; Myers, J.W.; Herschlag, D.; Ferrell, J.E.; Brown, P.O. Concordant regulation of translation and mRNA abundance for hundreds of targets of a human microRNA. PLoS Biol 2009, 7, e1000238, doi:10.1371/journal.pbio.1000238.

11. Guo, H.; Ingolia, N.T.; Weissman, J.S.; Bartel, D.P. Mammalian microRNAs predominantly act to decrease target mRNA levels. Nature 2010, 466, 835–840, doi:10.1038/nature09267.

12. Cloonan, N. Re-thinking miRNA-mRNA interactions: intertwining issues confound target discovery. Bioessays 2015, 37, 379–388, doi:10.1002/bies.201400191.

13. Chang, J.; Nicolas, E.; Marks, D.; Sander, C.; Lerro, A.; Buendia, M.A.; Xu, C.; Mason, W.S.; Moloshok, T.; Bort, R.;, et al. miR-122, a mammalian liver-specific microRNA, is processed from hcr mRNA and may downregulate the high affinity cationic amino acid transporter CAT-1. RNA Biol 2004, 1, 106–113, doi:10.4161/rna.1.2.1066.

14. Emig, D.; Salomonis, N.; Baumbach, J.; Lengauer, T.; Conklin, B.R.; Albrecht, M. AltAnalyze and DomainGraph: analyzing and visualizing exon expression data. Nucleic Acids Res 2010, 38, W755–762, doi:10.1093/nar/gkq405.

15. Zambon, A.C.; Gaj, S.; Ho, I.; Hanspers, K.; Vranizan, K.; Evelo, C.T.; Conklin, B.R.; Pico, A.R.; Salomonis, N. GO-Elite: a flexible solution for pathway and ontology over-representation. Bioinformatics 2012, 28, 2209–2210, doi:10.1093/bioinformatics/bts366.

16. Lu, Y.; Ma, Z.; Zhang, Z.; Xiong, X.; Wang, X.; Zhang, H.; Shi, G.; Xia, X.; Ning, G.; Li, X. Yin Yang 1 promotes hepatic steatosis through repression of farnesoid X receptor in obese mice. Gut 2014, 63, 170–178, doi:10.1136/gutjnl-2012-303150.

17. Speletas, M.; Argentou, N.; Germanidis, G.; Vasiliadis, T.; Mantzoukis, K.; Patsiaoura, K.; Nikolaidis, P.; Karanikas, V.; Ritis, K.; Germenis, A.E. Foxp3 expression in liver correlates with the degree but not the cause of inflammation. Mediators Inflamm 2011, 2011, 827565, doi:10.1155/2011/827565.

18. Yang, X.; Zu, X.; Tang, J.; Xiong, W.; Zhang, Y.; Liu, F.; Jiang, Y. Zbtb7 suppresses the expression of CDK2 and E2F4 in liver cancer cells: implications for the role of Zbtb7 in cell cycle regulation. Mol Med Rep 2012, 5, 1475–1480, doi:10.3892/mmr.2012.846.

19. Xu, Z.; Chen, L.; Leung, L.; Yen, T.S.; Lee, C.; Chan, J.Y. Liver-specific inactivation of the Nrf1 gene in adult mouse leads to nonalcoholic steatohepatitis and hepatic neoplasia. Proc Natl Acad Sci U S A 2005, 102, 4120–4125, doi:10.1073/pnas.0500660102.

20. Darpolor, M.M.; Basu, S.S.; Worth, A.; Nelson, D.S.; Clarke-Katzenberg, R.H.; Glickson, J.D.; Kaplan, D.E.; Blair, I.A. The aspartate metabolism pathway is differentiable in human hepatocellular carcinoma: transcriptomics and (13) C-isotope based metabolomics. NMR Biomed 2014, 27, 381–389, doi:10.1002/nbm.3072.

21. Sticht, C.; De La Torre, C.; Parveen, A.; Gretz, N. miRWalk: An online resource for prediction of microRNA binding sites. PLoS One 2018, 13, e0206239, doi:10.1371/journal.pone.0206239.

22. Ge, S.X.; Jung, D.; Yao, R. ShinyGO: a graphical gene-set enrichment tool for animals and plants. Bioinformatics 2020, 36, 2628–2629, doi:10.1093/bioinformatics/btz931.

23. Burchard, J.; Zhang, C.; Liu, A.M.; Poon, R.T.; Lee, N.P.; Wong, K.F.; Sham, P.C.; Lam, B.Y.; Ferguson, M.D.; Tokiwa, G.;, et al. microRNA-122 as a regulator of mitochondrial metabolic gene network in hepatocellular carcinoma. Mol Syst Biol 2010, 6, 402, doi:10.1038/msb.2010.58.

24. Liu, A.M.; Xu, Z.; Shek, F.H.; Wong, K.F.; Lee, N.P.; Poon, R.T.; Chen, J.; Luk, J.M. miR-122 targets pyruvate kinase M2 and affects metabolism of hepatocellular carcinoma. PLoS One 2014, 9, e86872, doi:10.1371/journal.pone.0086872.

25. Consortium, E.P. An integrated encyclopedia of DNA elements in the human genome. Nature 2012, 489, 57–74, doi:10.1038/nature11247.

26. Barajas, J.M.; Reyes, R.; Guerrero, M.J.; Jacob, S.T.; Motiwala, T.; Ghoshal, K. The role of miR-122 in the dysregulation of glucose-6-phosphate dehydrogenase (G6PD) expression in hepatocellular cancer. Sci Rep 2018, 8, 9105, doi:10.1038/s41598-018-27358-5.

27. Yang, H.C.; Stern, A.; Chiu, D.T. G6PD: A hub for metabolic reprogramming and redox signaling in cancer. Biomed J 2021, 44, 285–292, doi:10.1016/j.bj.2020.08.001.

28. Hu, H.; Ding, X.; Yang, Y.; Zhang, H.; Li, H.; Tong, S.; An, X.; Zhong, Q.; Liu, X.; Ma, L.;, et al. Changes in glucose-6-phosphate dehydrogenase expression results in altered behavior of HBV-associated liver cancer cells. Am J Physiol Gastrointest Liver Physiol 2014, 307, G611–622, doi:10.1152/ajpgi.00160.2014.

29. Dore, M.P.; Vidili, G.; Marras, G.; Assy, S.; Pes, G.M. Inverse Association between Glucose‒6‒ Phosphate Dehydrogenase Deficiency and Hepatocellular Carcinoma. Asian Pac J Cancer Prev 2018, 19, 1069–1073, doi:10.22034/APJCP.2018.19.4.1069.

30. Liu, B.; Fang, M.; He, Z.; Cui, D.; Jia, S.; Lin, X.; Xu, X.; Zhou, T.; Liu, W. Hepatitis B virus stimulates G6PD expression through HBx-mediated Nrf2 activation. Cell Death Dis 2015, 6, e1980, doi:10.1038/cddis.2015.322.

31. Wang, S.; Qiu, L.; Yan, X.; Jin, W.; Wang, Y.; Chen, L.; Wu, E.; Ye, X.; Gao, G.F.; Wang, F.;, et al. Loss of microRNA 122 expression in patients with hepatitis B enhances hepatitis B virus replication through cyclin G(1)-modulated P53 activity. Hepatology 2012, 55, 730–741, doi:10.1002/hep.24809.

32. Akuta, N.; Kawamura, Y.; Suzuki, F.; Saitoh, S.; Arase, Y.; Fujiyama, S.; Sezaki, H.; Hosaka, T.; Kobayashi, M.; Suzuki, Y.;, et al. Analysis of association between circulating miR-122 and histopathological features of nonalcoholic fatty liver disease in patients free of hepatocellular carcinoma. BMC Gastroenterol 2016, 16, 141, doi:10.1186/s12876-016-0557-6.

33. Dubin, P.H.; Yuan, H.; Devine, R.K.; Hynan, L.S.; Jain, M.K.; Lee, W.M.; Acute Liver Failure Study, G. Micro-RNA-122 levels in acute liver failure and chronic hepatitis C. J Med Virol 2014, 86, 1507-1514, doi:10.1002/jmv.23987.

34. Zheng, Q.; Fu, Q.; Xu, J.; Gu, X.; Zhou, H.; Zhi, C. Transcription factor E2F4 is an indicator of poor prognosis and is related to immune infiltration in hepatocellular carcinoma. J Cancer 2021, 12, 1792–1803, doi:10.7150/jca.51616.

35. Liu, J.; Xia, L.; Wang, S.; Cai, X.; Wu, X.; Zou, C.; Shan, B.; Luo, M.; Wang, D. E2F4 Promotes the Proliferation of Hepatocellular Carcinoma Cells through Upregulation of CDCA3. J Cancer 2021, 12, 5173–5180, doi:10.7150/jca.53708.

36. Bhawe, K.; Roy, D. Interplay between NRF1, E2F4 and MYC transcription factors regulating common target genes contributes to cancer development and progression. Cell Oncol (Dordr) 2018, 41, 465-484, doi:10.1007/s13402-018-0395-3.

37. Roderburg, C.; Benz, F.; Vargas Cardenas, D.; Koch, A.; Janssen, J.; Vucur, M.; Gautheron, J.; Schneider, A.T.; Koppe, C.; Kreggenwinkel, K.;, et al. Elevated miR-122 serum levels are an independent marker of liver injury in inflammatory diseases. Liver Int 2015, 35, 1172–1184, doi:10.1111/liv.12627.

38. Li, C.; Qin, F.; Wang, W.; Ni, Y.; Gao, M.; Guo, M.; Sun, G. hnRNPA2B1-Mediated Extracellular Vesicles Sorting of miR-122-5p Potentially Promotes Lung Cancer Progression. Int J Mol Sci 2021, 22, doi:10.3390/ijms222312866.

39. Hosen, M.R.; Goody, P.R.; Zietzer, A.; Xiang, X.; Niepmann, S.T.; Sedaghat, A.; Tiyerili, V.; Chennupati, R.; Moore, J.B.t.; Boon, R.A.;, et al. Circulating MicroRNA-122-5p Is Associated With a Lack of Improvement in Left Ventricular Function After Transcatheter Aortic Valve Replacement and Regulates Viability of Cardiomyocytes Through Extracellular Vesicles. Circulation 2022, 146, 1836–1854, doi:10.1161/CIRCULATIONAHA.122.060258.

40. Buccitelli, C.; Selbach, M. mRNAs, proteins and the emerging principles of gene expression control. Nat Rev Genet 2020, 21, 630–644, doi:10.1038/s41576-020-0258-4.

41. Perl, K.; Ushakov, K.; Pozniak, Y.; Yizhar-Barnea, O.; Bhonker, Y.; Shivatzki, S.; Geiger, T.; Avraham, K.B.; Shamir, R. Reduced changes in protein compared to mRNA levels across non-proliferating tissues. BMC Genomics 2017, 18, 305, doi:10.1186/s12864-017-3683-9.

42. Liu, Y.; Beyer, A.; Aebersold, R. On the Dependency of Cellular Protein Levels on mRNA Abundance. Cell 2016, 165, 535–550, doi:10.1016/j.cell.2016.03.014.

43. Fricker, L.D. Limitations of Mass Spectrometry-Based Peptidomic Approaches. J Am Soc Mass Spectrom 2015, 26, 1981–1991, doi:10.1007/s13361-015-1231-x.

44. Hu, T.; Zhang, C.; Tang, Q.; Su, Y.; Li, B.; Chen, L.; Zhang, Z.; Cai, T.; Zhu, Y. Variant G6PD levels promote tumor cell proliferation or apoptosis via the STAT3/5 pathway in the human melanoma xenograft mouse model. BMC Cancer 2013, 13, 251, doi:10.1186/1471-2407-13-251.

45. Stanton, R.C. Glucose-6-phosphate dehydrogenase, NADPH, and cell survival. IUBMB Life 2012, 64, 362–369, doi:10.1002/iub.1017.

46. Benes, V.; Collier, P.; Kordes, C.; Stolte, J.; Rausch, T.; Muckentaler, M.U.; Haussinger, D.; Castoldi, M. Identification of cytokine-induced modulation of microRNA expression and secretion as measured by a novel microRNA specific qPCR assay. Sci Rep 2015, 5, 11590, doi:10.1038/srep11590.

47. Schlotter, Y.M.; Veenhof, E.Z.; Brinkhof, B.; Rutten, V.P.; Spee, B.; Willemse, T.; Penning, L.C. A GeNorm algorithm-based selection of reference genes for quantitative real-time PCR in skin biopsies of healthy dogs and dogs with atopic dermatitis. Vet Immunol Immunopathol 2009, 129, 115–118, doi:10.1016/j.vetimm.2008.12.004.

48. Livak, K.J.; Schmittgen, T.D. Analysis of relative gene expression data using real-time quantitative PCR and the 2(-Delta Delta C(T)) Method. Methods 2001, 25, 402–408, doi:10.1006/meth.2001.1262.

49. Hellemans, J.; Mortier, G.; De Paepe, A.; Speleman, F.; Vandesompele, J. qBase relative quantification framework and software for management and automated analysis of real-time quantitative PCR data. Genome Biol 2007, 8, R19, doi:10.1186/gb-2007-8-2-r19.

50. Loher, P.; Rigoutsos, I. Interactive exploration of RNA22 microRNA target predictions. Bioinformatics 2012, 28, 3322–3323, doi:10.1093/bioinformatics/bts615.

50. Patra, B.G.; Roberts, K.; Wu, H. A content-based dataset recommendation system for researchers-a case study on Gene Expression Omnibus (GEO) repository. Database (Oxford) 2020, 2020, 1, doi:10.1093/database/baaa064.

52. Eden, E.; Navon, R.; Steinfeld, I.; Lipson, D.; Yakhini, Z. GOrilla: a tool for discovery and visualization of enriched GO terms in ranked gene lists. BMC Bioinformatics 2009, 10, 48, doi:10.1186/1471-2105-10-48.

53. Schira, J.; Falkenberg, H.; Hendricks, M.; Waldera-Lupa, D.M.; Kogler, G.; Meyer, H.E.; Muller, H.W.; Stuhler, K. Characterization of Regenerative Phenotype of Unrestricted Somatic Stem Cells (USSC) from Human Umbilical Cord Blood (hUCB) by Functional Secretome Analysis. Mol Cell Proteomics 2015, 14, 2630–2643, doi:10.1074/mcp.M115.049312.

54. Falkenberg, H.; Radke, T.F.; Kogler, G.; Stuhler, K. Proteomic Profiling of Ex Vivo Expanded CD34-Positive Haematopoetic Cells Derived from Umbilical Cord Blood. Stem Cells Int 2013, 2013, 245695, doi:10.1155/2013/245695.

54. Nagy, A.; Lanczky, A.; Menyhart, O.; Gyorffy, B. Validation of miRNA prognostic power in hepatocellular carcinoma using expression data of independent datasets Sci Rep 2018, 8, 9227, doi:10.1038/s41598-018-27521-y.

56. Bandopadhyay, M.; Sarkar, N.; Datta, S.; Das, D.; Pal, A.; Panigrahi, R.; Banerjee, A.; Panda, C.K.; Das, C.; Chakrabarti, S.;, et al. Hepatitis B virus X protein mediated suppression of miRNA-122 expression enhances hepatoblastoma cell proliferation through cyclin G1-p53 axis. Infect Agent Cancer 2016, 11, 40, doi:10.1186/s13027-016-0085-6.

57. Nobrega-Pereira, S.; Fernandez-Marcos, P.J.; Brioche, T.; Gomez-Cabrera, M.C.; Salvador-Pascual, A.; Flores, J.M.; Vina, J.; Serrano, M. G6PD protects from oxidative damage and improves healthspan in mice. Nat Commun 2016, 7, 10894, doi:10.1038/ncomms10894.

