## Supplementary Data for "Uncovering novel roles of miR-122 in the pathophysiology of the liver: Potential interaction with NRF1 and E2F4 signaling"

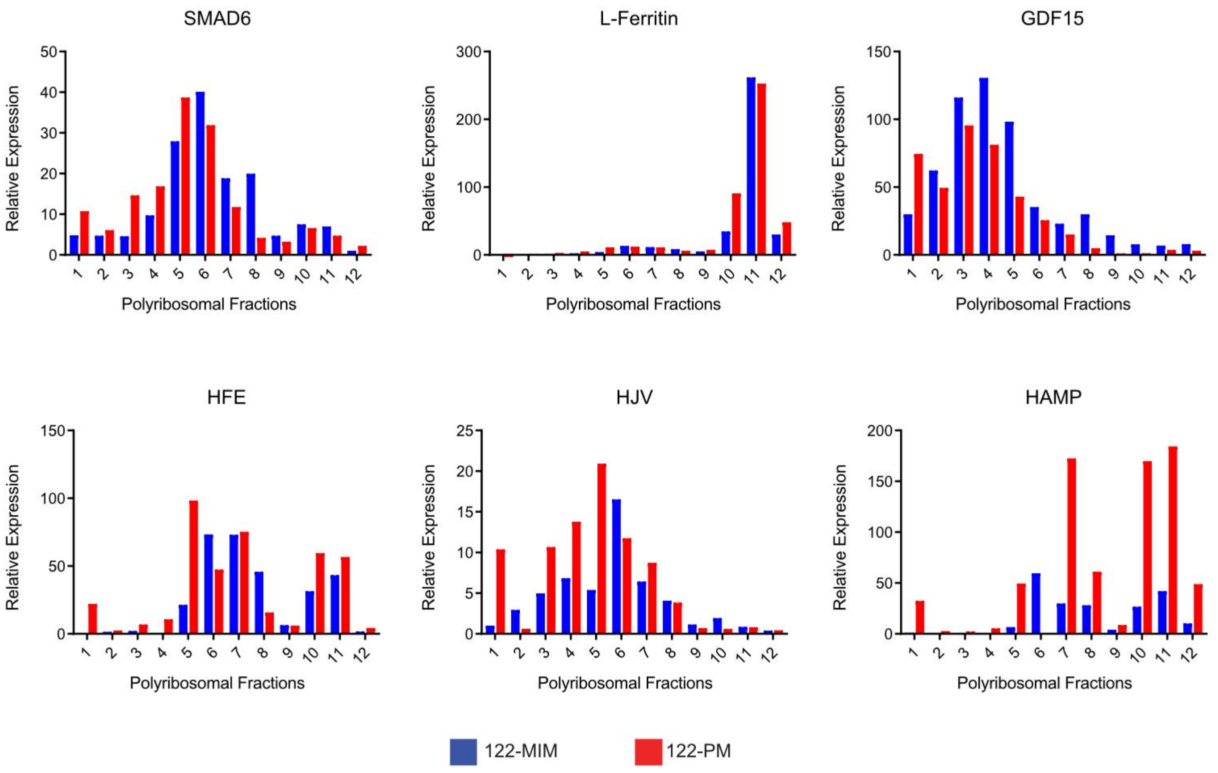


**Supplementary Figure S1 Distribution of non-responsive and miR-122 responsive gene on polyribosomes.** Additional qPCR quantification of mRNAs distribution for GOIs on the individual polyribosomes was categorized into two panels: those genes that are not anticipated to be influenced by alterations in miR-122 levels (SMAD6, L-Ferritin and GDF15; **top panel**), and those genes that have been previously demonstrated to be directly (HFE and HJV) or indirectly (HAMP) regulated by miR-122 (**lower panel**).

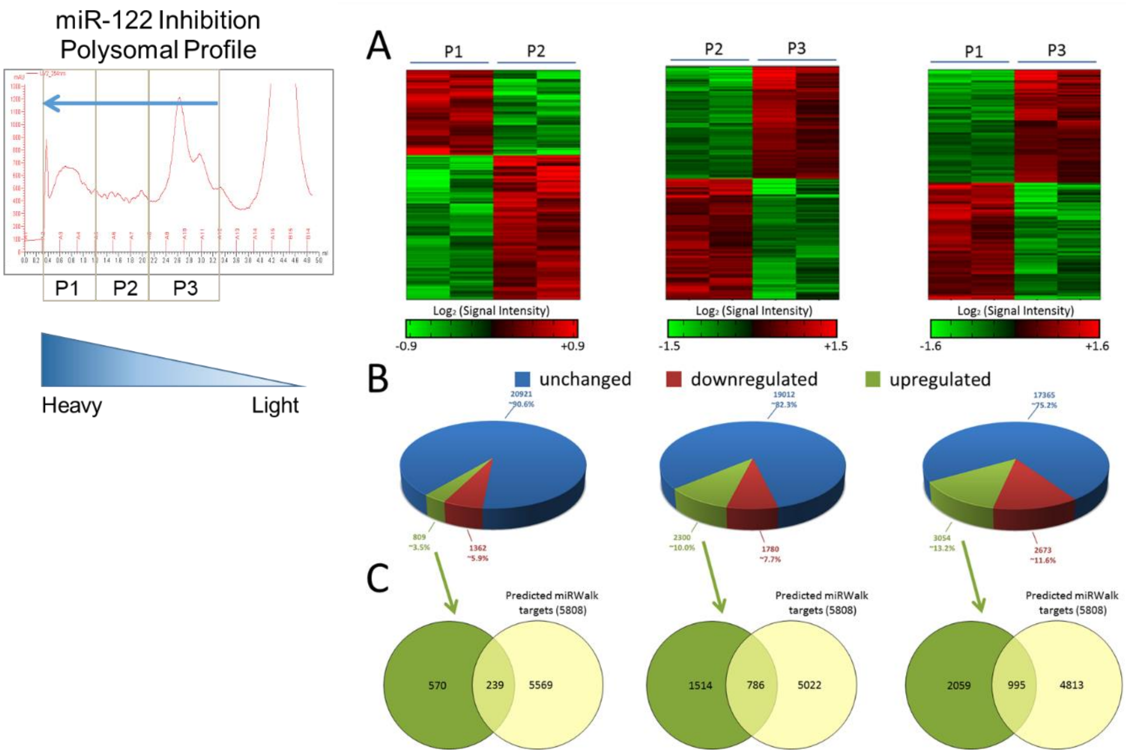


**Supplementary Figure S2: Identification of genes responsive to miR-122 inhibition**

AltAnalyze comparison of the polyribosomal pools in cells transfected with miR-122 inhibitors (122-PM).

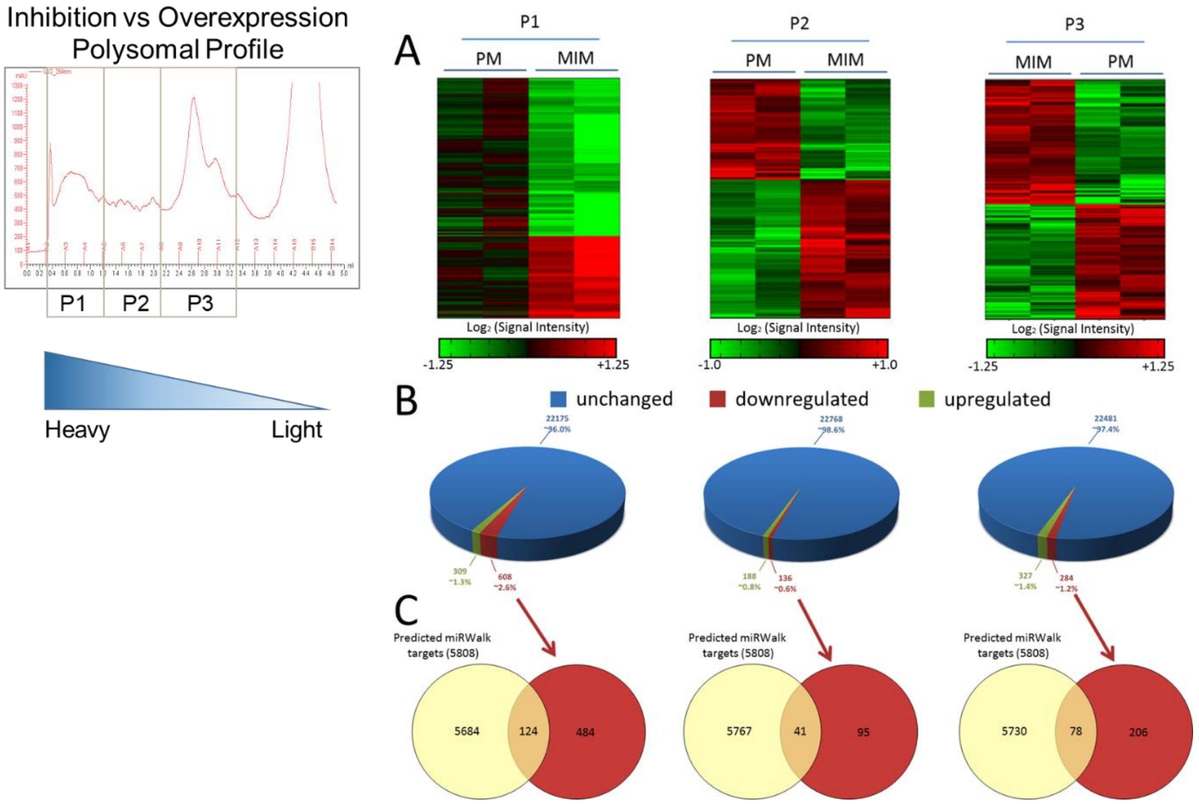


**Supplementary Figure S3: Identification of genes responsive to miR-122 inhibition and overexpression**

AltAnalyze comparison of the polyribosomal pools in cells transfected with miR-122 inhibitors (122-PM) compared to miR-122 overexpression (122-MIM).

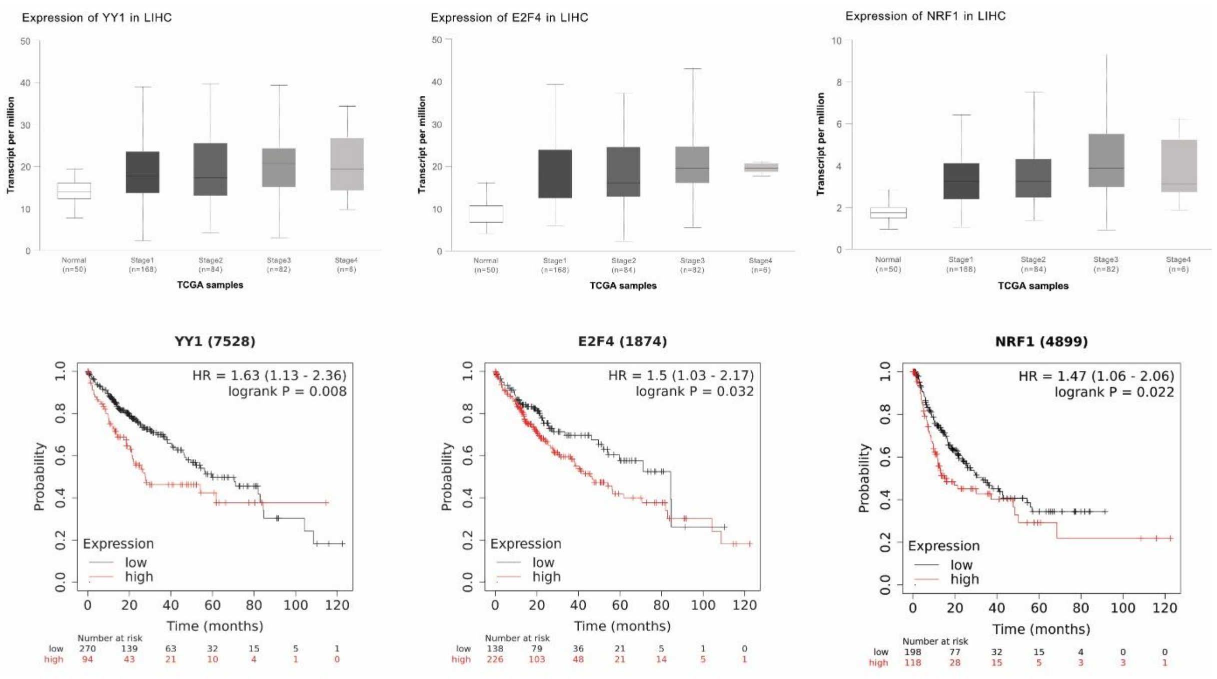


**Supplementary Figure S4: Kaplan-Meier survival curve for YY1, E2F4 and NRF1**

(**Top panel**) Mining of sequencing data for YY1, E2F4 and NRF1 in cohort of patients with liver hepatocellular carcinoma from TCGA. (**Bottom panel**) The Kaplan-Meier curve analysis of the liver hepatocellular carcinoma (LIHC) cohorts from TCGA reveals important prognostic factors for liver cancer. Specifically, lower expression levels of YY1, E2F4, and NRF1 transcription factors are correlated with higher chances of survival.

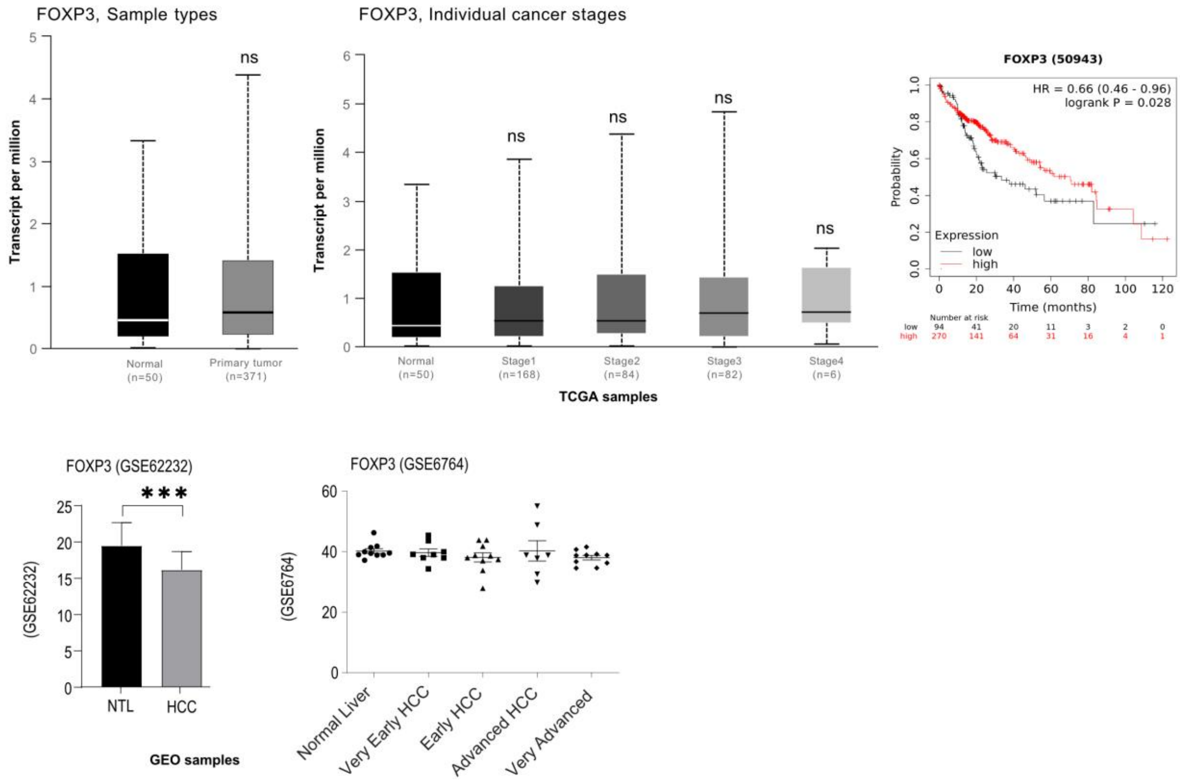


**Supplementary Figure S5: Higher FOXP3 expression is associated with better survival**

In contrast, to what observed for YY1, E2F4, and NRF1 TFs, no significant differences were found for FOXP3 expression in the TCGA LIHC-cohort. The Kaplan-Meier curve suggests that LIHC patients with higher FOXP3 expression had a significantly higher probability of survival. Query of the GEO database identified that FOXP3 expression was significantly reduced in the liver of HCC patients (GSE62232), but was found unchanged in a second cohort (GSE6764). Data are shown mean ± SD. Statistical analysis was performed with two-tailed T-test between two samples, or One-way ANOVA for three or more samples, $p \leq 0.05$ was considered significant. ns = Not significant; ***, $p \leq 0.001$.

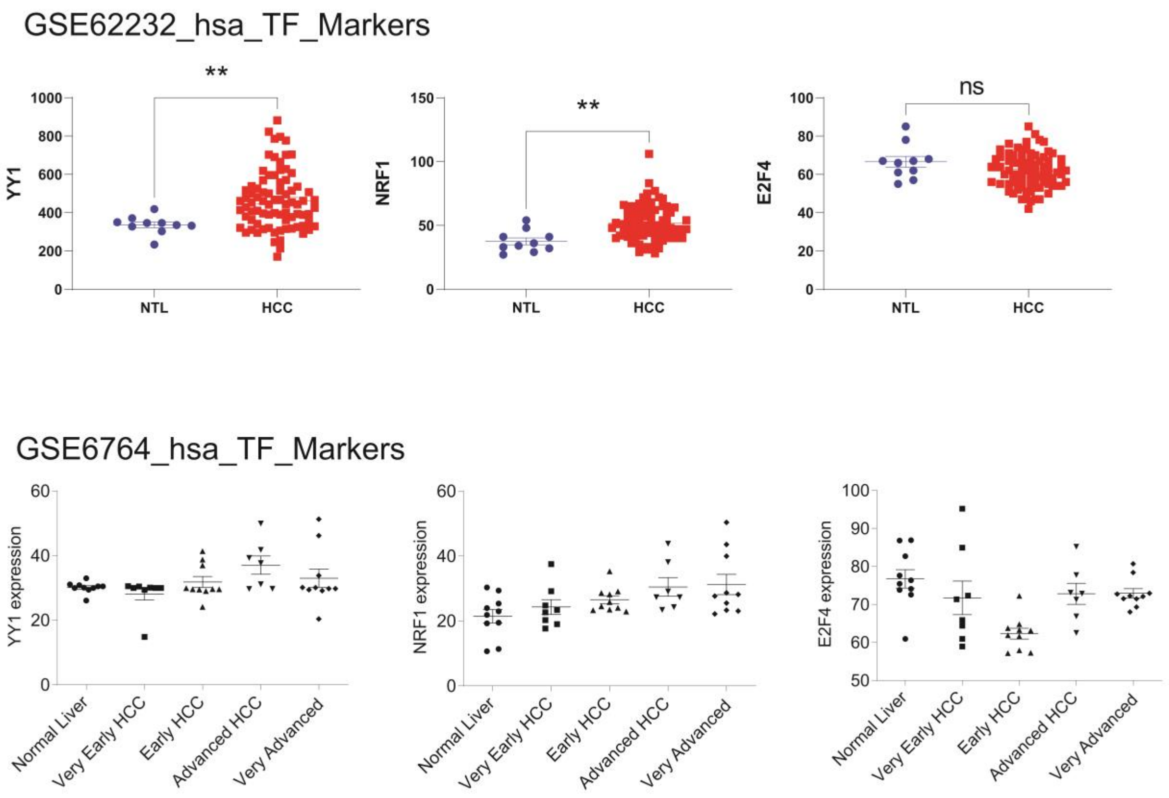


**Supplementary Figure S6: Analysis of YY1, E2F4 and NRF1 expression in independent cohorts of HCC patients**

The GEO database was queried and two different cohorts were selected for further analysis. Analysis of the GSE62232 cohort (**top panel**), which contains the comparison between tumor tissue (HCC) and non-tumor tissue (NTL), identified the significant upregulation of YY1, E2F4 and NRF1 in the liver of HCC patients. These data were independently validated for NRF1 and YY1, but not E2F4, via the analysis of the GSE6764 cohort (**lower panel**), which contains the comparison between tumor tissue at different stage. Data are shown as mRNA expressions for individual patients while the plot represent the mean ± SD. Statistical analysis was performed with two-tailed T-test between two samples, or One-way ANOVA for three or more samples, $p \leq 0.05$ was considered significant. n.s. = Not significant; **, $p \leq 0.01$.

A

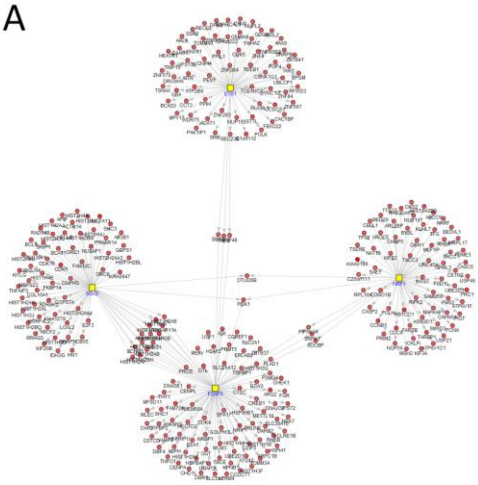

| B | Human Grade III HCC [GEO repository accession no. GSE45050 ] |
|---|---|
| | HCC-tissue *vs.* adjacent non-tumor tissue |
| YY1 | p=6.5x$10^{-2}$, n=67 (9.3%) |
| E2F4 | n.s. |
| FOXP3 | p=9.0x$10^{-5}$, n=98 (11.0%) |
| NRF1 | p=2.0x$10^{-5}$, n=70 (12.5%) |

| C | miR-122 *KO* mice [GEO repository accession no. GSE31453 and GSE27713] | | |
|---|---|---|---|
| | liver miR-122 *KO mice vs* liver wild-type mice (2 months-GSE27713) | HCC-tissue *KO* mice *vs.* healthy liver wild-type (11 months-GSE31453) | HCC-tissue *KO* mice *vs.* healthy liver wild-type (14 months-GSE31453) |
| YY1 | p=5.3x$10^{-5}$, n=18 (2.9%) | p=2.3x$10^{-5}$, n=5 (0.8%) | p=3.9x$10^{-5}$, n=28 (4.5%) |
| E2F4 | n.s. | n.s. | p=7.0x$10^{-2}$, n=10 (5.1%) |
| FOXP3 | p=1.9x$10^{-2}$, n=78 (8.2%) | p=6.8x$10^{-3}$, n=47 (4.5%) | p=3.0x$10^{-4}$, n=113 (11.9%) |
| NRF1 | n.s. | p=9.0x$10^{-2}$, n=10 (2.0%) | n.s. |

**Supplementary Figure S7**

(**A**) Significantly, the application of GO-Elite to the analysis of cohorts patients with grade III HCC (GEO accession number GSE45050) identified a significant enrichment for genes associated to YY1, NRF1 and FOXP3 transcription factors. (**B**) p values and number of genes found to be significantly enriched for YY1, NRF1 and FOXP3 (GSE45050) and (**C**) p values and genes significantly enriched for YY1, NRF1, E2F4 and FOXP3 in the liver of two months old miR-122 Ko mice (GSE27713) and in the livers of mouse model of liver cancer (GSE31453).

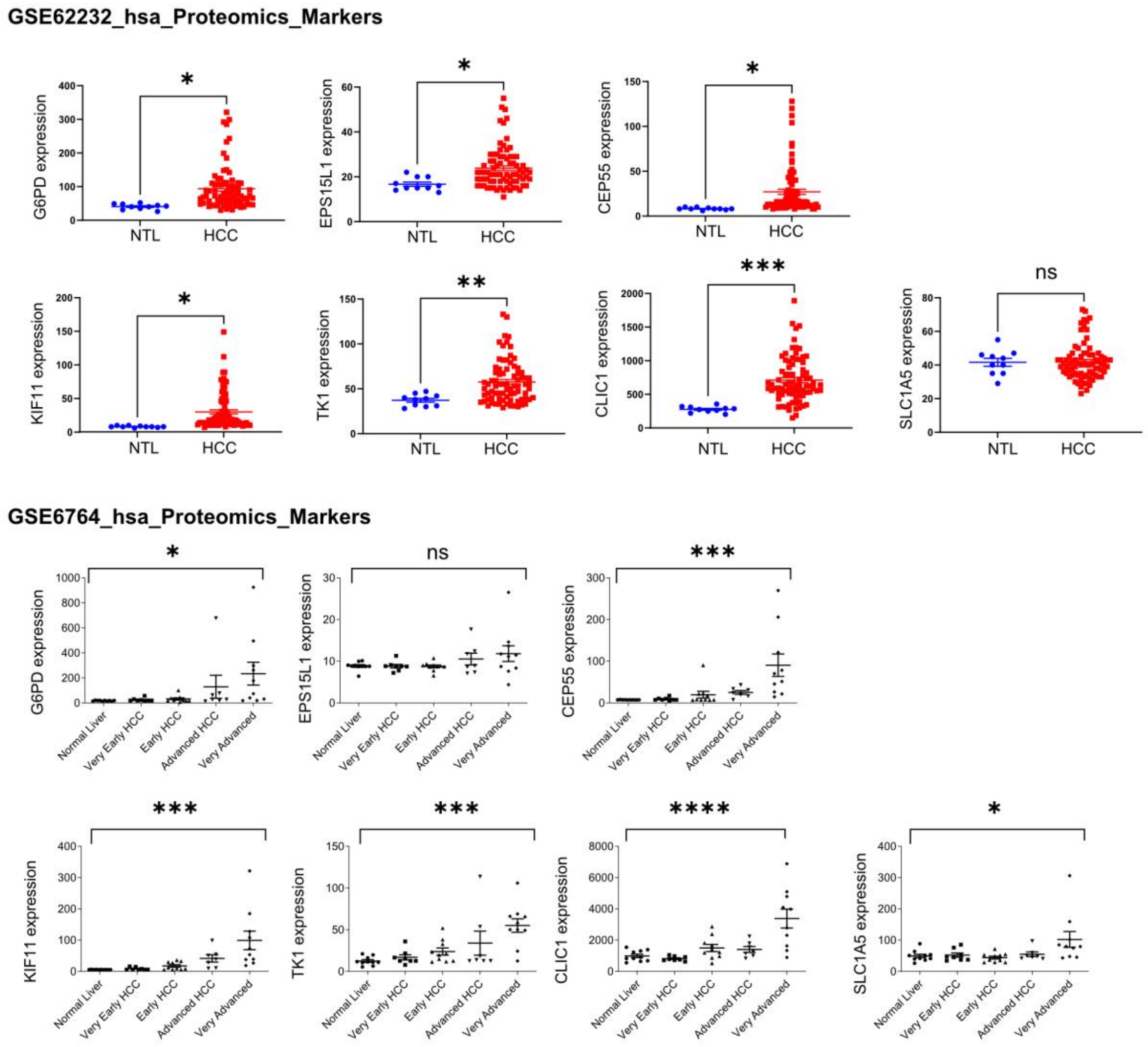


**Supplementary Figure S8**

The GEO database was queried and two different cohorts (GSE62232 and GSE6764) were selected for further analysis. Analysis of the GSE62232 cohort (**top panel**), which contains the comparison between tumor tissue (HCC) and non-tumor tissue (NTL), identified the significant upregulation of G6PD, KIF11, CEP55, TK1, CLIC1 and EPS15L1 in the liver of HCC patients. These data were independently validated via the analysis of the GSE6764 cohort (**lower panel**), which contains the comparison between tumor tissue at different stage. Data are shown as mRNA expressions for individual patients while the plot represent the mean ± SD. Statistical analysis was performed with two-tailed T-test between two samples, or One-way ANOVA for three or more samples, $p \leq 0.05$ was considered significant. ns = Not significant; *, $p \leq 0.05$; **, $p \leq 0.01$; ***, $p \leq 0.001$; ****, $p \leq 0.0001$.

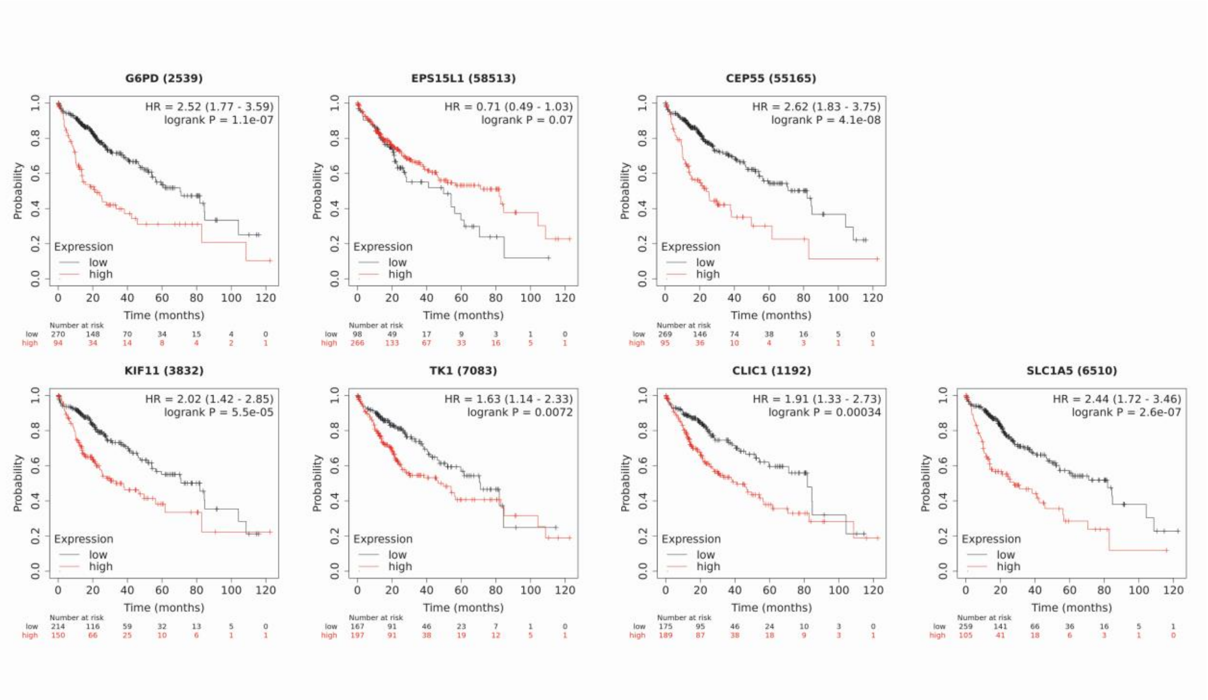


**Supplementary Figure S9 Kaplan-Meier survival curve for miR-122 responsive proteins**

The Kaplan-Meier curve analysis of the liver hepatocellular carcinoma (LIHC) cohorts from TCGA reveals important prognostic factors for liver cancer. Specifically, lower expression levels of CEP55, CLIC1, EPS15L1, G6PD, KIF11, SLC1A5, and TK1 are correlated with higher chances of survival.

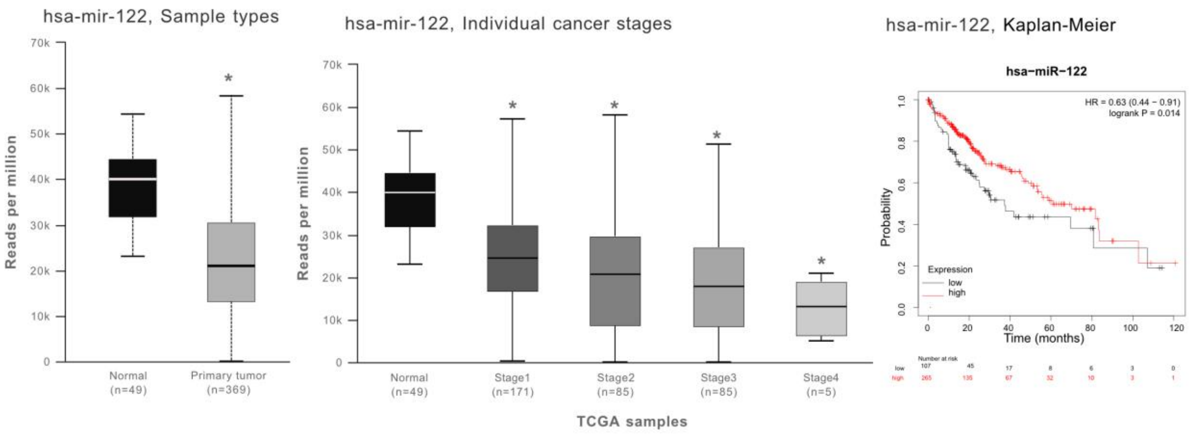


**Supplementary Figure S10: miR-122 is downregulated in liver tumor and it is associated with worst prognosis**

Analysis of LIHC cohorts in TCGA for miRNA expression identified that miR-22 was significantly downregulated in the livers of LIHC patients. The Kaplan-Meier curve analysis of the miR-122 in LIHC cohorts from TCGA indicates that patients with higher miR-122 expression have significantly higher chances of survival.

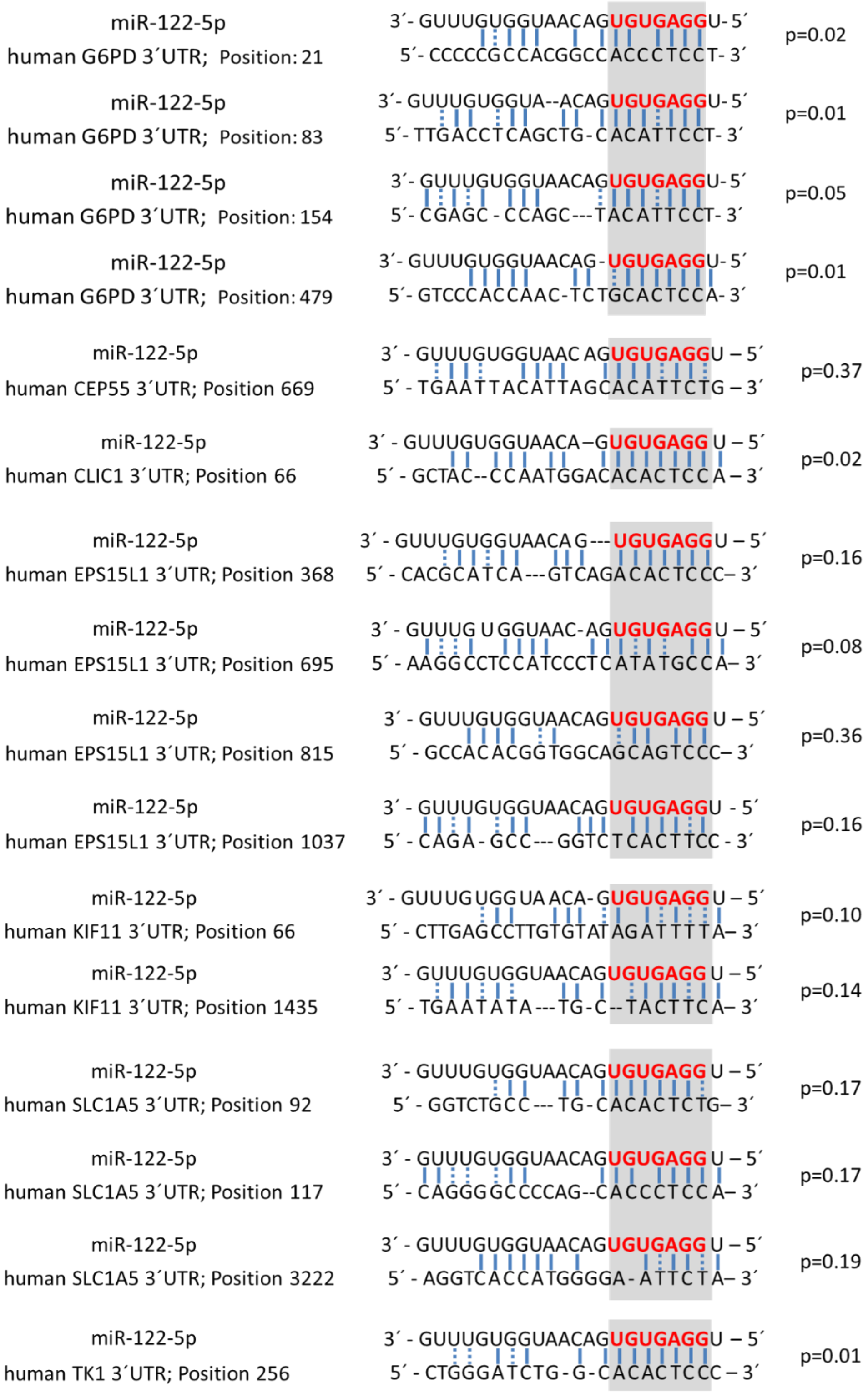


**Supplementary Figure S11: RNA22 predicted miR-122 binding site in the 3’UTR of the GOIs**

*RNA22* predicts miR-122 binding sites in the sequence of the 3´UTRs of CEP55, CLIC1, EPS15L1, G6PD, KIF11, SLC1A5, and TK1. Blue lines and dashed blue lines indicate potential hydrogen bonds between the nucleic acids. The seed sequence of miR 122 is illustrated in red font.
