## Supplementary Figures for "Uncovering novel roles of miR-122 in the pathophysiology of the liver: Potential interaction with NRF1 and E2F4 signaling": Paluschinski_Supplementary_Data_S1.pdf

**Supplementary Materials**

**Isolation and fractionation of polyribosomes**

Polyribosomes were isolated from Huh-7 cells transfected with miR-122 mimic or miR-122 inhibitor 48 h after transfection. To prevent the dissociation of ribosomes from mRNA, 200 µg/mL of cycloheximide (CHX ; Sigma Aldrich, Munich, Germany) was added to the tissue culture medium 10 min before cell lysis. Cells were then washed for 5 min with PBS supplemented with 100 µg/mL CHX (PBS-CHX), collected in 1 mL of fresh PBS-CHX and centrifuged 1000 rpm/4 °C/5 min. The supernatant was removed and the pelleted cells were lysed in 750 µL lysis buffer (5 mM Tris HCl pH 7.5, 1.5 mM KCl, 2.5 mM $MgCl_2$, 0.2 mM CHX), in presence of 120 U/mL RNase Inhibitor (Thermo Fisher Scientific), 120 U/µL DNase I (M0303L, New England Bio Labs, Frankfurt a.M., Germany), 0.5% sodium deoxycholate (Sigma Aldrich) and 0.5% Triton X-100 and passed through a 25G needle. Cell nuclei were removed by centrifugation at 16.000 rpm/4 °C/10 min, and 50 µL of the supernatant was used for total RNA analysis, whereas the remaining supernatant was layered on top of a 10 mL linear sucrose gradient (10 - 50% sucrose (w/v) in 5 mM Tris HCl pH 7.5, 1.5 mM KCl, 2.5 mM MgCl2, 0.2 mM CHX). The samples were centrifuged in a SW 41 Ti rotor (Beckam Coulter, Krefeld, Germany) at 33.500 rpm/4 °C/3 h with brake off to separate polyribosomes of different densities. Fractions of 700 µL were collected by eluting with 60% sucrose solution (supplemented with 0.01% bromophenol blue). RNAs were recovered from each fraction by extracting with an equal volume of phenol-chloroform-isoamyl alcohol (25:24:1, v/v/v; Sigma Aldrich) and precipitating with ethanol. Pelleted RNA was washed twice with 70% ethanol and resolved in RNase-free water.

**cDNA synthesis for miRNA or mRNA and analysis by quantitative real-time PCR (qPCR)**

cDNA synthesis and primer design for the analysis of miRNA expression profiling was carried out by means of miQPCR as previously described. qPCR assays were performed on a ViiA™7 Real-Time PCR System (Thermo Fisher Scientific) with GoTaq qPCR Master Mix (Promega, Mannheim, Germany). cDNA for mRNA analysis was carried out with 200 ng of total RNA using Prime Script (Takara-Clontech, Saint-Germain-en-Laye, France) reverse transcriptase and random priming technique (Thermo Fisher Scientific) according the manufacturer´s instructions. Primers used for miRNA and mRNA quantification are listed in supplementary tables 1 and 2. qPCR data were analyzed by qBase software v.1.3.5 using either DEDD, HPRT1 or β-Actin as reference gene. The selection of appropriate reference genes was made based on GeNorm algorithm to find the most stable gene in a given set of experiment. Statistical analysis was carried out with a significance level of $p<0.05$ by applying unpaired student’s t-test for comparison of two groups, or using one-way ANOVA when three or more groups were compared.

**Human samples and qPCR from HCC tissue**

Human HCC tissue were provided by the Institute of Pathology University Hospital RWTH Aachen. The ethic committee of the Medical Faculty of RWTH Aachen gave the approval (Ethical vote EK122/16) for using the samples. All patients involved in this study were informed and provided their consent to participate.

Formalin-fixed paraffin embedded HCC tissue sections were dewaxed in xylene (50 °C/3 min, twice), washed with 100% ethanol and dried for 10 min at room temperature (RT). The tissue was digested using 100 µg/mL proteinase K in lysis buffer (50 mM Tris-HCl pH 7.4, 10mM EDTA pH 8.0, 0.5% SDS) for 2 h at 56 °C. Thereafter, samples were treated with *DNase I* (AS1260, Promega) in DNase buffer (10mM Tris-HCl pH 7.5, 2.5 mM $MgCl_2$, 0.5mM $CaCl_2$) for 15 min/RT. Total RNA was extracted with Qiazol and chloroform (5:1 (v/v)) and RNA purified using miRNeasy kit (Qiagen, Hilden, Germany). cDNA synthesis was carried out using a gene-specific approach with Prime Script$^{TM}$ RTase (Takara Clontech). Primers used for miRNA and mRNA quantification are listed in supplementary table 1 and 2, respectively. qPCR data were analyzed by qBase software v.1.3.5 (238). Statistical analysis was carried out by unpaired student’s t-test with a significance level of $p<0.05$.

**Sample preparation for mass spectometry**

miR-122 mimic, miR-122 antagomiRs and scrambled oligo control transfected Huh-7 cells were harvested 48 h after transfection. After washing with PBS, cells were scraped from one 10 cm plate per condition (4-5 replicates per condition), centrifuged and lysed with buffer consisting of 30 mM Tris base (Sigma Aldrich, Cat. #T1503), 7 M urea (Sigma Aldrich, Cat. #U1250) and 2 M thiourea (Sigma Aldrich, Cat. #33717). Cells were disrupted by high-speed shaking using a TissueLyser (Qiagen) and sonication for 6 x 10 s. Cell lysate was centrifuged for 15 min at 16,000 g and protein content of the supernatant was determined by Pierce™ 660 nm Protein Assay (Thermo Fisher Scientific). After short SDS-gel-electrophoresis of 5 µg total protein for each sample (~ 5 mm running distance, 10 min) and silver staining (242), the resulting lane was cut out and decolorized with a 1:1 mix of 30 mM sodium thiosulfate (Fluka) and 100 mM potassium hexacyanoferrate (III) (Merck kGaA) washed, reduced with 10 mM dithiothreitol (Serva, Heidelberg, Germany) and alkylated with 55 mM iodacetamide (Sigma Aldrich). Proteins were digested with 2 µg trypsin (Serva) overnight at 37°C. Peptides were extracted with 50% acetonitrile (Sigma Aldrich) and 0.05% trifluoroacetic acid (Fluka).

**Liquid chromatography-tandem mass spectrometry (LC-MS/MS)**

Extracted peptides were separated using a Ultimate 3000 RSCLnano System (Thermo Fisher Scientific) with a Acclaim PepMap100 trap column (3 µm C18 particle size, 100 Å pore size, 75 µm inner diameter, 2 cm length, Thermo Scientific) as a precolumn using 0.1% TFA as a mobile phase and a Acclaim PepMapRSLC (2 µm C18 particle size, 100 Å pore size, 75 µm inner diameter, 25 cm length, Thermo Fisher Scientific) as analytical column with a constant flow rate of 300 nL/min using a 2 h gradient of 0.1% FA (Fluka) to 0.1% FA / 60% acetonitrile. Separated peptides were eluted via nano electrospray ionization into the mass spectrometer (QExactive plus hybrid quadrupole-orbitrap mass spectrometer, Thermo Fisher Scientific). MS spectra were recorded in positive ion mode within a mass range of 300–2000 m/z and a resolution of 70,000. Up to ten precursors (+2, +3 charge states) were isolated within a 2 m/z isolation window and fragmented via higher-energy collisional dissociation. MS/MS spectra were recorded in centroid mode with a maximal ion time of 60 ms and a target value for the automatic gain control set to 100.000. The resolution was 17,500 at an available scan range of 200 to 2,000 m/z. Already fragmented precursors were excluded from further isolation for the next 100 s.

**MS data analysis**

For protein identification, the Proteome Discoverer (Version 1.4, Thermo Fisher Scientific http://www.thermoscientific.com/en/product/proteome-discoverer-software.html) and Mascot search engine were used. MS/MS spectra were searched against the UniProtKB/Swiss-Prot database (version from 2016/02, total entries: 550.552) with the following search parameters: Mass tolerance of 10 ppm precursor mass and 10 mmu (fragment masses), enzyme specificity was trypsin, two missed cleavage sites were considered during the search against human database. Carbamidomethylation of cysteine was set as fixed modification. Oxidation of methionine was accepted as variable modification. For peptide and protein acceptance, we applied the Percolator node and considered a false discovery rate (FDR) of < 1%. Considered proteins showed at least 2 unique peptides with a Mascot score >20. Label-free relative quantification was performed with Progenesis QI for proteomics 2.0 (Nonlinear Dynamics, Newcastle upon Tyne, http://www.nonlinear.com/progenesis/qi-for-proteomics/v2.0/faq/how-does-hi-n-work.aspx for further details). For quantification, only non-conflicting peptides were considered. Automatic alignment of runs to reference run was at least > 80%. For filtering, peak picking limits were set to automatic and the maximum charge was set to 3. Normalization factor was ≤ 2. Peptides with a score < 20 and a mass error >10 ppm were excluded from quantification. R was used for statistical analysis calculating differences between samples by ANOVA (FDR corrected, q<0.05) and Tukey´s post-hoc (p<0.05) test.

**Online data bases**

miRNA sequences were acquired from www.mirbase.org (Version21, release June 2014). Melting temperatures for miRNA and mRNA primers were calculated by use of the online tool OligoAnalyzer 3.1 at https://eu.idtdna.com/calc/analyzer. Primers form RNA amplification were designed with respect to their secondary structure and the potential occurrence of homo- or heterodimers. MiRWalk (http://www.umm.uni-heidelberg.de/apps/zmf/mirwalk/) and RNA22 version 2.0 (https://cm.jefferson.edu/rna22/) were utilized to obtain target prediction data sets of human miR-122. Genomic DNA sequences of the human *MIR-122* gene and its promoter regions were downloaded from Ensembl Genome Browser (release GRCh38.p3 at http://www.ensembl.org/index.html). Venn diagrams and intersection between different gene lists were generated with the online tool Venny (available at http://www.bioinfogp.cnb.csic.es/tools/venny).
