## Supplementary Figures for "Uncovering novel roles of miR-122 in the pathophysiology of the liver: Potential interaction with NRF1 and E2F4 signaling": Paluschinski_Supplementary_Data_S2.pdf

**Supplementary Data S2A:** Literature-mining research for a link between miR-122 responsive genes identified by polyribosome analysis and key words such as `inflammation´, `cancer´, `viral infection´, `liver disease´, or `cytokine signaling´

| ENSEMBL ID | Gene | Gene Name | Associated TF | Keywort | Reference |
|---|---|---|---|---|---|
| ENSG00000085224 | **ATRX** | Alpha Thalassemia/Mental Retardation Syndrome X-Linked | YY1 | Carcinogenesis | (1–7) |
| ENSG00000107262 | **BAG1** | BCL2-Associated Athanogene | YY1 | Carcinogenesis<br>HCC | (8) |
| ENSG00000087088 | **BAX** | BCL2-Associated X Protein | YY1 | HCC<br>NASH<br>HCV | (9–12) |
| ENSG00000180329 | **CCDC43** | Coiled-Coil Domain Containing 43 | YY1 | Viral Infections | (13) |
| ENSG00000196776 | **CD47** | CD47 Molecule | FOXP3 | Tumorigenity<br>HCC | (14–17) |
| ENSG00000153551 | **CMTM7** | CKLF-Like MARVEL Transmembrane Domain Containing 7 | FOXP3 | Cytokine Signaling<br>Cancer<br>Cancer Cells | (18–20) |
| ENSG00000158796 | **DEDD** | Death Effector Domain Containing | YY1 | Cancer metastasis<br>TGFb Signalling | (21,22) |
| ENSG00000150764 | **DIXDC1** | DIX Domain Containing 1 | FOXP3 | Cancer cell invasion and metastasis<br>Cancer cell Proliferation<br>HCC | (23–26) |
| ENSG00000046604 | **DSG2** | Desmoglein 2 | FOXP3 | HCC<br>HBV<br>Cytokine Signaling<br>Cancer | (27–33) |
| ENSG00000102189 | **EEA1** | Early Endosome Antigen 1 | FOXP3 | Viral Infection<br>HCV | (34) |
| ENSG00000164220 | **F2RL2** | Coagulation Factor II (Thrombin) Receptor-Like 2 | FOXP3 | cancer Cell Proliferation<br>Cancer | (35–37) |

**Supplementary Data S2A (continued)**

| ENSEMBL ID | Gene | Gene Name | Associated TF | Keywort | Reference |
|---|---|---|---|---|---|
| ENSG00000088832 | **FKBP1A** | FK506 Binding Protein 1A, 12kDa | FOXP3 | TGFb Signaling<br>HCV<br>HBV<br>HCC | (38–42) |
| ENSG00000138757 | **G3BP2** | GTPase Activating Protein (SH3 Domain) Binding Protein 2 | NRF1 | Cancer cell migration<br>Viral Replication<br>Viral Infection<br>Cytokine Signalling | (43–48) |
| ENSG00000167110 | **GOLGA2** | Golgin A2 | NRF1 | Cancer cell invasion<br>Viral Replication | (49,50) |
| ENSG00000172534 | **HCFC1** | Host Cell Factor C1 | YY1 | Viral Infection<br>Cancer | (51,52) |
| ENSG00000115541 | **HSPE1** | Heat Shock 10kDa Protein 1 | NRF1 | Cancer<br>HCC<br>Inflammation | (53–56) |
| ENSG00000162434 | **JAK1** | Janus Kinase 1 | NRF1 | HCC<br>HBV<br>Cytokine Signalling | (57–60) |
| ENSG00000054523 | **KIF1B** | Kinesin Family Member 1B | E2F4 | HCC<br>HBV | (61–65) |
| ENSG00000131437 | **KIF3A** | Kinesin Family Member 3A | NRF1 | Cancer<br>HCV | (66–69) |
| ENSG00000025800 | **KPNA6** | Karyopherin Alpha 6 (Importin Alpha 7) | YY1 | Cytokine Signaling<br>HBV | (70–73) |
| ENSG00000108424 | **KPNB1** | Karyopherin (Importin) Beta 1 | FOXP3 | Cancer<br>HCV<br>Cytokine Signaling<br>Inflammation | (74–81) |

**Supplementary Data S2A (continued)**

| ENSEMBL ID | Gene | Gene Name | Associated TF | Keywort | Reference |
|---|---|---|---|---|---|
| ENSG00000141503 | **MINK1** | Misshapen-Like Kinase 1 | FOXP3 | Cytokine Signaling<br>Inflammation<br>Cancer<br>Viral Infection | (82–85) |
| ENSG00000132182 | **NUP210** | Nucleoporin 210kDa | FOXP3 | PBC<br>HBV | (86–91) |
| ENSG00000122884 | **P4HA1** | Prolyl 4-Hydroxylase, Alpha Polypeptide I | NRF1 | miR122 target in HSC<br>HCV<br>HCC<br>Fibrosis | (92–95) |
| ENSG00000277258 | **PCGF2** | Polycomb Group Ring Finger 2 | FOXP3 | Cancer<br>Cytokine Signaling | (96–99) |
| ENSG00000071994 | **PDCD2** | Programmed Cell Death 2 | FOXP3 | Cytokine Signaling<br>Cancer | (100,101) |
| ENSG00000150593 | **PDCD4** | Programmed Cell Death 4 (Neoplastic Transformation Inhibitor) | NRF1 | Cytokine Signaling<br>Fibrosis<br>HBV<br>HCC | (102–106) |
| ENSG00000100142 | **POLR2F** | Polymerase (RNA) II (DNA Directed) Polypeptide F | NRF1 | Cancer<br>Viral Infections | (107–109) |
| ENSG00000113522 | **RAD50** | RAD50 Homolog (S. Cerevisiae) | NRF1 | Tumorigenesis<br>Cirrhosis<br>HCC | (110–113) |
| ENSG00000173456 | **RNF26** | Ring Finger Protein 26 | YY1 | Cancer cell<br>Viral Infection | (114,115) |
| ENSG00000101654 | **RNMT** | RNA (Guanine-7-) Methyltransferase | FOXP3 | HCC | (116) |
| ENSG00000134318 | **ROCK2** | Rho-Associated, Coiled-Coil Containing Protein Kinase 2 | FOXP3 | HCC<br>CCC Tumorigenesis<br>Cirrhosis<br>HCV | (34,117–122) |

**Supplementary Data S2A (continued)**

| ENSEMBL ID | Gene | Gene Name | Associated TF | Keywort | Reference |
|---|---|---|---|---|---|
| ENSG00000101665 | **SMAD7** | SMAD Family Member 7 | FOXP3 | TGF Signaling<br>Carcinogenesis<br>HCC<br>HBV | (123–128) |
| ENSG00000084070 | **SMAP2** | Small ArfGAP2 | FOXP3 | HCV<br>HBV | (41,129) |
| ENSG00000198952 | **SMG5** | SMG5 Nonsense Mediated MRNA Decay Factor | YY1 | Viral Infection | (130) |
| ENSG00000198369 | **SPRED2** | Sprouty-Related, EVH1 Domain Containing 2 | FOXP3 | Cancer cell Survival<br>TGFb Signaling | (131–135) |
| ENSG00000166888 | **STAT6** | Signal Transducer And Activator Of Transcription 6, Interleukin-4 Induced | FOXP3 | Cytokine Signaling<br>HCC<br>Inflammation<br>Viral Infection | (136–140) |
| ENSG00000065491 | **TBC1D22B** | TBC1 Domain Family, Member 22B | YY1 | Viral Infection<br>HCV | (141,142) |
| ENSG00000083312 | **TNPO1** | Transportin 1 | FOXP3 | Chemokine Signaling<br>HCV<br>HCC | (77,143–146) |
| ENSG00000063244 | **U2AF2** | U2 Small Nuclear RNA Auxiliary Factor 2 | NRF1 | Cancer Cells<br>HCC<br>Steatosis<br>Viral Infection | (147–150) |
| ENSG00000120942 | **UBIAD1** | UbiA Prenyltransferase Domain Containing 1 | YY1 | Cancer<br>Cholesterol/Lipid metabolism<br>(HCC) | (151–154) |
| ENSG00000156467 | **UQCRB** | Ubiquinol-Cytochrome C Reductase Binding Protein | YY1 | HCC<br>Cytokine Signaling | (155–158) |
| ENSG00000126562 | **WNK4** | WNK Lysine Deficient Protein Kinase 4 | FOXP3 | Tumorigenesis | (159,160) |
| ENSG00000176871 | **WSB2** | WD Repeat And SOCS Box Containing 2 | FOXP3, YY1 | Cytokine Signaling<br>Cancer | (161–165) |

**Supplementary Data S2A (continued)**

| ENSEMBL ID | Gene | Gene Name | Associated TF | Keywort | Reference |
|---|---|---|---|---|---|
| ENSG00000079246 | **XRCC5** | X-Ray Repair Complementing Defective Repair In Chinese Hamster Cells 5 (Double-Strand-Break Rejoining) | NRF1 | HBV<br>HCC<br>Viral Infection | (166–170) |

**Supplementary Data S2A, part 2** Subset of miR-122 responsive proteins as identified by mass spectroscopy (FC>1.5, p<0.05) which are associated to liver disease

| UniProt ID | Gene | Gene Name | Associated to Disease | Reference |
|---|---|---|---|---|
| Q53EZ4 | **CEP55** | Centrosomal Protein 55 | HCC | (171) |
| P52732 | **KIF11** | Kinesin Family Member 11 | HCC | (67,172–174) |
| Q15758 | **SLC1A5** | Solute Carrier Family 1 Member 5 | HCC | (175,176) |
| P11413 | **G6PD** | Glucose-6-Phosphate Dehydrogenase | HCC, HBV | (177–181) |
| Q9UBC2 | **EPS15L1** | Epidermal Growth Factor Receptor Pathway Substrate 15 Like 1 | HCC | (182) |
| Q01650 | **SLC7A5** | Solute Carrier Family 7 Member 5 | HCC (cannine) liver metastasis | (176,183–188) |
| Q8WUP2 | **FBLIM1** | Filamin Binding LIM Protein 1 | HCC metastasis | (189) |
| P14618 | **PKM** | Pyruvate Kinase, Muscle | HCC | (190–193) |
| O14907 | **TAX1BP3** | Tax1 Binding Protein 3 | NAFLD | (194) |

| UniProt ID | Gene | Gene Name | Associated to Disease | Reference |
|---|---|---|---|---|
| P55786 | **NPEPPS** | Aminopeptidase Puromycin Sensitive | HCC | (195) |
| P21291 | **CSRP1** | Cysteine And Glycine Rich Protein 1 | HCC | (196) |
| P46940 | **IQGAP1** | IQ Motif Containing GTPase Activating Protein 1 | HCC | (197,198) |
| P04183 | **TK1** | Thymidine Kinase 1 | HCC | (199,200) |
| P21266 | **GSTM3** | Glutathione S-Transferase Mu 3 | HCC<br>Alcoholic liver disease | (201–203) |
| P07148 | **FABP1** | Fatty Acid Binding Protein 1 | HCC | (204,205) |
| O00299 | **CLIC1** | Chloride Intracellular Channel 1 | HCC | (206–211) |
| P04818 | **TYMS** | Thymidylate Synthetase | HCC | (212–216) |
| P18858 | **LIG1** | DNA Ligase 1 | HCC,<br>HCV | (195,217,218) |
| Q13952 | **NFYC** | Nuclear Transcription Factor Y Subunit Gamma | HBV | (219) |
| Q99523 | **SORT1** | Sortilin 1 | HBV | (220,221) |

### Supplementary Data S2B, part 1: Description of FFPE samples from livers’ of patients included in this study.

| Sample | Sample | Sex | Age | Risk factors | Cirrhosis | Max. tumor size (cm) | No. Nodules | Grading | Vascular invasion |
|---|---|---|---|---|---|---|---|---|---|
| RWTH_01 | 1 | male | 74 | HBV | yes | 4.2 | 1 | 3 | 1 |
| RWTH_02 | 2 | male | 63 | HBV | no | 7.0 | 1 | 3 | 0 |
| RWTH_03 | 3 | male | 62 | HBV | no | 17.5 | multiple | 3 | 1 |
| RWTH_04 | 4 | male | 60 | HBV | yes | 6.7 | 2 | 1 | 0 |
| RWTH_05 | 5 | male | 67 | HBV | yes | 8.0 | 1 | 3 | 0 |
| RWTH_06 | 6 | male | 60 | HBV | yes | 1.3 | 1 | 3 | 1 |
| RWTH_07 | 7 | male | 54 | HBV | yes | 5.9 | 1 | 2 | 1 |
| RWTH_08 | 8 | female | 73 | unknown | no | 7.0 | 1 | 1 | 0 |
| RWTH_09 | 9 | female | 71 | type 2 diabetes | no | 6.0 | 1 | 3 | 1 |
| RWTH_10 | 10 | male | 60 | unknown | no | 4.8 | 1 | 3 | 1 |
| RWTH_11 | 11 | male | 67 | unknown | no | 4.9 | multiple | 3 | 0 |
| RWTH_12 | 12 | female | 75 | C2 | yes | 3.2 | 1 | 3 | 1 |
| RWTH_13 | 13 | male | 63 | type 2 diabetes | yes | 7.5 | 1 | 1 | 0 |
| RWTH_14 | 14 | male | 60 | unknown | yes | 5.5 | 2 | 2 | 0 |
| RWTH_15 | 15 | male | 78 | type 2 diabetes | yes | 3.8 | 3 | 3 | 0 |
| RWTH_16 | 16 | male | 68 | C2 | yes | 4.8 | 1 | 2 | 1 |
| RWTH_17 | 17 | male | 67 | unknown | no | 3.5 | 1 | 2 | 1 |
| RWTH_18 | 18 | male | 86 | unknown | no | 9.0 | 3 | 2 | 1 |
| RWTH_19 | 19 | male | 82 | unknown | no | 12.0 | multiple | 3 | 0 |
| RWTH_20 | 20 | male | 79 | unknown | no | 6.0 | 1 | 2 | 0 |
| RWTH_21 | 21 | male | 65 | unknown | yes | 3.6 | 1 | 3 | 1 |
| RWTH_22 | 22 | male | 52 | unknown | no | 3.0 | 2 | 1 | 0 |
| RWTH_23 | 23 | female | 79 | unknown | no | 5.5 | 1 | 2 | 0 |
| RWTH_24 | 24 | female | 68 | unknown | no | 4.5 | 1 | 3 | 0 |
| RWTH_25 | 25 | male | 63 | NASH | yes | 2.5 | 1 | 2 | 1 |
| RWTH_26 | 26 | male | 61 | unknown | yes | 4.9 | 1 | 2 | 0 |
| RWTH_27 | 27 | male | 77 | PBC | yes | 3.0 | 1 | 2 | 0 |
| RWTH_28 | 28 | male | 74 | type 2 diabetes | no | 2.0 | 1 | 3 | 0 |

### Supplementary Data S2B, part 2: Description of samples included in this study obtained from liver resections of patients with HCC or without HCC

| ID | Pathology | Etiology | Control | Age | Sex | Recurrence | RFS | Survival | OS | AFP | GGT | AST | ALT | GLDH | Bilirubin | Diagnosed |
|---|---|---|---|---|---|---|---|---|---|---|---|---|---|---|---|---|
| UKA-01 | Haemangioma | n.a. | healthy tissue | 37 | female | n.a. | n.a. | n.a. | n.a. | missing | 32 | 24 | 34 | missing | 0.23 | n.a. |
| UKA-02 | Adenoma | n.a. | healthy tissue | 24 | female | n.a. | n.a. | n.a. | n.a. | missing | 63 | missing | 72 | missing | 0.24 | n.a. |
| UKA-03 | FNH | n.a. | healthy tissue | 27 | male | n.a. | n.a. | n.a. | n.a. | missing | 122 | missing | 34 | missing | 0.59 | n.a. |
| UKA-04 | Haemangioma | n.a. | healthy tissue | 42 | female | n.a. | n.a. | n.a. | n.a. | missing | 25 | 20 | 17 | missing | 0.87 | n.a. |
| UKA-05 | FNH | n.a. | healthy tissue | 26 | female | n.a. | n.a. | n.a. | n.a. | missing | 89 | 27 | 44 | missing | 0.23 | n.a. |
| UKA-06 | FNH | n.a. | healthy tissue | 26 | female | n.a. | n.a. | n.a. | n.a. | missing | 140 | 19 | 19 | missing | 0.29 | n.a. |
| UKA-07 | Haemangioma | n.a. | healthy tissue | 48 | female | n.a. | n.a. | n.a. | n.a. | missing | 11 | 54 | 18 | missing | 0.57 | n.a. |
| UKA-08 | HCC | NAFLD | n.a. | 80 | male | yes | 5 | no | 31 | 6 | 53 | 25 | 25 | 0.48 | 13.4 | CT (also confirmed by histology) |
| UKA-09 | HCC | NAFLD+HBV | n.a. | 65 | male | yes | 0 | yes | 1 | 4982 | 242 | 35 | 35 | 0.45 | 13 | CT (also confirmed by histology) |
| UKA-10 | HCC | C2 | n.a. | 72 | male | yes | | yes | 25 | 901 | 129 | 24 | 24 | 1.11 | 13.5 | CT (also confirmed by histology) |
| UKA-11 | HCC | NAFLD | n.a. | 87 | male | yes | 6 | no | 22 | 2 | 22 | 22 | 14 | 0.65 | 16.7 | CT (also confirmed by histology) |
| UKA-12 | HCC | NAFLD | n.a. | 84 | male | yes | 21 | no | 23 | 2 | 316 | 48 | 38 | 0.84 | 14.7 | CT (also confirmed by histology) |
| UKA-13 | HCC | NAFLD | n.a. | 68 | female | yes | 4 | yes | 11 | 20 | 452 | 58 | 39 | 1.85 | 11.8 | CT (also confirmed by histology) |
| UKA-14 | HCC | NAFLD + C2 | n.a. | 78 | male | yes | 7 | no | 21 | 30 | 288 | 43 | 14 | 0.74 | 14.4 | CT (also confirmed by histology) |

**Supplementary Data S2C part 1: Primer sequences used for qPCR amplification of GOIs.**

| Gene | Primer | Sequence 5´- 3´ |
|---|---|---|
| hsa β-Actin | Forward | CAG CAA GCA GGA GTA TGA CG |
| | Reverse | AAA GTC ATG CCA ATC TCA TC |
| hsa BAG1 | Forward | CAT TTG GAG AAG TCT GTG GAG A |
| | Reverse | AAA TCC TTG GGC AGA AAA CC |
| hsa BAX | Forward | AGC AAA CTG GTG CTC AAG G |
| | Reverse | TCT TGG ATC CAG CCC AAC |
| hsa CCDC43 | Forward | GAA GAG GAG AAG CAG AGA AAA GC |
| | Reverse | GCA CCT GAA TCA TCC TTC TCA |
| hsa CD47 | Forward | AGT GAC ACG GTA GCA CCA GTT |
| | Reverse | GAA CAC AGT GCT CTG AGA ACA AG |
| hsa CEP55 | Forward | AGA AGA AGA GAT CCG AAG AGC |
| | Reverse | AGC AGA GAT GTG TAA AGA AAC TG |
| hsa CLIC1 | Forward | CCT GTT GCC AAA GTT ACA CA |
| | Reverse | GTG AAT CCC CGG TAC TTC TT |
| hsa CMTM7 | Forward | TAT CAG CTG GCC CCT GTC |
| | Reverse | CTT GGA AGC TGC CAC AAT G |
| hsa DEDD | Forward | AGC CCT CAG TGA TCC AGA AC |
| | Reverse | GGC AAC ACA CCA CAG GAT AG |
| hsa DIXDC1 | Forward | TTA CGC CCT TCA TGG TCA AT |
| | Reverse | TCC TTC CCG ATC AAT AGC TG |
| hsa DSG2 | Forward | AAT TGC GCT CAT GAT TTT GG |
| | Reverse | GCA ATG GCA CAT CAG CAG TA |
| hsa E2F4 | Forward | GGT ATC GGG CTA ATC GAG AA |
| | Reverse | AAT CTC CCG GGT ATT GCA G |
| hsa EEA1 | Forward | GAA TTG CAA AGA AAG CTG GAT AA |
| | Reverse | TTC AAC GCT TGT GTA TGT TTG A |
| hsa EPS15L | Forward | TTA CCT CGG ATC CAT TCA CG |
| | Reverse | TCA CTG GAT TCA AAG GGG TC |
| hsa F2RL2 | Forward | CTA CGT CCA GGC CAC CTC TA |
| | Reverse | GTG AAG TGG TGG AGG GTA GG |
| hsa G3BP2 | Forward | CCT GTT TCT CTG CCA CAA GA |
| | Reverse | GGA GGC AGG TTT TTA CTG GTC |
| hsa G6PD | Forward | CTG GTG GCC ATG GAG AAG |
| | Reverse | **TGC ATT TCA ACA CCT TGA CCT** |
| hsa Hamp | Forward | CAA CAG ACG GGA CAA CTT GC |
| | Reverse | AGC AGA AAA TGC AGA TGG GGA |
| hsa HCFC1 | Forward | CGC AAT GAG AAG GGC TAT G |
| | Reverse | TGG TGC CAG AGC TGT CTT TA |
| hsa HPRT1 | Forward | AGG TCG CAA GCT TGC TGG |
| | Reverse | **CCA ACA CTT CGT GGG GTC** |

**Supplementary Data S2C part 1 (continued)**

| Gene | Primer | Sequence 5´- 3´ |
|---|---|---|
| hsa KIF1B | Forward | AAG GAC CTT CTT CGT GCT CA |
| | Reverse | GAG GGA CCC CAT GGA TGT A |
| hsa KIF3A | Forward | AAC TTC AAA GGG GAA AGC AAG |
| | Reverse | TTT CAG GCT TTG CAG AAC G |
| hsa KPNA6 | Forward | CTT ATT GTG GCC TCA TAG AGG AA |
| | Reverse | GCC TTC TGG TAG ATC TCC TGG T |
| hsa KPNB1 | Forward | CTG GAA TCG TCC AGG GAT TA |
| | Reverse | TCT GGG TTG TAC CAG CAT CA |
| hsa MINK1 | Forward | AGT TCC TGT GTG AGC GGA AT |
| | Reverse | TGC AGT TAC GGT TCA GAG TCA |
| hsa NRF1 | Forward | CAG TCA CTA TGG CGC TTA ACA |
| | Reverse | ATC TGT CCC CCA CCT TGT AA |
| hsa NUP210 | Forward | GGA CAC AGC CCC CAC TAT T |
| | Reverse | CAT AGG CTG GGC TCC ACA |
| hsa P4HA1 | Forward | AAG ATC TAA CAG GAC TAG ATG TTT CCA |
| | Reverse | TCC TCC AAC TCC ATA ATT TGC |
| hsa PCDC4 | Forward | TGG AAA GCG TAA AGA TAG TGT GTG |
| | Reverse | AAT ATT CTT TCA GCA GCA TAT CAA TC |
| hsa PCGF2 | Forward | TTC TCC GCA ACA AGA TGG AT |
| | Reverse | AGT GGC TCG TCC TCG TAC A |
| hsa PDCD2 | Forward | TGG TGC CAA GAG AAT ATT GGA |
| | Reverse | CCC AGT CTG TCA GCC TTC A |
| hsa POLR2F | Forward | GCG AAT CAC CAC ACC ATA CA |
| | Reverse | ACC ATC ACA GGG GCA CAC |
| hsa pri-miR-122 | Forward | TTT CCT AGA CTG CAG AAT TGA TCA C |
| | Reverse | ATA ATC TGG CCG AAT GAA TGG ATA C |
| hsa RNF26 | Forward | GGC GTT GGG GTT AGT ATC TCT |
| | Reverse | GCC TCA TCA GAC GAT CAC AG |
| hsa RNMT | Forward | TTG GAC CTG GGA TGT GGT |
| | Reverse | GAC AGA AAC ATC GGC AAT ATC A |
| hsa SLC1A5 | Forward | GAT TCG TTC CTG GAT CTT GC |
| | Reverse | GGT AGA GTA TGA GCG AAA GG |
| hsa SLC7A1 | Forward | TCA TCA CCG GCT GGA ACT |
| | Reverse | CCC TCG CTA CGC TTG AAG TA |
| hsa SMAD7 | Forward | AAA CAG GGG GAA CGA ATT ATC |
| | Reverse | ACC ACG CAC CAG TGT GAC |
| hsa SMG5 | Forward | GAT TTG CTG AAG AAG GAA CAC C |
| | Reverse | TTC TGG CAG CGA ATG TAC C |
| hsa SPRED2 | Forward | GAG CAC CGG AGG ATT TAT ACC |
| | Reverse | GAA GCT CAC CTG GCG GTA G |

**Supplementary Data S2C, part 1 (continued)**

| Gene | Primer | Sequence 5´- 3´ |
|---|---|---|
| hsa TBC1D22B | Forward | GAG GCT GAC AGC TTT TGG TG |
| | Reverse | CCT GGT TGT GCA AAG GTG TA |
| hsa TFR2 | Forward | AAG CTG CGG CAG GAG ATC TA |
| | Reverse | GCG ACA CGT ACT GGG AAA GG |
| hsa TK1 | Forward | GTC ATA GGC ATC GAC GAG G |
| | Reverse | GCA GAA CTC CAC GAT GTC A |
| hsa TNPO1 | Forward | TGA TGA TAC AAT TTC TGA CTG GAA TC |
| | Reverse | GGC AGC AGT TCA TCA CGA TA |
| hsa U2AF2 | Forward | CAG GCC TCA CGA CTA CCA G |
| | Reverse | GGG ACC ACA GTG GAC ACA A |
| hsa WSB2 | Forward | TCC TAT GAC CAA TGG GCT TT |
| | Reverse | CGT GGC CAT CTC TTG TCC |
| hsa XRCC5 | Forward | CAA AGA GGA AGC CTC TGG AA |
| | Reverse | AGC TGC TGT GTC TCC ACT TG |
| hsa YY1 | Forward | TGG AGA GAA CTC ACC TCC TGA |
| | Reverse | TCT TTA ATT TTT CTT GGC TTC ATT C |

**Supplementary Data S2C, part 2: Oligonucleotide primer sequences used for miQPCR.**

| miRNA | Primer | Sequence 5´- 3´ |
|---|---|---|
| hsa-miR-122-5p | Forward | GTG ACA ATG GTG TTT GGG |
| hsa-miR-192-5p | Forward | TGA CCT ATG AAT TGA CAG CCG |
| Upm2A | Reverse | CCC AGT TAT GGC CGT TTA |

**Supplementary Data S2D, part1: Primer sequences G6PD 3´UTR into luciferase reporter plasmids.** Primer sequence is given as well as the recognition site for restriction enzymes (RE) used for cloning purposes.

| Gene | Primer | Sequence 5´- 3´ | RE |
|---|---|---|---|
| hsa G6PD 3´UTR | Forward | GAG GTA CCG GGT TTC CAG TAT GAG GGC A | *KpnI* |
| | Reverse | GAG CTA GCT TGC GGA TTT AAT GGC AGG G | *NheI* |

**Supplementary Data S2D, part2: Primer sequences the cloning of the 3’UTR of GOIs into luciferase reporter plasmids by using In Fusing HD.** 3’UTRs sequences are given in **BLACK**, In Fusing Sequences (i.e., overlapping with vector sequences) are given in **RED**.

| Gene | Primer | Sequence 5´- 3´ |
|---|---|---|
| | TK1(+) InFus For | AATCGATAGGTACCGAGCTC**GGGACCTGCGAGGGCCGC** |
| hsa TK1(+) 3'UTR | TK1(+) InFus Rev | TCGAGCCCGGGCTAGC**TGCGTCCACCAACCAGTGAATTTTC** |
| | TK1(-) InFus For | AATCGATAGGTACCGAGCTC**TGCGTCCACCAACCAGTGAATTTTC** |
| hsa TK1(-) 3'UTR | TK1(-) InFus Rev | TCGAGCCCGGGCTAGC**GGGACCTGCGAGGGCCGC** |
| | CLIC1(+) InFus For | AATCGATAGGTACCGAGCTC**GCCCCTCCTGGGACTCCC** |
| hsa CLIC1(+) 3'UTR | CLIC1(+) InFus Rev | TCGAGCCCGGGCTAGC**TTGCGTAAAAACACTTGATTTTT** |
| | CLIC1(-) InFus For | AATCGATAGGTACCGAGCTC**TTGCGTAAAAACACTTGATTTTT** |
| hsa CLIC1(-) 3'UTR | CLIC1(-) InFus Rev | TCGAGCCCGGGCTAGC**GCCCCTCCTGGGACTCCC** |
| | CEP55(+) InFus For | AATCGATAGGTACCGAGCTC**CAAAATAAGTATTTGTTTTG** |
| hsa CEP55(+) 3'UTR | CEP55(+) InFus Rev | TCGAGCCCGGGCTAGC**TTAAAACATTAAATAATTTTATTC** |
| | CEP55(-) InFus For | AATCGATAGGTACCGAGCTC**TTAAAACATTAAATAATTTTATTC** |
| hsa CEP55(-) 3'UTR | CEP55(-) InFus Rev | TCGAGCCCGGGCTAGC**CAAAATAAGTATTTGTTTTG** |
| | KIF11(+) InFus For | AATCGATAGGTACCGAGCTC**TTCACTTGGGGGTTGGCA** |
| hsa KIF11(+) 3'UTR | KIF11(+) InFus Rev | TCGAGCCCGGGCTAGC**TTAATGTAGAAACCACATTTATTAA** |
| | KIF11(-) InFus For | AATCGATAGGTACCGAGCTC**TTAATGTAGAAACCACATTTATTAA** |
| hsa KIF11(-) 3'UTR | KIF11(-) InFus Rev | TCGAGCCCGGGCTAGC**TTCACTTGGGGGTTGGCA** |
| | EPS15L1(+) InFus For | AATCGATAGGTACCGAGCTC**AGGAAAGCAGATGAGGTGTG** |
| hsa EPS15L1(+) 3'UTR | EPS15L1(+) InFus Rev | TCGAGCCCGGGCTAGC**TTTCATTTCCCTTAGCATTTTATTT** |
| | EPS15L1(-) InFus For | AATCGATAGGTACCGAGCTC**TTTCATTTCCCTTAGCATTTTATTT** |
| hsa EPS15L1(-) 3'UTR | EPS15L1(-) InFus Rev | TCGAGCCCGGGCTAGC**AGGAAAGCAGATGAGGTGTG** |
| | SLC1A5(+) InFus For | AATCGATAGGTACCGAGCTC**ACCCCGGGAGGGACCTTC** |
| hsa SLC1A5(+) 3'UTR | SLC1A5(+) InFus Rev | TCGAGCCCGGGCTAGC**TTAAATAGTTGACACTCAATTTTAT** |
| | SLC1A5(-) InFus For | AATCGATAGGTACCGAGCTC**TTAAATAGTTGACACTCAATTTTAT** |
| hsa SLC1A5(-) 3'UTR | SLC1A5(-) InFus Rev | TCGAGCCCGGGCTAGC**ACCCCGGGAGGGACCTTC** |

**Supplementary references**

1. Wood LD, Heaphy CM, Daniel HD-J, Naini B V, Lassman CR, Arroyo MR, et al. Chromophobe hepatocellular carcinoma with abrupt anaplasia: a proposal for a new subtype of hepatocellular carcinoma with unique morphological and molecular features. Mod. Pathol. 2013;26:1586–1593.

2. Amorim JP, Santos G, Vinagre J, Soares P. The role of ATRX in the alternative lengthening of telomeres (ALT) phenotype. Genes (Basel). 2016;7.

3. Hironaka K, Factor VM, Calvisi DF, Conner EA, Thorgeirsson SS. Dysregulation of DNA repair pathways in a transforming growth factor α/c-myc transgenic mouse model of accelerated hepatocarcinogenesis. Lab. Investig. 2003;83:643–654.

4. Nowacka-Zawisza M, Bryś M, Romanowicz-Makowska H, Zadrozny M, Kulig A, Małgorzata Krajewska W. Loss of heterozygosity and microsatellite instability at RAD52 and RAD54 loci in breast cancer. Polish J. Pathol. 2006;57:83–89.

5. Qadeer ZA, Harcharik S, Valle-Garcia D, Chen C, Birge MB, Vardabasso C, et al. Decreased Expression of the Chromatin Remodeler ATRX Associates With Melanoma Progression. J. Invest. Dermatol. 2014;134:1768–1772.

6. Heaphy CM, de Wilde RF, Jiao Y, Klein AP, Edil BH, Shi C, et al. Altered Telomeres in Tumors with ATRX and DAXX Mutations Christopher. Science (80-. ). 2011;333:425.

7. Lukashchuk V, McFarlane S, Everett RD, Preston CM. Human Cytomegalovirus Protein pp71 Displaces the Chromatin-Associated Factor ATRX from Nuclear Domain 10 at Early Stages of Infection. J. Virol. 2008;82:12543–12554.

8. Ni W, Chen B, Zhou G, Lu C, Xiao M, Guan C, et al. Overexpressed nuclear BAG-1 in human hepatocellular carcinoma is associated with poor prognosis and resistance to doxorubicin. J. Cell. Biochem. 2013;114:2120–2130.

9. Garcia EJ, Lawson D, Cotsonis G, Cohen C. Hepatocellular carcinoma and markers of apoptosis (bcl-2, bax, bcl-x): Prognostic significance. Appl. Immunohistochem. Mol. Morphol. 2002;10:210–217.

10. Zheng JY, Yang GS, Wang WZ, Li J, Li KZ, Guan WX, et al. Overexpression of Bax induces apoptosis and enhances drug sensitivity of hepatocellular cancer-9204 cells. World J. Gastroenterol. 2005;11:3498–3503.

11. Li CP, Li JH, He SY, Li P, Zhong XL. Roles of Fas/Fasl, Bcl-2/Bax, and Caspase-8 in rat nonalcoholic fatty liver disease pathogenesis. Genet. Mol. Res. 2014;13:3991–3999.

12. Deng L, Adachi T, Kitayama K, Bungyoku Y, Kitazawa S, Ishido S, et al. Hepatitis C Virus Infection Induces Apoptosis through a Bax-Triggered, Mitochondrion-Mediated, Caspase 3-Dependent Pathway. J. Virol. 2008;82:10375–10385.

13. Impens F, Timmerman E, Staes A, Moens K, Ariën KK, Verhasselt B, et al. A catalogue of putative HIV-1 protease host cell substrates. Biol. Chem. 2012;393:915–931.

14. Xiaoa Z, Chung H, Babak Banan, Manning PT, Ott KC, Lin S, et al. Antibody mediated therapy targeting CD47 inhibits tumor progression of hepatocellular carcinoma. Cancer Lett. 2015;360:302–309.

15. Lo J, Lau EYT, Ching RHH, Cheng BYL, Ma MKF, Ng IOL, et al. Nuclear factor kappa B-mediated CD47 up-regulation promotes sorafenib resistance and its blockade synergizes the effect of sorafenib in hepatocellular carcinoma in mice. Hepatology. 2015;62:534–545.

16. Lee TKW, Cheung VCH, Lu P, Lau EYT, Ma S, Tang KH, et al. Blockade of CD47-mediated cathepsin S/protease-activated receptor 2 signaling provides a therapeutic target for hepatocellular carcinoma. Hepatology. 2014;60:179–191.

17. Roberts DD, Kaur S, Soto-Pantoja DR. Therapeutic targeting of the thrombospondin-1 receptor CD47 to treat liver cancer. J. Cell Commun. Signal. 2015;9:101–102.

18. Liu Q, Yu S, Guan-Chao J, Zu-Li Z, Bao-Cai L, Liang B, et al. Change of CMTM7 expression, a potential tumor suppressor, is associated with poor clinical outcome in human non-small cell lung cancer. Chin. Med. J. (Engl). 2013;126:3006–12.

19. Li H, Li J, Su Y, Fan Y, Guo X, Li L, et al. A novel 3p22.3 gene CMTM7 represses oncogenic EGFR

signaling and inhibits cancer cell growth. Oncogene. 2014;33:3109–3118.

20. Liu B, Su Y, Li T, Yuan W, Mo X, Li H, et al. CMTM7 knockdown increases tumorigenicity of human non-small cell lung cancer cells and EGFR-AKT signaling by reducing Rab5 activation. Oncotarget. 2015;6:41092–41107.

21. Lv Q, Hua F, Hu Z-W. Use of the Tumor Repressor DEDD as a Prognostic Marker of Cancer Metastasis. Methods Mol. Biol. 2014;1165:197–222.

22. Xue JF, Hua F, Lv Q, Lin H, Wang ZY, Yan J, et al. DEDD negatively regulates transforming growth factor-β1 signaling by interacting with Smad3. FEBS Lett. 2010;584:3028–3034.

23. Zhou S, Shen J, Lin S, Liu X, Xu M, Shi L, et al. Downregulated expression of DIXDC1 in hepatocellular carcinoma and its correlation with prognosis. Tumor Biol. 2016;37:13607–13616.

24. Tan C, Qiao F, Wei P, Chi Y, Wang W, Ni S, et al. DIXDC1 activates the Wnt signaling pathway and promotes gastric cancer cell invasion and metastasis. Mol. Carcinog. 2016;55:397–408.

25. Xu Z, Liu D, Fan C, Luan L, Zhang X, Wang E. DIXDC1 increases the invasion and migration ability of non-small-cell lung cancer cells via the PI3K-AKT/AP-1 pathway. Mol. Carcinog. 2014;53:917–925.

26. Wang L, Cao XX, Chen Q, Zhu TF, Zhu HG, Zheng L. DIXDC1 targets p21 and cyclin D1 via PI3K pathway activation to promote colon cancer cell proliferation. Cancer Sci. 2009;100:1801–1808.

27. Kim BY, Lee JG, Park S, Ahn JY, Ju YJ, Chung JH, et al. Feature genes of hepatitis B virus-positive hepatocellular carcinoma, established by its molecular discrimination approach using prediction analysis of microarray. Biochim. Biophys. Acta. 2004;1739:50–61.

28. Kaposi-Novak P, Libbrecht L, Woo H, Lee Y, Sears NC, Conner EA, et al. Central Role of c-Myc during Malignant Conversion in Human Hepatocarcinogenesis. Cancer Res. 2009;69:2775–2782.

29. Kamekura R, Kolegraff KN, Nava P, Hilgarth RS, Feng M, Parkos CA, et al. Loss of the desmosomal cadherin desmoglein-2 suppresses colon cancer cell proliferation through EGFR signaling. Oncogene. 2014;33:4531–4536.

30. Gupta A, Nitoiu D, Brennan-Crispi D, Addya S, Riobo NA, Kelsell DP, et al. Cell cycle- and cancer-associated gene networks activated by Dsg2: Evidence of cystatin a deregulation and a potential role in cell-cell adhesion. PLoS One. 2015;10:1–20.

31. Barber AG, Castillo-Martin M, Bonal DM, Rybicki BA, Christiano AM, Cordon-Cardo C. Characterization of desmoglein expression in the normal prostatic gland. Desmoglein 2 is an independent prognostic factor for aggressive prostate cancer. PLoS One. 2014;9.

32. Saaber F, Chen Y, Cui T, Yang L, Mireskandari M, Petersen I. Expression of desmogleins 1-3 and their clinical impacts on human lung cancer. Pathol. Res. Pract. 2015;211:208–213.

33. Fang W-K, Gu W, Liao L-D, Chen B, Wu Z-Y, Wu J-Y, et al. Prognostic significance of desmoglein 2 and desmoglein 3 in esophageal squamous cell carcinoma. Asian Pacific J. Cancer Prev. 2014;15:871–6.

34. Berger KL, Cooper JD, Heaton NS, Yoon R, Oakland TE, Jordan TX, et al. Roles for endocytic trafficking and phosphatidylinositol 4-kinase III alpha in hepatitis C virus replication. Proc. Natl. Acad. Sci. 2009;106:7577–7582.

35. Jan YJ, Ko BS, Liu TA, Wu YM, Liang SM, Chen SC, et al. Expression of partitioning defective 3 (Par-3) for predicting extrahepatic metastasis and survival with hepatocellular carcinoma. Int. J. Mol. Sci. 2013;14:1684–1697.

36. Elste AP, Petersen I. Expression of proteinase-activated receptor 1-4 (PAR 1-4) in human cancer. J. Mol. Histol. 2010;41:89–99.

37. Kaufmann R, Rahn S, Pollrich K, Hertel J, Dittmar Y, Hommann M, et al. Thrombin-mediated hepatocellular carcinoma cell migration: Cooperative action via proteinase-activated receptors 1 and 4. J. Cell. Physiol. 2007;211:699–707.

38. Huse M, Chen YG, Massagué J, Kuriyan J. Crystal structure of the cytoplasmic domain of the type I TGFβ receptor in complex with FKBP12. Cell. 1999;96:425–436.

39. Chen YG, Liu F, Massague J. Mechanism of TGFβ receptor inhibition by FKBP12. EMBO J. 1997;16:3866–76.

40. Colman H, Le Berre-Scoul C, Hernandez C, Pierredon S, Bihouee A, Houlgatte R, et al. Genome-Wide Analysis of Host mRNA Translation during Hepatitis C Virus Infection. J. Virol. 2013;87:6668–6677.

41. Xu Z, Zhai L, Yi T, Gao H, Fan F, Li Y, et al. Hepatitis B virus X induces inflammation and cancer in mice liver through dysregulation of cytoskeletal remodeling and lipid metabolism. Oncotarget. 2016;7.

42. Wang F, Anderson PW, Salem N, Kuang Y, Tennant BC, Lee Z. Gene expression studies of hepatitis virus-induced woodchuck hepatocellular carcinoma in correlation with human results. Int. J. Oncol. 2007;30:33–44.

43. Huang J, Yang J, Lei Y, Gao H, Wei T, Luo L, et al. An ANCCA/PRO2000-miR-520a-E2F2 regulatory loop as a driving force for the development of hepatocellular carcinoma. Oncogenesis. 2016;5:e229.

44. Wei SC, Fattet L, Tsai JH, Guo Y, Pai VH, Majeski HE, et al. Matrix stiffness drives Epithelial-Mesenchymal Transition and tumour metastasis through a TWIST1-G3BP2 mechanotransduction pathway. Nat. Cell Biol. 2015;17:678–688.

45. Schwerk J, Jarret AP, Joslyn RC, Savan R. Landscape of post-transcriptional gene regulation during hepatitis C virus infection. Curr. Opin. Virol. 2015;12:74–84.

46. Bidet K, Dadlani D, Garcia-Blanco MA. G3BP1, G3BP2 and CAPRIN1 Are Required for Translation of Interferon Stimulated mRNAs and Are Targeted by a Dengue Virus Non-coding RNA. PLoS Pathog. 2014;10.

47. Cristea IM, Rozjabek H, Molloy KR, Karki S, White LL, Rice CM, et al. Host Factors Associated with the Sindbis Virus RNA-Dependent RNA Polymerase: Role for G3BP1 and G3BP2 in Virus Replication. J. Virol. 2010;84:6720–6732.

48. Prigent M, Barlat I, Langen H, Dargemont C. IκBα and IκBα/NF-κB complexes are retained in the cytoplasm through interaction with a novel partner, RasGAP SH3-binding protein 2. J. Biol. Chem. 2000;275:36441–36449.

49. Chang S-H, Hong S-H, Jiang H-L, Minai-Tehrani A, Yu K-N, Lee J-H, et al. GOLGA2/GM130, cis-Golgi Matrix Protein, is a Novel Target of Anticancer Gene Therapy. Mol. Ther. 2012;20:2052–2063.

50. Khadka S, Vangeloff AD, Zhang C, Siddavatam P, Heaton NS, Wang L, et al. A Physical Interaction Network of Dengue Virus and Human Proteins. Mol. Cell. Proteomics. 2011;10:M111.012187.

51. Johnson KM, Mahajan SS, Wilson C. Herpes simplex virus transactivator VP16 discriminates between HCF-1 and a novel family member, HCF-2. J. Virol. 1999;73:3930–40.

52. Parker JB, Palchaudhuri S, Yin H, Wei J, Chakravarti D. A Transcriptional Regulatory Role of the THAP11-HCF-1 Complex in Colon Cancer Cell Function. Mol. Cell. Biol. 2012;32:1654–1670.

53. Chu S, Wen Q, Qing Z, Luo J, Wang W, Chen L, et al. High expression of heat shock protein 10 correlates negatively with estrogen/progesterone receptor status and predicts poor prognosis in invasive ductal breast carcinoma. Hum. Pathol. 2017;61:173–180.

54. Ye Y, Huang A, Huang C, Liu J, Wang B, Lin K, et al. Comparative mitochondrial proteomic analysis of hepatocellular carcinoma from patients. Proteomics - Clin. Appl. 2013;7:403–415.

55. Qi LN, Li LQ, Chen YY, Chen ZH, Bai T, Xiang B De, et al. Genome-wide and differential proteomic analysis of hepatitis B virus and aflatoxin B1 related hepatocellular carcinoma in Guangxi, China. PLoS One. 2013;8:1–15.

56. Jia H, Halilou AI, Hu L, Cai W, Liu J, Huang B. Heat shock protein 10 (Hsp10) in immune-related diseases: One coin, two sides. Int. J. Biochem. Mol. Biol. 2011;2:47–57.

57. Xie H, Bae H, Noh J, Eun J, Kim J, Jung K, et al. Mutational analysis of JAK1 gene in human hepatocellular carcinoma H. Neoplasma. 2009;56:136–140.

58. Yang S, Luo C, Gu Q, Xu Q, Wang G, Sun H, et al. Activating JAK1 mutation may predict the sensitivity of JAK-STAT inhibition in hepatocellular carcinoma. Oncotarget. 2016;7:5461–5469.

59. Lee Y, Yun Y. HBx Protein of Hepatitis B Virus Activates Jak1-STAT Signaling. J. Biol. Chem. 1998;273:25510–25515.

60. Arbuthnot P, Capovilla A, Kew M. Putative role of hepatitis B virus X protein in hepatocarcinogenesis: effects on apoptosis, DNA repair, mitogen-activated protein kinase and JAK/STAT pathways. J. Gastroenterol. Hepatol. 2000;15:357–68.

61. Wang ZC, Gao Q, Shi JY, Yang LX, Zhou J, Wang XY, et al. Genetic Polymorphism of the Kinesin-Like Protein KIF1B Gene and the Risk of Hepatocellular Carcinoma. PLoS One. 2013;8:1–6.

62. Yang SZ, Wang JT, Yu WW, Liu Q, Wu YF, Chen SG. Downregulation of KIF1B mRNA in hepatocellular carcinoma tissues correlates with poor prognosis. World J. Gastroenterol. 2015;21:8418–8424.

63. Al-Qahtani A, Al-Anazi M, Viswan NA, Khalaf N, Abdo AA, Sanai FM, et al. Role of Single Nucleotide Polymorphisms of KIF1B Gene in HBV-Associated Viral Hepatitis. PLoS One. 2012;7:5–10.

64. Chen J, Wang Y, Lv W, Gan Y, Chang W, Tian N-N, et al. Effects of interactions between environmental factors and KIF1B genetic variants on the risk of hepatocellular carcinoma in a Chinese cohort. World J. Gastroenterol. 2016;22:4183.

65. Zhong R, Tian Y, Liu L, Qiu Q, Wang Y, Rui R, et al. HBV-related hepatocellular carcinoma susceptibility gene KIF1B is not associated with development of chronic hepatitis B. PLoS One. 2012;7:3–6.

66. Liu Z, Rebowe RE, Wang Z, Li Y, Wang Z, DePaolo JS, et al. KIF3a Promotes Proliferation and Invasion via Wnt Signaling in Advanced Prostate Cancer. Mol. Cancer Res. 2014;12:491–503.

67. Yu Y, Feng YM. The role of kinesin family proteins in tumorigenesis and progression. Cancer. 2010;116:5150–5160.

68. Giugliano S, Kriss M, Golden-Mason L, Dobrinskikh E, Stone AEL, Soto-Gutierrez A, et al. HCV Infection Induces Autocrine Interferon Signaling by Human Liver Endothelial Cell and Release of Exosomes, Which Inhibits Viral Replication. Gastroenterology. 2015;148:392–402.

69. Gaudin R, de Alencar BC, Jouve M, Bèrre S, Le Bouder E, Schindler M, et al. Critical role for the kinesin KIF3A in the HIV life cycle in primary human macrophages. J. Cell Biol. 2012;199:467–479.

70. Ma J, Cao X. Regulation of Stat3 nuclear import by importin α5 and importin α7 via two different functional sequence elements. Cell. Signal. 2006;18:1117–1126.

71. Kann M, Schmitz A, Rabe B. Intracellular transport of hepatitis B virus. World J. Gastroenterol. 2007;13:39–47.

72. Laurila E, Vuorinen E, Savinainen K, Rauhala H, Kallioniemi A. KPNA7, a nuclear transport receptor, promotes malignant properties of pancreatic cancer cells in vitro. Exp. Cell Res. 2014;322:159–167.

73. Sun Z, Wu T, Zhao F, Lau A, Birch CM, Zhang DD. KPNA6 (Importin α-7)-Mediated Nuclear Import of Keap1 Represses the Nrf2-Dependent Antioxidant Response. Mol. Cell. Biol. 2011;31:1800–1811.

74. Yang L, Hu B, Zhang Y, Qiang S, Cai J, Huang W, et al. Suppression of the nuclear transporter-KPNβ1 expression inhibits tumor proliferation in hepatocellular carcinoma. Med. Oncol. 2015;32.

75. Liang P, Zhang H, Wang G, Li S, Cong S, Luo Y, et al. KPNB1, XPO7 and IPO8 Mediate the Translocation of NF-κB/p65 in to the Nucleus. Traffic. 2013;14:1132–1143.

76. van der Watt PJ, Ngarande E, Leaner VD. Overexpression of Kpnβ1 and Kpnα2 importin proteins in cancer derives from deregulated E2F activity. PLoS One. 2011;6:1–10.

77. Germain M-A, Chatel-Chaix L, Gagné B, Bonneil É, Thibault P, Pradezynski F, et al. Elucidating Novel Hepatitis C Virus–Host Interactions Using Combined Mass Spectrometry and Functional Genomics Approaches. Mol. Cell. Proteomics. 2014;13:184–203.

78. Gagné B, Tremblay N, Park AY, Baril M, Lamarre D. Importin β1 targeting by hepatitis C virus NS3/4A protein restricts IRF3 and NF-κB signaling of IFNB1 antiviral response. Traffic. 2017;18:362–377.

79. Nishitsuji H, Ujino S, Shimizu Y, Harada K, Zhang J, Sugiyama M, et al. Novel reporter system to

monitor early stages of the hepatitis B virus life cycle. Cancer Sci. 2015;106:1616–1624.

80. Kojima Y, Nakayama M, Nishina T, Nakano H, Koyanagi M, Takeda K, et al. Importin β1 protein-mediated nuclear localization of Death Receptor 5 (DR5) limits DR5/Tumor Necrosis Factor (TNF)-related apoptosis-inducing ligand (TRAIL)-induced cell death of human tumor cells. J. Biol. Chem. 2011;286:43383–43393.

81. Stelma T, Leaner VD. KPNB1-mediated nuclear import is required for motility and inflammatory transcription factor activity in cervical cancer cells. Oncotarget. 2017;8:32833–32847.

82. Daulat AM, Luu O, Sing A, Zhang L, Wrana JL, McNeill H, et al. Mink1 Regulates -Catenin-Independent Wnt Signaling via Prickle Phosphorylation. Mol. Cell. Biol. 2012;32:173–185.

83. Kaneko S, Chen X, Lu P, Yao X, Wright TG, Rajurkar M, et al. Smad inhibition by the Ste20 kinase Misshapen. Proc. Natl. Acad. Sci. 2011;108:11127–11132.

84. Leong SY, Ong BKT, Chu JJH. The Role of Misshapen NCK-related kinase (MINK), a Novel Ste20 Family Kinase, in the IRES-Mediated Protein Translation of Human Enterovirus 71. PLoS Pathog. 2015;11:1–33.

85. Esau C, Davis S, Murray SF, Yu XX, Pandey SK, Pear M, et al. miR-122 regulation of lipid metabolism revealed by in vivo antisense targeting. Cell Metab. 2006;3:87–98.

86. Yamagiwa S, Kamimura H, Takamura M, Aoyagi Y. Autoantibodies in primary biliary cirrhosis: Recent progress in research on the pathogenetic and clinical significance. World J. Gastroenterol. 2014;20:2606–2612.

87. Luettig B, Boeker KHW, Schoessler W, Will H, Loges S, Schmidt E, et al. The antinuclear autoantibodies Sp100 and gp210 persist after orthotopic liver transplantation in patients with primary biliary cirrhosis. J. Hepatol. 1998;28:824–828.

88. Nakamura M, Kondo H, Mori T, Komori A, Matsuyama M, Ito M, et al. Anti-gp210 and anti-centromere antibodies are different risk factors for the progression of primary biliary cirrhosis. Hepatology. 2007;45:118–127.

89. Czaja AJ. Autoantibodies as prognostic markers in autoimmune liver disease. Dig. Dis. Sci. 2010;55:2144–2161.

90. Nakamura M, Takii Y, Ito M, Komori A, Yokoyama T, Shimizu-Yoshida Y, et al. Increased expression of nuclear envelope gp210 antigen in small bile ducts in primary biliary cirrhosis. J. Autoimmun. 2006;26:138–145.

91. Li B-A, Liu J, Hou J, Tang J, Zhang J, Xu J, et al. Autoantibodies in Chinese patients with chronic hepatitis B: Prevalence and clinical associations. World J. Gastroenterol. 2015;21:283–291.

92. Li J, Ghazwani M, Zhang Y, Lu J, Li J, Fan J, et al. miR-122 regulates collagen production via targeting hepatic stellate cells and suppressing P4HA1 expression. J. Hepatol. 2013;58:522–528.

93. Lou G, Yang Y, Liu F, Ye B, Chen Z, Zheng M, et al. MiR-122 modification enhances the therapeutic efficacy of adipose tissue-derived mesenchymal stem cells against liver fibrosis. J. Cell. Mol. Med. 2017;21:2963–2973.

94. Feng G, Shi H, Li J, Yang Z, Fang R, Ye L, et al. MiR-30e suppresses proliferation of hepatoma cells via targeting prolyl 4-hydroxylase subunit alpha-1 (P4HA1) mRNA. Biochem. Biophys. Res. Commun. 2016;472:516–522.

95. Hu J, Gao DZ. Distinction immune genes of hepatitis-induced heptatocellular carcinoma. Bioinformatics. 2012;28:3191–3194.

96. Wang W, Yuasa T, Tsuchiya N, Ma Z, Maita S, Narita S, et al. The novel tumor-suppressor Mel-18 in prostate cancer: Its functional polymorphism, expression and clinical significance. Int. J. Cancer. 2009;125:2836–2843.

97. Liu X, Wei W, Li X, Shen P, Ju D, Wang Z, et al. BMI1 and MEL18 Promote Colitis-Associated Cancer in Mice via REG3B and STAT3. Gastroenterology. 2017;153:1607–1620.

98. Wang M, Li C, Cui J, Jiao M, Wu T, Jing L, et al. BMI-1, a promising therapeutic target for human

cancer (Review). Oncol. Lett. 2015;10:583–588.

99. Lu YW, Li J, Guo WJ. Expression and clinicopathological significance of Mel-18 and Bmi-1 mRNA in gastric carcinoma. J. Exp. Clin. Cancer Res. 2010;29:143.

100. Chen Q, Yan CQ, Liu FJ, Tong J, Miao SL, Chen JP. Overexpression of the PDCD2-like gene results in Inhibited TNF-α production in activated Daudi cells. Hum. Immunol. 2008;69:259–265.

101. Yang Y, Jin Y, Du W. Programmed cell death 2 functions as a tumor suppressor in osteosarcoma. Int. J. Clin. Exp. Pathol. 2015;8:10894–10900.

102. Matsuhashi S, Hamajima H, Xia JH, Zhang H, Mizuta T, Anzai K, et al. Control of a tumor suppressor PDCD4: Degradation mechanisms of the protein in hepatocellular carcinoma cells. Cell. Signal. 2014;26:603–610.

103. Zhang S, Li J, Jiang Y, Xu Y, Qin C. Programmed cell death 4 (PDCD4) suppresses metastastic potential of human hepatocellular carcinoma cells. J. Exp. Clin. Cancer Res. 2009;28:1–11.

104. Zhang Z, Zha Y, Hu W, Huang Z, Gao Z, Zang Y, et al. The autoregulatory feedback loop of MicroRNA-21/programmed cell death protein 4/Activation protein-1 (MiR-21/PDCD4/AP-1) as a driving force for hepatic fibrosis development. J. Biol. Chem. 2013;288:37082–37093.

105. Damania P, Sen B, Dar SB, Kumar S, Kumari A, Gupta E, et al. Hepatitis B virus induces cell proliferation via HBx-induced microRNA-21 in hepatocellular carcinoma by targeting Programmed Cell Death Protein4 (PDCD4) and Phosphatase and Tensin Homologue (PTEN). PLoS One. 2014;9:2–10.

106. Kamel RR, Amr KS, Afify M, Elhosary YA, Hegazy AE, Fahim HH, et al. Relation between microRNAs and apoptosis in hepatocellular carcinoma. Maced. J. Med. Sci. 2016;4:31–37.

107. Antonacopoulou AG, Grivas PD, Skarlas L, Kalofonos M, Scopa CD, Kalofonos HP. POLR2F, ATP6V0A1 and PRNP expression in colorectal cancer: New molecules with prognostic significance? Anticancer Res. 2008;28:1221–1227.

108. Jia Z, Ai X, Sun F, Zang T, Guan Y, Gao F, et al. Identification of New Hub Genes Associated with Bladder Carcinoma via Bioinformatics Analysis. Tumori J. 2015;101:117–22.

109. Gupta P, Liu B, Wu JQ, Soriano V, Vispo E, Carroll AP, et al. Genome-wide mRNA and miRNA analysis of peripheral blood mononuclear cells (PBMC) reveals different miRNAs regulating HIV/HCV co-infection. Virology. 2014;450–451:336–349.

110. Roset R, Inagaki A, Hohl M, Brenet F, Lafrance-Vanasse J, Lange J, et al. The Rad50 hook domain regulates DNA damage signaling and tumorigenesis. Genes Dev. 2014;28:451–462.

111. El Idrissi M, Hervieu V, Merle P, Mortreux F, Wattel E. Cause-specific telomere factors deregulation in hepatocellular carcinoma. J. Exp. Clin. Cancer Res. 2013;32:1–11.

112. Lin Z, Xu SH, Wang HQ, Cai YJ, Ying L, Song M, et al. Prognostic value of DNA repair based stratification of hepatocellular carcinoma. Sci. Rep. 2016;6:1–13.

113. Machida K, McNamara G, Cheng KT-H, Huang J, Wang C-H, Comai L, et al. Hepatitis C Virus Inhibits DNA Damage Repair through Reactive Oxygen and Nitrogen Species and by Interfering with the ATM-NBS1/Mre11/Rad50 DNA Repair Pathway in Monocytes and Hepatocytes. J. Immunol. 2010;185:6985–6998.

114. Qin Y, Zhou MT, Hu MM, Hu YH, Zhang J, Guo L, et al. RNF26 Temporally Regulates Virus-Triggered Type I Interferon Induction by Two Distinct Mechanisms. PLoS Pathog. 2014;10.

115. Katoh M. Molecular cloning and characterization of RNF26 on human chromosome 11q23 region, encoding a novel ring finger protein with leucine zipper. Biochem. Biophys. Res. Commun. 2001;282:1038–1044.

116. Stefanska B, Cheishvili D, Suderman M, Arakelian A, Huang J, Hallett M, et al. Genome-wide study of hypomethylated and induced genes in patients with liver cancer unravels novel anticancer targets. Clin. Cancer Res. 2014;20:3118–3132.

117. Wong CCL, Wong CM, Tung EKK, Man K, Ng IOL. Rho-kinase 2 is frequently overexpressed in hepatocellular carcinoma and involved in tumor invasion. Hepatology. 2009;49:1583–1594.

118. Huang D, Du X, Yuan R, Chen L, Liu T, Wen C, et al. Rock2 promotes the invasion and metastasis of hepatocellular carcinoma by modifying MMP2 ubiquitination and degradation. Biochem. Biophys. Res. Commun. 2014;453:49–56.

119. Peng F, Jiang J, Yu Y, Tian R, Guo X, Li X, et al. Direct targeting of SUZ12/ROCK2 by miR-200b/c inhibits cholangiocarcinoma tumourigenesis and metastasis. Br. J. Cancer. 2013;109:3092–3104.

120. Liu T, Yu X, Li G, Yuan R, Wang Q, Tang P, et al. Rock2 regulates Cdc25A through ubiquitin proteasome system in hepatocellular carcinoma cells. Exp. Cell Res. 2012;318:1994–2003.

121. Zheng F, Liao YJ, Cai MY, Liu YH, Liu TH, Chen SP, et al. The putative tumour suppressor microRNA-124 modulates hepatocellular carcinoma cell aggressiveness by repressing ROCK2 and EZH2. Gut. 2012;61:278–289.

122. Iizuka M, Murata T, Hori M, Ozaki H. Increased contractility of hepatic stellate cells in cirrhosis is mediated by enhanced $Ca^{2+}$-dependent and $Ca^{2+}$-sensitization pathways. Am. J. Physiol. Gastrointest. Liver Physiol. 2011;300:G1010–G1021.

123. Kavsak P, Rasmussen RK, Causing CG, Bonni S, Zhu H, Thomsen GH, et al. Smad7 Binds to Smurf2 to Form an E3 Ubiquitin Ligase that Targets the TGFbeta Receptor for Degradation. Mol. Cell. 2000;6:1365–1375.

124. Wang J, Zhao J, Chu ESH, Mok MTS, Go MYY, Man K, et al. Inhibitory role of Smad7 in hepatocarcinogenesis in mice and in vitro. J. Pathol. 2013;230:441–452.

125. Feng T, Dzieran J, Gu X, Marhenke S, Vogel A, Machida K, et al. Smad7 regulates compensatory hepatocyte proliferation in damaged mouse liver and positively relates to better clinical outcome in human hepatocellular carcinoma. Clin. Sci. 2015;128:761–774.

126. Liu N, Jiao T, Huang Y, Liu W, Li Z, Ye X. Hepatitis B Virus Regulates Apoptosis and Tumorigenesis through the MicroRNA-15a-Smad7-Transforming Growth Factor Beta Pathway. J. Virol. [Internet]. 2015;89:2739–2749. Available from: http://jvi.asm.org/lookup/doi/10.1128/JVI.02784-14

127. Kan H, Guo W, Huang Y, Liu D. MicroRNA-520g induces epithelial-mesenchymal transition and promotes metastasis of hepatocellular carcinoma by targeting SMAD7. FEBS Lett. 2015;589:102–109.

128. Argentou N, Germanidis G, Hytiroglou P, Apostolou E, Vassiliadis T, Patsiaoura K, et al. TGF-β signaling is activated in patients with chronic HBV infection and repressed by SMAD7 overexpression after successful antiviral treatment. Inflamm. Res. 2016;65:355–365.

129. Li H, Yang X, Yang G, Hong Z, Zhou L, Yin P, et al. Hepatitis C Virus NS5A Hijacks ARFGAP1 To Maintain a Phosphatidylinositol 4-Phosphate-Enriched Microenvironment. J. Virol. 2014;88:5956–5966.

130. Balistreri G, Horvath P, Schweingruber C, Zünd D, McInerney G, Merits A, et al. The host nonsense-mediated mRNA decay pathway restricts mammalian RNA virus replication. Cell Host Microbe. 2014;16:403–411.

131. Ma XN, Liu XY, Yang YF, Xiao FJ, Li QF, Yan J, et al. Regulation of human hepatocellular carcinoma cells by Spred2 and correlative studies on its mechanism. Biochem. Biophys. Res. Commun. 2011;410:803–808.

132. Wakabayashi H, Ito T, Fushimi S, Nakashima Y, Itakura J, Qiuying L, et al. Spred-2 deficiency exacerbates acetaminophen-induced hepatotoxicity in mice. Clin. Immunol. 2012;144:272–282.

133. Momeny M, Khorramizadeh MR, Ghaffari SH, Yousefi M, Yekaninejad MS, Esmaeili R, et al. Effects of silibinin on cell growth and invasive properties of a human hepatocellular carcinoma cell line, HepG-2, through inhibition of extracellular signal-regulated kinase 1/2 phosphorylation. Eur. J. Pharmacol. 2008;591:13–20.

134. Kachroo N, Valencia T, Warren AY, Gnanapragasam VJ. Evidence for downregulation of the negative regulator SPRED2 in clinical prostate cancer. Br. J. Cancer. 2013;108:597–601.

135. Villar V, Kocić J, Santibanez JF. Spred2 inhibits TGF-β1-induced urokinase type plasminogen activator expression, cell motility and epithelial mesenchymal transition. Int. J. Cancer. 2010;127:77–85.

136. Qing T, Yamin Z, Guijie W, Yan J, Zhongyang S. STAT6 silencing induces hepatocellular carcinoma-derived cell apoptosis and growth inhibition by decreasing the RANKL expression. Biomed.

Pharmacother. 2017;92:1–6.

137. Yang W, Lu Y, Xu Y, Xu L, Zheng W, Wu Y, et al. Estrogen represses hepatocellular carcinoma (HCC) Growth via Inhibiting Alternative Activation of Tumor-associated Macrophages (TAMs). J. Biol. Chem. 2012;287:40140–40149.

138. Lim YP, Hsu YA, Tsai KH, Tsai FJ, Peng CY, Liao WL, et al. The impact of polymorphisms in STAT6 on treatment outcome in HCV infected Taiwanese Chinese. BMC Immunol. 2013;14.

139. Jaruga B, Hong F, Sun R, Radaeva S, Gao B. Crucial role of IL-4/STAT6 in T cell-mediated hepatitis: up-regulating eotaxins and IL-5 and recruiting leukocytes. J. Immunol. 2003;171:3233–3244.

140. Gao B, Wang H, Lafdil F, Feng D. STAT proteins - key regulators of anti-viral responses, inflammation, and tumorigenesis in the liver. J. Hepatol. 2012;57:430–441.

141. Greninger AL, Knudsen GM, Betegon M, Burlingame ALB, DeRisi JL. ACBD3 interaction with TBC1 domain 22 protein is differentially affected by enteroviral and kobuviral 3A protein binding. MBio. 2013;4:e00098-13.

142. Ngo HTT, Pham L V., Kim J-W, Lim Y-S, Hwang SB. Modulation of Mitogen-Activated Protein Kinase-Activated Protein Kinase 3 by Hepatitis C Virus Core Protein. J. Virol. 2013;87:5718–5731.

143. Levin A, Neufeldt CJ, Pang D, Wilson K, Loewen-Dobler D, Joyce MA, et al. Functional characterization of nuclear localization and export signals in hepatitis C virus proteins and their role in the membranous web. PLoS One. 2014;9:1–36.

144. Zhao J, Chen J, Lu B, Dong L, Wang H, Bi C, et al. TIP30 induces apoptosis under oxidative stress through stabilization of p53 messenger RNA in human hepatocellular carcinoma. Cancer Res. 2008;68:4133–4141.

145. Lai KKY, Kweon S, Chi F, Hwang E, Kabe Y, Higashiyama R, et al. Stearoyl-CoA Desaturase Promotes Liver Fibrosis and Tumor Development in Mice via Wnt Signaling and Stabilization of Low Density Lipoprotein Receptor-related Proteins 5 and 6. Gastroenterology. 2017;152:1477–1491.

146. Zhu M, Yin F, Fan X, Jing W, Chen R, Liu L, et al. Decreased TIP30 promotes Snail-mediated epithelial-mesenchymal transition and tumor-initiating properties in hepatocellular carcinoma. Oncogene. 2015;34:1420–1431.

147. Tian M, Cheng H, Wang Z, Su N, Liu Z, Sun C, et al. Phosphoproteomic analysis of the highly-metastatic hepatocellular carcinoma cell line, MHCC97-H. Int. J. Mol. Sci. 2015;16:4209–4225.

148. Imai H, Chan EKL, Kiyosawa K, Fu XD, Tan EM. Novel nuclear autoantigen with splicing factor motifs identified with antibody from hepatocellular carcinoma. J. Clin. Invest. 1993;92:2419–2426.

149. Zhang J-Y, Tan EM. Autoantibodies to tumor-associated antigens as diagnostic biomarkers in hepatocellular carcinoma and other solid tumors. Expert Rev. Mol. Diagn. 2010;6:321–328.

150. Kouwaki T, Okamoto T, Ito A, Sugiyama Y, Yamashita K, Suzuki T, et al. Hepatocyte Factor JMJD5 Regulates Hepatitis B Virus Replication. J. Virol. 2016;90:3530–3542.

151. McGarvey TW, Nguyen T, Puthiyaveettil R, Tomaszewski JE, Malkowicz SB. TERE1, a novel gene affecting growth regulation in prostate carcinoma. Prostate. 2003;54:144–55.

152. Fredericks WJ, Sepulveda J, Lai P, Tomaszewski JE, Lin M-F, McGarvey T, et al. The tumor suppressor TERE1 (UBIAD1) prenyltransferase regulates the elevated cholesterol phenotype in castration resistant prostate cancer by controlling a program of ligand dependent SXR target genes. Oncotarget. 2013;4:1075–92.

153. Fredericks WJ, McGarvey T, Wang H, Lal P, Puthiyaveettil R, Tomaszewski J, et al. The Bladder Tumor Suppressor Protein TERE1 (UBIAD1) Modulates Cell Cholesterol: Implications for Tumor Progression. DNA Cell Biol. 2011;30:851–864.

154. Jinghe X, Mizuta T, Ozaki I. Vitamin K and hepatocellular carcinoma: The basic and clinic. World J. Clin. Cases. 2015;3:757.

155. Park ER, Kim SB, Lee JS, Kim YH, Lee DH, Cho EH, et al. The mitochondrial hinge protein, UQCRH, is a novel prognostic factor for hepatocellular carcinoma. Cancer Med. 2017;6:749–760.

156. Woo HG, Park ES, Lee J, Lee Y, Ishikawa T, Kim YJ, et al. Identification of potential driver genes in human liver carcinoma by genome-wide screening. Cancer Res. 2009;69:4059–4066.

157. Sun Y-H, Zhang X-Y, Xie W-Q, Liu G-J, He X-X, Huang Y-L, et al. Identification of UQCRB as an oxymatrine recognizing protein using a T7 phage display screen. J. Ethnopharmacol. 2016;193:133–139.

158. Jung HJ, Kim Y, Chang J, Kang SW, Kim JH, Kwon HJ. Mitochondrial UQCRB regulates VEGFR2 signaling in endothelial cells. J. Mol. Med. 2013;91:1117–1128.

159. Moniz S, Jordan P. Emerging roles for WNK kinases in cancer. Cell. Mol. Life Sci. 2010;67:1265–1276.

160. Van Der Lubbe N, Lim CH, Meima ME, Van Veghel R, Rosenbaek LL, Mutig K, et al. Aldosterone does not require angiotensin II to activate NCC through a WNK4-SPAK-dependent pathway. Pflugers Arch. Eur. J. Physiol. 2012;463:853–863.

161. Mutlu P, Ural AU, Gündüz U. Differential oncogene-related gene expressions in myeloma cells resistant to prednisone and vincristine. Biomed. Pharmacother. 2012;66:506–511.

162. Erkeland SJ, Aarts LH, Irandoust M, Roovers O, Klomp A, Valkhof M, et al. Novel role of WD40 and SOCS box protein-2 in steady-state distribution of granulocyte colony-stimulating factor receptor and G-CSF-controlled proliferation and differentiation signaling. Oncogene. 2007;26:1985–1994.

163. Li XG, Song JD, Wang YQ. Differential expression of a novel colorectal cancer differentiation-related gene in colorectal cancer. World J. Gastroenterol. 2001;7:551–554.

164. Nara H, Onoda T, Rahman M, Araki A, Juliana FM, Tanaka N, et al. Regulation of interleukin-21 receptor expression and its signal transduction by WSB-2. Biochem. Biophys. Res. Commun. 2010;392:171–177.

165. Hu X, Ma S, Huang X, Jiang X, Zhu X, Gao H, et al. Interleukin-21 is upregulated in hepatitis B-related acute-on-chronic liver failure and associated with severity of liver disease. J. Viral Hepat. 2011;18:458–467.

166. Xu Y, Liu AJ, Gao YX, Hu MG, Zhao GD, Zhao ZM, et al. Expression of Ku86 and presence of Ku86 Antibody as biomarkers of hepatitis B virus related hepatocellular carcinoma. Dig. Dis. Sci. 2014;59:614–622.

167. Wei S, Xiong M, Zhan DQ, Liang BY, Wang YY, Gutmann DH, et al. Ku80 functions as a tumor suppressor in hepatocellular carcinoma by inducing s-phase arrest through a p53-dependent pathway. Carcinogenesis. 2012;33:538–547.

168. Li R, Yang Y, An Y, Zhou Y, Liu Y, Yu Q, et al. Genetic polymorphisms in DNA double-strand break repair genes XRCC5, XRCC6 and susceptibility to hepatocellular carcinoma. Carcinogenesis. 2011;32:530–536.

169. Quinn JP, Simpson J, Farina AR. The Ku complex is modulated in response to viral infection and other cellular changes. Biochim. Biophys. Acta. 1992;1131:181–187.

170. Long XD, Zhao D, Wang C, Huang XY, Yao JG, Ma Y, et al. Genetic polymorphisms in DNA repair genes XRCC4 and XRCC5 and aflatoxin B1-related hepatocellular carcinoma. Epidemiology. 2013;24:671–681.

171. Jeffery J, Sinha D, Srihari S, Kalimutho M, Khanna KK. Beyond cytokinesis: The emerging roles of CEP55 in tumorigenesis. Oncogene. 2016;35:683–6390.

172. Liu C, Zhou N, Li J, Kong J, Guan X, Wang X. Eg5 Overexpression Is Predictive of Poor Prognosis in Hepatocellular Carcinoma Patients. Dis. Markers. 2017;2017:2176460.

173. Chen J, Rajasekaran M, Xia H, Zhang X, Kong SN, Sekar K, et al. The microtubule-associated protein PRC1 promotes early recurrence of hepatocellular carcinoma in association with the Wnt/β-catenin signalling pathway. Gut. 2016;65:1522–1534.

174. Doan CC, Doan NT, Nguyen QH, Nguyen MH, Do MS, Le VD. Downregulation of kinesin spindle protein inhibits proliferation, induces apoptosis and increases chemosensitivity in hepatocellular carcinoma cells. Iran. Biomed. J. 2015;19:1–16.

175. Sun HW, Yu XJ, Wu WC, Chen J, Shi M, Zheng L, et al. GLUT1 and ASCT2 as predictors for

prognosis of hepatocellular carcinoma. PLoS One. 2016;11:1–14.

176. Namikawa M, Kakizaki S, Kaira K, Tojima H, Yamazaki Y, Horiguchi N, et al. Expression of amino acid transporters (LAT1, ASCT2 and xCT) as clinical significance in hepatocellular carcinoma. Hepatol. Res. 2015;45:1014–1022.

177. Wu Q, Li J V., Seyfried F, Le Roux CW, Ashrafian H, Athanasiou T, et al. Metabolic phenotype-microRNA data fusion analysis of the systemic consequences of Roux-en-Y gastric bypass surgery. Int. J. Obes. 2015;39:1126–1134.

178. Liu B, Fang M, He Z, Cui D, Jia S, Lin X, et al. Hepatitis B virus stimulates G6PD expression through HBx-mediated Nrf2 activation. Cell Death Dis. 2015;6:1–10.

179. Hu H, Ding X, Yang Y, Zhang H, Li H, Tong S, et al. Changes in glucose-6-phosphate dehydrogenase expression results in altered behavior of HBV-associated liver cancer cells. Am. J. Physiol. Gastrointest. Liver Physiol. [Internet]. 2014;307:G611–G622. Available from: http://www.physiology.org/doi/10.1152/ajpgi.00160.2014

180. Hong X, Song R, Song H, Zheng T, Wang J, Liang Y, et al. PTEN antagonises Tcl1/hnRNPK-mediated G6PD pre-mRNA splicing which contributes to hepatocarcinogenesis. Gut. 2014;63:1635–1647.

181. Rao K, Elm M, Kelly R, Chandar N, Brady E, Rao B, et al. Hepatic hyperplasia and cancer in rats: Metabolic alterations associated with cell growth. Gastroenterology. 1997;113:238–248.

182. Niehof M, Borlak J. EPS15R, TASP1, and PRPF3 Are Novel Disease Candidate Genes Targeted by HNF4α Splice Variants in Hepatocellular Carcinomas. Gastroenterology. 2008;134:1191–1202.

183. Ogihara K, Naya Y, Sato R, Onda K, Ochia H. Analysis of L-type amino acid transporter in canine hepatocellular carcinoma. J. Vet. Med. Sci. 2015;77:527–534.

184. Li J, Qiang J, Chen S-F, Wang X, Fu J, Chen Y. The impact of L-type amino acid transporter 1 (LAT1) in human hepatocellular carcinoma. Tumor Biol. 2013;34:2977–2981.

185. Park Y-Y, Sohn BH, Johnson RL, Kang M-H, Kim SB, Shim J-J, et al. YAP1 and TAZ Activates mTORC1 Pathway by Regulating Amino Acid Transporters in hepatocellular carcinoma. Hepatology. 2016;63:159–172.

186. Tamai S, Masuda H, Ishii Y, Suzuki S, Kanai Y, Endou H. Expression of L-type amino acid transporter 1 in a rat model of liver metastasis: positive correlation with tumor size. Cancer Detect. Prev. 2001;25:439–45.

187. Ohkame H, Masuda H, Ishii Y, Kanai Y. Expression of L-type amino acid transporter I (LAT1) and 4F2 heavy chain (4F2hc) in liver tumor lesions of rat models. J. Surg. Oncol. 2001;78:265–271.

188. Kondoh N, Imazeki N, Arai M, Hada A, Hatsuse K, Matsuo H, et al. Activation of a system A amino acid transporter, ATA1/SLC38A1, in human hepatocellular carcinoma and preneoplastic liver tissues. Int. J. Oncol. 2007;31:81–87.

189. Gkretsi V, Bogdanos DP. Experimental evidence of Migfilin as a new therapeutic target of hepatocellular carcinoma metastasis. Exp. Cell Res. 2015;334:219–227.

190. Liu AM, Xu Z, Shek FH, Wong KF, Lee NP, Poon RT, et al. MiR-122 targets pyruvate kinase M2 and affects metabolism of hepatocellular carcinoma. PLoS One. 2014;9:1–9.

191. Chen Z, Lu X, Wang Z, Jin G, Wang Q, Chen D, et al. Co-expression of PKM2 and TRIM35 predicts survival and recurrence in hepatocellular carcinoma. Oncotarget. 2015;6:2538–48.

192. Liu W-R, Tian M-X, Yang L-X, Lin Y-L, Jin L, Ding Z-B, et al. PKM2 promotes metastasis by recruiting myeloid-derived suppressor cells and indicates poor prognosis for hepatocellular carcinoma. Oncotarget. 2015;6.

193. Dong T, Yan Y, Chai H, Chen S, Xiong X, Sun D, et al. Pyruvate kinase M2 affects liver cancer cell behavior through up-regulation of HIF-1α and Bcl-xL in culture. Biomed. Pharmacother. 2015;69:277–284.

194. Moylan CA, Pang H, Dellinger A, Suzuki A, Garrett ME, Guy CD, et al. Hepatic gene expression profiles differentiate presymptomatic patients with mild versus severe nonalcoholic fatty liver disease.

Hepatology. 2014;59:471–482.

195. Wang HW, Hsieh TH, Huang SY, Chau GY, Tung CY, Su CW, et al. Forfeited hepatogenesis program and increased embryonic stem cell traits in young hepatocellular carcinoma (HCC) comparing to elderly HCC. BMC Genomics. 2013;14:1.

196. Hirasawa Y, Arai M, Imazeki F, Tada M, Mikata R, Fukai K, et al. Methylation status of genes upregulated by demethylating agent 5-aza-2′-deoxycytidine in hepatocellular carcinoma. Oncology. 2007;71:77–85.

197. Chen F, Zhu HH, Zhou LF, Wu SS, Wang J, Chen Z. IQGAP1 is overexpressed in hepatocellular carcinoma and promotes cell proliferation by Akt activation. Exp. Mol. Med. 2010;42:477–483.

198. Xia F-D, Wang Z-L, Chen H-X, Huang Y, Li J-D, Wang Z-M, et al. Differential expression of IQGAP1/2 in Hepatocellular carcinoma and its relationship with clinical outcomes. Asian Pacific J. Cancer Prev. 2014;15:4951–6.

199. Zhang S, Lin B, Li B. Evaluation of the diagnostic value of alpha- L -fucosidase , alpha-fetoprotein and thymidine kinase 1 with ROC and logistic regression for hepatocellular carcinoma. FEBS Open Bio. 2015;5:240–4.

200. Sangro B, Mazzolini G, Ruiz M, Ruiz J, Quiroga J, Herrero I, et al. A phase i clinical trial of thymidine kinase-based gene therapy in advanced hepatocellular carcinoma. Cancer Gene Ther. 2010;17:837–843.

201. Sun Y, Wang Y, Yin Y, Chen X, Sun Z. GSTM3 reverses the resistance of hepatoma cells to radiation by regulating the expression of cell cycle/apoptosis-related molecules. Oncol. Lett. 2014;8:1435–1440.

202. White DL, Li D, Nurgalieva Z, El-Serag HB. Genetic variants of glutathione S-transferase as possible risk factors for hepatocellular carcinoma: A HuGE systematic review and meta-analysis. Am. J. Epidemiol. 2008;167:377–389.

203. Brind AM, Hurlstone A, Edrisinghe D, Gilmore I, Fisher N, Pirmohamed M, et al. The role of polymorphisms of glutathione S-transferases GSTM1, M3, P1, T1 and A1 in susceptibility to alcoholic liver disease. Alcohol Alcohol. 2004;39:478–483.

204. Inoue M, Takahashi Y, Fujii T, Kitagawa M, Fukusato T. Significance of downregulation of liver fatty acid-binding protein in hepatocellular carcinoma. World J. Gastroenterol. 2014;20:17541–17551.

205. Wang B, Tao X, Huang C-Z, Liu J-F, Ye Y-B, Huang A-M. Decreased expression of liver-type fatty acid-binding protein is associated with poor prognosis in hepatocellular carcinoma. Hepatogastroenterology. 2014;61:1321–6.

206. Zhang J, Sun M, Li R, Liu S, Mao J, Huang Y, et al. Ech1 is a potent suppressor of lymphatic metastasis in hepatocarcinoma. Biomed. Pharmacother. 2013;67:557–560.

207. Zhang S, Wang XM, Yin ZY, Zhao WX, Zhou JY, Zhao BX, et al. Chloride intracellular channel 1 is overexpression in hepatic tumor and correlates with a poor prognosis. Acta Pathol. Microbiol. Immunol. Scand. 2013;121:1047–1053.

208. Megger DA, Bracht T, Kohl M, Ahrens M, Naboulsi W, Weber F, et al. Proteomic Differences Between Hepatocellular Carcinoma and Nontumorous Liver Tissue Investigated by a Combined Gel-based and Label-free Quantitative Proteomics Study. Mol. Cell. Proteomics. 2013;12:2006–2020.

209. Li RK, Zhang J, Zhang YH, Li ML, Wang M, Tang JW. Chloride intracellular channel 1 is an important factor in the lymphatic metastasis of hepatocarcinoma. Biomed. Pharmacother. 2012;66:167–172.

210. Zhang J, Li M, Song M, Chen W, Mao J, Song L, et al. Clic1 plays a role in mouse hepatocarcinoma via modulating Annexin A7 and Gelsolin in vitro and in vivo. Biomed. Pharmacother. 2015;69:416–419.

211. Wei X, Li J, Xie H, Wang H, Wang J, Zhang X, et al. Chloride intracellular channel 1 participates in migration and invasion of hepatocellular carcinoma by targeting maspin. J. Gastroenterol. Hepatol. 2015;30:208–216.

212. Baba H, Teramoto K, Kawamura T, Mori A, Imamura M, Arii S. Dihydropyrimidine dehydrogenase and thymidylate synthase activities in hepatocellular carcinomas and in diseased livers. Cancer Chemother. Pharmacol. 2003;52:469–476.

213. Sunaga M, Tomonaga T, Yoshikawa M, Ebara M, Shimada H, Saisho H, et al. Gene Expression of 5-Fluorouracil Metabolic Enzymes in Hepatocellular Carcinoma and Non-Tumor Tissue. J. Chemother. 2007;19:709–15.

214. Yu MC, Yuan JM, Lu SC. Alcohol, cofactors and the genetics of hepatocellular carcinoma. J. Gastroenterol. Hepatol. 2008;23:S92-7.

215. Libra M, Navolanic PM, Talamini R, Cecchin E, Sartor F, Tumolo S, et al. Thymidylate synthetase mRNA levels are increased in liver metastases of colorectal cancer patients resistant to fluoropyrimidine-based chemotherapy. BMC Cancer. 2004;4:11.

216. Nii A, Shimada M, Ikegami T, Harino Y, Imura S, Morine Y, et al. Significance of dihydropyrimidine dehydrogenase and thymidylate synthase mRNA expressions in hepatocellular carcinoma. Hepatol. Res. 2009;39:274–281.

217. Krupa R, Czarny P, Wigner P, Wozny J, Jablkowski M, Kordek R, et al. The Relationship Between Single-Nucleotide Polymorphisms, the Expression of DNA Damage Response Genes, and Hepatocellular Carcinoma in a Polish Population. DNA Cell Biol. 2017;36:693–708.

218. Wang SC, Yang JF, Wang CL, Huang CF, Lin YY, Chen YY, et al. Distinct subpopulations of hepatitis C virus infectious cells with different levels of intracellular hepatitis C virus core protein. Kaohsiung J. Med. Sci. 2016;32:487–493.

219. Li K, Ding S, Chen K, Qin D, Qu J, Wang S, et al. Hepatitis B virus X protein up-regulates AKR1C1 expression through nuclear factor-Y in human hepatocarcinoma cells. Hepat. Mon. 2013;13:1–9.

220. Wang L, Huang J, Jiang M, Chen Q, Jiang Z, Feng H. CAMK1 phosphoinositide signal-mediated protein sorting and transport network in human hepatocellular carcinoma (HCC) by biocomputation. Cell Biochem. Biophys. 2014;70:1011–1016.

221. Besharat S, Katoonizadeh A, Moossavi S, Darvishi Z, Roshandel, Gholamreza Poustchi H, Mohamadkhani A. The Possible Impact of Sortilin in Reducing HBsAg Expression in Chronic Hepatitis B. J. Med. Virol. 2016;88:647–652.

222. Li C, Wang Y, Wang S, Wu B, Hao J, Fan H, et al. Hepatitis B Virus mRNA-Mediated miR-122 Inhibition Upregulates PTTG1-Binding Protein, Which Promotes Hepatocellular Carcinoma Tumor Growth and Cell Invasion. J. Virol. 2013;87:2193–2205.

223. Thakral S, Ghoshal K. miR-122 is a Unique Molecule with Great Potential in Diagnosis, Prognosis of Liver Disease, and Therapy Both as miRNA Mimic and Antimir. Curr. Gene Ther. 2015;15:142–150.

224. Karimi-Googheri M, Daneshvar H, Nosratabadi R, Zare-Bidaki M, Hassanshahi G, Ebrahim M, et al. Important Roles Played by TGF- b in Hepatitis B Infection. 2014;86:102–108.

225. Yu X, Guo R, Ming D, Deng Y, Su M, Lin C, et al. The transforming growth factor β1/interleukin-31 pathway is upregulated in patients with hepatitis B virus-related acute-on-chronic liver failure and is associated with disease severity and survival. Clin. Vaccine Immunol. 2015;22:484–492.

226. Yang P, Markowitz GJ, Wang X-F. The hepatitis B virus-associated tumor microenvironment in hepatocellular carcinoma. Natl. Sci. Rev. 2014;1:396–412.

227. Gori M, Arciello M, Balsano C. MicroRNAs in Nonalcoholic Fatty Liver Disease : Novel Biomarkers and Prognostic Tools during the Transition from Steatosis to Hepatocarcinoma. Biomed Res. Int. 2014;2014:741465.

228. Dooley S, ten Dijke P. TGF- β in progression of liver disease. Cell Tissue Res. 2012;347:245–256.

229. Chen W-X, Li Y-M, Yu C-H, Cai W-M, Zheng M, Chen F. Quantitative analysis of transforming growth factor beta 1 mRNA in patients with alcoholic liver disease. World J. Gastroenterol. 2002;8:379–381.

230. Yang L, Roh YS, Song J, Zhang B, Liu C, Loomba R, et al. TGF-β Signaling in Hepatocytes Participates in Steatohepatitis Through Regulation of Cell Death and Lipid Metabolism. Hepatology. 2014;59:483–495.

231. Halász T, Horváth G, Pár G, Werling K, Kiss A, Schaff Z, et al. miR-122 negatively correlates with liver fibrosis as detected by histology and fibroscan. World J. Gastroenterol. 2015;21:7814–7823.

232. Pohlers D, Brenmoehl J, Löffler I, Müller CK, Leipner C, Schultze-Mosgau S, et al. TGF-β and fibrosis in different organs — molecular pathway imprints. Biochim. Biophys. Acta. 2009;1792:746–756.

233. Bandiera S, Pfeffer S, Baumert TF, Zeisel MB. Review miR-122 – A key factor and therapeutic target in liver disease. J. Hepatol. 2015;62:448–457.

234. Giannelli G, Villa E, Lahn M. Transforming Growth Factor-β as a Therapeutic Target in Hepatocellular Carcinoma. Cancer Res. 2014;74:1890–1894.

235. Padgett KA, Lan RY, Leung PC, Lleo A, Dawson K, Pfeiff J, et al. Primary Biliary Cirrhosis is Associated With Altered Hepatic microRNA Expression. J. Autoimmun. 2009;32:246–253.

236. Neuman M, Angulo P, Malkiewicz I, Jorgensen R, Shear N, Dickson ER, et al. Tumor necrosis factor-α and transforming growth factor-β reflect severity of liver damage in primary biliary cirrhosis. J. Gastroenterol. Hepatol. 2002;17:196–202.

237. Eden E, Navon R, Steinfeld I, Lipson D, Yakhini Z. GOrilla: A tool for discovery and visualization of enriched GO terms in ranked gene lists. BMC Bioinformatics. 2009;10:1–7.

238. Miranda KC, Huynh T, Tay Y, Ang YS, Tam WL, Thomson AM, et al. A Pattern-Based Method for the Identification of MicroRNA Binding Sites and Their Corresponding Heteroduplexes. Cell. 2006;126:1203–1217.

239. Yoo YD, Ueda H, Park K, Flanders KC, Lee YI, Jay G, et al. Regulation of transforming growth factor-beta 1 expression by the hepatitis B virus (HBV) X transactivator. Role in HBV pathogenesis. J. Clin. Invest. 1996;97:388–395.

240. Chen Y, Shen A, Rider PJ, Yu Y, Wu K, Mu Y, et al. A liver-specific microRNA binds to a highly conserved RNA sequence of hepatitis B virus and negatively regulates viral gene expression and replication. FASEB J. 2011;12:4511–4521.
